# The gut microbiota-immune-brain axis in a wild vertebrate: dynamic interactions and health impacts

**DOI:** 10.1101/2024.08.01.605092

**Authors:** Hugo Pereira, Joseph I. Hoffamn, Oliver Krüger, Gábor Á. Czirják, Tony Rinaud, Meinolf Ottensmann, Kai-Philipp Gladow, Barbara A. Caspers, Öncü Maraci, Sylvia Kaiser, Nayden Chakarov

## Abstract

The gut microbiota-immune-brain axis is a feedback network which influences diverse physiological processes and plays a pivotal role in overall health and well-being. Although research in humans and laboratory mice has shed light into the associations and mechanisms governing this communication network, evidence of such interactions in wild, especially in young animals, is lacking. We therefore investigated these interactions during early development in a population of common buzzards (*Buteo buteo*) and their effects on individual condition. In a longitudinal study, we used a multi-marker approach to establish potential links between the bacterial and eukaryotic gut microbiota, a panel of immune assays and feather corticosterone measurements as a proxy for long-term stress. Using Bayesian structural equation modelling, we found no support for feedback between gut microbial diversity and immune or stress parameters. However, we did find strong relationships in the feedback network. Immunity was negatively correlated with corticosterone levels, and microbial diversity was positively associated with nestling body condition. Furthermore, corticosterone levels and eukaryotic microbiota diversity decreased with age while immune activity increased. The absence of conclusive support for the microbiota-immune-brain axis in common buzzard nestlings, coupled with the evidence for stress mediated immunosuppression, suggests a dominating role of stress-dominated maturation of the immune system during early development. Confounding factors inherent to wild systems and developing animals might override associations known from adult laboratory model subjects. The positive association between microbial diversity and body condition indicates the potential health benefits of possessing a diverse and stable microbiota.

## 1 INTRODUCTION

A growing body of evidence highlights a crucial relationship among gut microbes, the environment and host immunity, which also moulds the maturation and function of the central nervous system (CNS) (Fung, 2020; Chin et al., 2020). This intricate feedback system encompasses the brain, the autonomic and enteric nervous systems, the endocrine and immune systems, and the gut microbiome (Martin et al., 2018). Gut-brain communication occurs through various mechanisms, including neuroanatomical pathways and the neuroendocrine axis (hypothalamic-pituitary-adrenal or HPA axis). However, increasing importance has been given to the interactions between the microbiota and the immune system as a fundamental pathway regulating microbiota-gut-brain communication (Fachi et al., 2019). For example, the products resulting from the metabolic activities of the gut microbiota include bioactive peptides such as neurotransmitters, short-chain fatty acids, intestinal hormones, and branched-chain amino acids that play pivotal roles in regulating communication between the gut, brain and the immune system (Sperringer et al., 2017; Cryan et al., 2019; Ding et al., 2020; Krishnamurthy et al., 2023). These metabolites are able to enter the circulatory system to communicate with the brain, eliciting stimulation of the HPA axis (Lynch et al., 2024; Sudo et al., 2004). Additionally, they exert a direct influence on the mucosal immune system, where specific metabolites can function as immune signalling molecules, activating or enhancing systemic immune responses (Ding et al., 2020; Long-Smith et al., 2020). In turn, the brain regulates the gastrointestinal tract and overall organismal homeostasis (Mindus et al., 2021).

Various factors, including, predation (Mohring et al., 2023), heat stress (Lin et al., 2006), high energetic demands during reproduction (Fletcher et al., 2015), and food availability (Romero and Wikelski, 2001) are known to induce elevated stress levels in animals. Corticosterone, a key glucocorticoid (GC) in various vertebrate species, is recognised as a classic endocrine response to stress but also for its role in energy regulation (Almasi et al., 2009). GCs mediate ongoing stress responses, either via maintaining basal levels, allowing other aspects of the stress response to act efficiently, or by actively triggering the stress response (Sapolsky et al., 2000). An alternative view suggests that GCs may suppress the stress response, preventing detrimental over-activation (Sapolsky et al., 2000). Research shows the importance of glucocorticoids in gut-brain communication, with increased GC levels affecting microbial diversity and composition (Petrullo et al., 2022). The gut microbiota also contributes to the development of the host immune system during early life stages, regulates it to maintain gut homeostasis and protects the host from the colonisation of potential pathogens (Lo et al., 2021; Berman et al., 2023).

Increasing importance has been given to the neonatal microbiome and its developmental trajectory during early life which affects host immunity and neural activity maturation (Jašarević and Bale, 2019; Ratsika et al., 2023). For example, in humans, the mode of delivery significantly influences the normal colonisation of the gut microbiota (Ratsika et al., 2023). C-section deliveries have been linked to immune disturbances (allergies and asthma) (Roduit et al., 2009; Bisgaard et al., 2011) and disruptions in the structure and function of the CNS, with indications of increased neuronal cell death (Castillo-Ruiz et al., 2018) and deficits in early communication skills and social behaviour later in life (Morais et al., 2020). In murine models, penicillin treatment during late pregnancy and early postnatal life has been shown to induce alterations in the gut microbiota, leading to heightened cytokine expression, altered integrity of the blood–brain barrier, and significant differences in behaviour (Ratsika et al., 2023). Interactions among these functional systems, particularly during sensitive developmental periods, carry consequences later in life, influencing behaviour, disease susceptibility, general health and survival (Strange et al., 2016; Francella et al., 2022; Ratsika et al., 2023).

Substantial evidence emphasises the importance of the microbiota-immune-brain axis in humans and laboratory animals (Fung, 2020). Yet, the understanding of these links in wild animals remains limited (Hird, 2017; Davidson et al., 2020). Studies in birds associated exploratory behaviour with microbiota diversity, while learning and memory performance have been correlated with compositional differences (Florkowski and Yorzinski, 2023; Slevin et al., 2020). The gut microbiota’s links to stress have also been explored: in common toad tadpoles (*Bufo bufo*), elevated baseline corticosterone associates with higher microbial diversity (Gabor et al., 2022), while American red squirrels (*Tamiasciurus hudsonicus*) show lower alpha diversity and fewer gastrointestinal pathogens in response to elevated glucocorticoids (Petrullo et al., 2022). Studies of the microbiota-immune axis in wild barn swallows (*Hirundo rustica*) and Egyptian fruit bats (*Rousettus aegyptiacus*) demonstrate that antigen challenges (phytohaemagglutinin and lipopolysaccharides, respectively) can induce changes in gut microbiota composition, which in turn predict the strength of the immune response (Kreisinger et al., 2018; Berman et al., 2023). However, since the majority of these studies have involved adult individuals, it is uncertain whether similar patterns would be observed in nestlings and juveniles. In a rare exception, Stoffel et al. (2020) found that a small proportion of the variation in beta diversity among northern elephant seal juveniles was explained by health status (assessed by counting various white blood cell populations) yet a clear pattern emerged where healthier individuals exhibited higher microbiota diversity.

Here, we investigated the gut microbiota-immune-brain axis in a wild vertebrate population, drawing on predictions based on human and murine models. Using a wild population of common buzzards (*Buteo buteo*), we sampled nestlings and collected information on bacterial and eukaryotic microbiota diversity (microbiota component). Long-term stress was measured from feather corticosterone (brain component). We performed a series of immune assays (immune component) and estimated the body condition of each individual. Furthermore, we assessed each component at two distinct time points throughout nestling development to incorporate developmental trajectories and dynamic changes. Our approach diverges from traditional methodologies, which often rely on single-point estimates. All components were subsequently integrated into a structural equation modelling framework (Grace, 2006; Lefcheck, 2016) incorporating the following assumptions: **1.** All components of the axis exert an influence on the body condition of the nestlings; **2.** The immune system is influenced by both stress levels and the microbiota; **3.** Corticosterone levels affect microbiota diversity. We previously showed that as individuals mature, microbiota diversity declines suggesting a shift towards a stable community following uncontrolled colonisation after hatching (Pereira et al., 2024); therefore, we propose that as individuals age, their gut microbiota diversity will decline, their immune system will mature (increased immune capacity), and stress levels will decrease, leading to improved body condition.

## 2 METHODS

We used existing microbiome data from Pereira et al. (2024) and complemented these with new data on feather corticosterone and a set of immune markers. We successfully obtained all sample types from a set of 43 common buzzard individuals, each sampled at two different time points during their nestling phase, totalling 86 samples (Appendix A Table S1-S3). Individuals were sampled across two different habitats: North of the Teutoburg Forest and south of the Teutoburg Forest (8° 25’E and 52° 6’N; Eastern Westphalia, Germany), as described in Krüger (2004) and Jonker et al. (2014).

### 2.1 Sample collection

In brief, body weight and wing length were recorded at each sampling event, with a nine-day interval on average. The first sampling occurred on average at 19 days of age (mean ± s.d. = 19.3 ± 5.29 days) and the second at 28 days (mean ± s.d. = 27.8 ± 5.16 days). Due to difficulties in precise age estimation prior to sampling (the hatching date was estimated by the number of droppings on the ground below the nest; age was determined following the initial visit to the nest and subsequent measurement of wing length), it was impossible to sample individuals at exactly the same age at sampling points 1 and 2. Nestling age was calculated (post-first sampling) using a sex-specific polynomial regression model on wing length (Bijlmsa, 1999). Body condition index (BCI) was determined by extracting the residuals of a logarithmic regression of weight on wing length, adjusting for sex. Blood samples (500 µl) were collected from the ulnar vein and stored in heparinized tubes. A small blood drop was used for smears, a portion was stored in ethanol for sex determination, and the remaining volume was centrifuged in the field until separation of plasma was visible. Separated plasma was transferred to a new tube, stored in dry ice, and subsequently stored at −80 °C until further analysis. The remaining red blood cells were resuspended in PBS solution and stored at −20 °C. Cloacal swabs for gut microbiota analysis were obtained and stored in RNAlater, first in dry ice and then long term at −80 °C. In order to assess corticosterone levels, one interscapular feather was pulled from each bird and these were individually stored in paper envelopes.

### 2.2 Microbiome profiling: DNA isolation, sequencing, and data processing

For detailed procedures on microbiome sequencing data processing see Pereira et al. (2024).

#### 2.2.1 DNA isolation and sequencing

Cloacal swabs underwent DNA extraction using a modified phenol-chloroform protocol. For gene library preparation, the “Illumina 16S Metagenomic Library Preparation Guide” (15044223 Rev.B) was followed. A multimarker approach was used, targeting the V4 region of the 16S ribosomal RNA (rRNA) with the primers 515F (Parada) (Parada et al., 2016) and 806R (Apprill) (Apprill et al., 2015). To be able to capture not only bacteria but also eukaryotes, the D8-D9 region of the 28S rRNA was targeted with the primers GA20F[5]/RM9Rb (Machida and Knowlton, 2012). PCRs were conducted in 25 µl reaction volumes containing 5 µl DNA, 12.5 µl KAPA HiFi HotStart ReadyMix, 1 µl of each primer (1 µM), and 6 µl of PCR-grade water. Index-PCRs utilised Illumina Nextera XT V2 index kits. Libraries were equimolarly pooled, and sequenced on an Illumina MiSeq platform (0.4% MiSeq run) with a 2×300 bp paired-end reads protocol.

#### 2.2.2 16S rRNA sequence data processing

Sequence data were imported into QIIME2 (Quantitative Insights Into Microbial Ecology 2, version 2022.11) (Bolyen et al., 2019). Quality assessment was done by visualising quality plots and Amplicon Sequencing Variants (ASVs) were inferred using the Divisive Amplicon Denoising Algorithm pipeline (DADA2) (Callahan et al., 2016). Taxonomy was assigned using a SILVA 138.1-trained naive Bayes taxonomic classifier (Quast et al., 2012). Contaminants were identified and removed with the decontam package version 1.18 (Davis et al., 2018). ASVs assigned to Mitochondria, Chloroplast, Vertebrata, Eukaryota, and unassigned taxa were filtered out using QIIME2. Singletons and samples with a minimum frequency below 500 reads were removed. The remaining ASV’s were aligned with MAFFT (Katoh et al., 2002) and used to construct a phylogeny with FastTree (Price et al., 2010), both implemented in QIIME2.

#### 2.2.3 28S rRNA sequence data processing

Demultiplexed Illumina sequence data were imported into R version 4.2.2 (R Core Team, 2022). Locus-specific primers were removed using Cutadapt (version 4.4) (Martin et al., 2018). In QIIME2, quality assessment was performed through visualisation of quality plots. ASV inference was conducted using DADA2 following the methodology outlined by Callahan et al. (2016). Sequences were trimmed to eliminate low quality regions and paired-end reads were concatenated. In QIIME2, taxonomy was assigned using the naive Bayes taxonomic classifier trained on the SILVA 138.1 database. The decontam pipeline in R was applied to remove contaminants and to perform taxonomy-based filtering (host reads, Mitochondria, Chloroplast, Vertebra and unassigned reads were removed), removal of unique features, and filtering of samples with fewer than 500 reads in QIIME2. The resulting ASV’s were aligned using MAFFT, and a phylogeny was constructed using FastTree.

### 2.3 Assessment of immunity

Due to the complexity of the immune system, we measured four innate immune parameters (bacterial killing activity against Escherichia coli (BKA), lysozyme and haptoglobin concentrations, natural antibodies (HA) and complement (HL) titres) and one component of the acquired immune system (total immunoglobulin Y (IgY) concentration). All these assays are regularly used in wild bird species, including raptor nestlings, both in comparative and within-species studies (see references below).

#### 2.3.1 Bacterial Killing Activity

The bacterial killing activity (BKA) against *Escherichia coli* (ATCC No 8739) was used to characterise the functional activity of a bird’s constitutive innate immune system (Irene Tieleman et al., 2005) using a spectrophotometric version of the assay (Nebel et al., 2021; Brust et al., 2022; Vincze et al., 2022). This assay measures the plasma’s capacity to kill microbes *ex vivo*, determining the organism’s ability to eliminate bacterial pathogens encountered, providing an environmentally relevant immune response (Millet et al., 2007; Tieleman, 2018). The assay evaluates the synergic function of several immune components (humoral ones in case of working with plasma), including antibacterial enzymes, complement components, and natural antibodies (French and Neuman-Lee, 2012). Briefly, 12 µl of 1:7 PBS-diluted sample was pipetted in duplicate into 96-well-plate and mixed with 4 µl of *≈*1.5 × 105 colony-forming units (CFU)/ml. A positive (not containing any plasma) and a negative control (not containing any *E. coli* or plasma) was run on each plate. After incubation for 30 min at 37°C, 83µl of tryptic soy broth (#22092, Fluka) was added to each well. Absorbance at 300nm was measured with a spectrophotometer (Biotek; µQuant Microplate Spectrophotometer) to determine background absorbance and again after the plates had been incubated for 12 h at 37°C. The BKA was quantified as the bacteria growth in plasma after 12 h (in %) subtracted by the background absorption in relation to the positive control (Brust et al., 2022).

#### 2.3.2 Lysozyme

Lysozyme is an antibacterial enzyme that causes rapid cell lysis, especially in Gram-positive bacteria. It is part of the constitutive innate immune system and is often measured to assess inflammation-induced levels in plasma (Millet et al., 2007). To measure its concentration in plasma, we used the lysoplate assay Prüter et al. (2020); Brust et al. (2022): 10 µl of sample was inoculated in the test holes of a 1% Noble agar gel (A5431, Sigma) containing 50 mg/100 ml lyophilized *Micrococcus lysodeikticus* (M3770, Sigma), a bacterium which is particularly sensitive to lysozyme concentration. Crystalline hen egg white lysozyme (L6876, Sigma) (concentration: 0.5, 0.8, 1, 2, 4, 8, 10, 20 and 40 µg/ml) was used to prepare a standard curve for each plate. After 20h incubation at 37°C, a clear zone developed in the area of the gel surrounding the sample inoculation site corresponding to the bacterial lysis. The diameters of the cleared zones are proportional to the log of the lysozyme concentration. This area was measured three times digitally using the software ImageJ (version 1.48, ImageJ) and the mean was converted to a semi-logarithmic plot into hen egg lysozyme equivalents (HEL equivalents, expressed in µg/mL) according to the standard curve Prüter et al. (2020); Brust et al. (2022).

#### 2.3.3 Haptoglobin

Haptoglobin, is an acute-phase protein in birds and is part of the induced innate immune system. Acute phase proteins are key indicators of immunological function. Their concentrations fluctuate over time, reflecting changes in health and physiological condition (Hõrak et al., 2002, 2003). Levels can rise quickly in response to infection, inflammation, or trauma (Millet et al., 2007). Elevated plasma haptoglobin often signifies the onset of a non-specific immune response (Matson et al., 2012). We measured haptoglobin concentrations with a commercial kit (TP801, Tri-Delta Diagnostics, Inc.) following the instructions of the manufacturer. Haptoglobin concentrations (mg/ml) in undiluted plasma samples were calculated according to the standard curve on each plate (Prüter et al., 2020; Nebel et al., 2021).

#### 2.3.4 Haemolysis–haemagglutination assay

The levels of the natural antibodies and complement were assessed by using a haemolysis-haemagglutination assay as described by (Matson et al., 2005; Nebel et al., 2021; Brust et al., 2022). Natural antibodies, quantified as Haemagglutination (HA) titers, bind non-specifically to various antigens and play crucial roles in opsonization (Ochsenbein et al., 1999; Forthal, 2014). Haemolysis (HL), facilitated by the complement system, is a component of the innate immune system, its activation leads to cell lysis, particularly of cells that have been opsonized (Nauta et al., 2004). After pipetting 10 µl of plasma into the first two columns of a U-shaped 96-well microtiter plate, 10 µl sterile PBS was added to columns 2–12. The content of the second column wells was serially diluted (1:2) until the 11*^th^* column, resulting in a dilution series for each sample from 1/1 to 1/1024. The last column of the plate was used as negative control, containing PBS only. 10 µl of 1% rabbit red blood cells (supplied by Innovative Research) suspension was added to all wells and incubated at 37°C for 90 min. After incubation, in order to increase the visualisation of agglutination, the plates were tilted at a 45° angle at room temperature. Agglutination and lysis, which reflect the activity of the natural antibodies and the interaction between these antibodies and complement (Matson et al., 2005; Prüter et al., 2020), was recorded after 20 and 90 min, respectively. Haemagglutination is characterised by the appearance of clumped red blood cells as a result of antibodies binding multiple antigens, while during haemolysis, the red blood cells are destroyed. Titres of the natural antibodies and complement were given as the log2 of the reciprocal of the highest dilution of serum showing positive haemagglutination or lysis, respectively (Matson et al., 2005; Prüter et al., 2020; Brust et al., 2022).

#### 2.3.5 Total immunoglobulin Y (IgY) concentration

Total IgY, the avian equivalent to mammalian immunoglobulin G, is the primary humoral effector of the adaptive immune system, playing a critical role in neutralising pathogens (Warr et al., 1995). IgY is essential for long-term immunity, providing protection against recurrent infections. Total IgY was measured using an ELISA with commercial anti-chicken antibodies (Bourgeon et al., 2010; Hanssen et al., 2013). Ninety-six well high-binding ELISA plates (82.1581.200, Sarstedt) were coated with 100 µl of diluted plasma sample (1:4000 diluted in carbonate–bicarbonate buffer, in duplicates) and incubated first for 1 h at 37°C and then overnight at 4°C. After incubation, the plates were washed with a 200 µl solution of PBS and PBS–Tween, before 100 µl of a solution of 1% gelatine in PBS–Tween was added. Plates were then incubated at 37°C for 1 h, washed with PBS–Tween and 100 µl of polyclonal rabbit anti-chicken IgY conjugated with peroxidase (A-9046, Sigma) at 1:250 (v/v) was added. Following 2 h incubation at 37°C, the plates were washed again with PBS–Tween three times. After washing, 100 µl of revealing solution [peroxide diluted 1:1000 in ABTS (2,20-azino-bis-(3-ethylbenzthiazoline-6-sulphonic acid))] was added, and the plates were incubated for 1 h at 37°C. The final absorbance was measured at 405 nm using a photometric microplate reader (µQuant Microplate Spectrophotometer, Biotek) and subsequently defined as total serum IgY levels (Brust et al., 2022).

### 2.4 Feather-corticosterone (f-CORT) determination

The entire feather was weighed and placed in a tube. For every 10 mg of feather, 1 ml of 100% methanol (p.a.) was added, followed by crushing with scissors. Subsequently, the samples were subjected to an ultrasonic bath incubation at 30°C for 30 minutes, followed by overnight incubation in an overhead shaker (25 rpm). On the following day, centrifugation (2 min, 19800 x g) was performed, and the supernatant transferred to a new tube. The feather pellet underwent two washes, each with half of the extraction volume (2 min each at 19800 x g); the resulting supernatants were then combined and one further centrifugation was performed for 10 min at 19800 x g. The supernatant was then filtered through a PTFE membrane (Merck Millipore: Ultrafree-CL), and the filtrate was temporarily stored at −20°C. A defined amount of methanol (500 µl) from the temporarily stored filtrate was evaporated in a vacuum concentrator. The samples were re-suspended in 250 µl ELISA buffer (Cayman Chemicals Inc. [# 501320]) and stored at −20°C for at least overnight before conducting the ELISA. Corticosterone concentrations were determined in triplicate following the manufacturer’s instructions. f-CORT assay validation is presented in Appendix F.

### 2.5 Statistical analysis

In order to evaluate sequencing depth and sample coverage, rarefaction curves were constructed in QIIME2. Rarefaction was then applied to the 16S rRNA dataset at 4000 reads and to the 28S rRNA dataset at 2000 reads. Utilising the q2-diversity alpha plugin, three alpha diversity metrics were calculated: Shannon diversity index (Shannon, 1948); Faith’s Phylogenetic Diversity (Faith PD) (Faith and Baker, 2006) and number of observed ASV’s. Prior to constructing the Structural Equation Model (SEM), we explored associations of the variables of interest (BCI, f-CORT, immune assays) with host and environmental factors examined by Pereira et al. (2024). These included sex, habitat and rank (dominance hierarchy within the brood). We have previously established that none of these variables affects microbiota alpha diversity (Pereira et al., 2024). This preliminary analysis aimed to assess the relevance of these variables for the construction of the SEM. For each variable of interest, individual linear mixed models (LMMs) with a Gaussian distribution were constructed using the lmer function from the lme4 package in R (Bates et al., 2015). The significance of factors was assessed through analysis of variance (ANOVA). To account for repeated samples and nest sharing, Nest ID and Individual ID were incorporated as nested random effects (Individual ID nested within Nest ID). Additionally, the Benjamini-Hochberg method was applied to correct the p-values for multiple hypothesis testing (Benjamini and Hochberg, 1995). Scripts and results are presented in Appendix B.

#### 2.5.1 Structural Equation Modelling Approach (SEM)

In order to tackle the complex interactions within the gut microbiota-immune-brain axis and their impact on individual condition, we employed a SEM strategy. SEM is a statistical methodology that allows for the simultaneous testing of complex relationships among multiple variables. Integrating factor analysis and multivariate regression analysis, SEM provides enhanced flexibility, enabling variables to both depend on and influence other variables. Furthermore, SEM allows for the incorporation of mediation effects, facilitating the quantification of direct, indirect, and total effects (Grace, 2006; Lefcheck, 2016). Of particular relevance to this study is SEM’s capability to construct and model latent variables: ones that cannot be directly measured but are hypothesised to exist (Bollen, 2002; Lefcheck, 2016).

#### 2.5.2 Exploratory factor analysis

We started by constructing a latent variable representing Immunity; for this, an exploratory factor analysis (EFA) was performed using the ‘lavaan’ (version 0.6-16) package (Rosseel, 2012) in R. The latent variable model incorporated all immune assays as indicators, with “cluster= Individual ID” set to address repeated sampling and Full Information Maximum Likelihood (FIML) utilised for handling missing data. Estimation was performed using Maximum Likelihood with robust standard errors (MLR), and standardisation of values for the latent variable was implemented (”std.lv =T”). Model fit was evaluated through the examination of fit indices, including the chi-squared p-value, Comparative Fit Index (CFI), Root Mean Square Error of Approximation (RMSEA), and Standardised Root Mean Square Residual (SRMR). The conventional “rule of thumb”: CFI *>* 0.9; RMSEA *>* 0.08; SRMR *>* 0.08 was used for model fit evaluation (Hu and Bentler, 1999; MacCallum et al., 1996; Sharma et al., 2005). Predicted values for the latent construct were extracted using the “lav.predict” function from “lavaan” with default settings. During the initial assessment, it became evident that haptoglobin concentration did not significantly load on the latent variable Immunity (Fig. 1), leading to lower model fit (Table 1). As a result, haptoglobin concentration was subsequently removed and treated individually.

**Figure 1.**
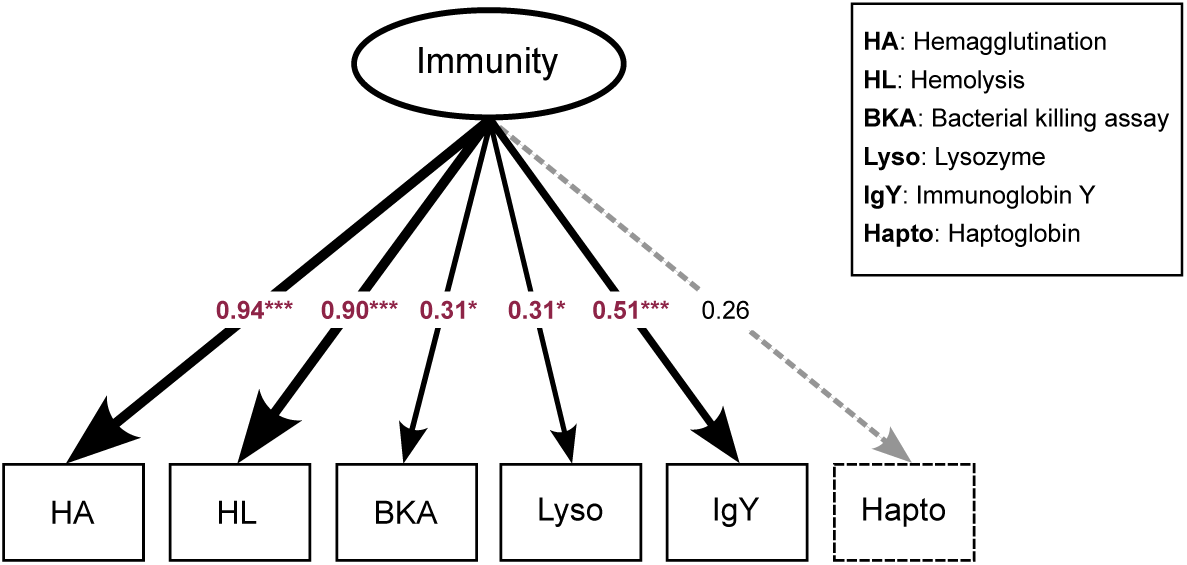
Latent variable model for *”Immunity”*. Arrows point to different immune assays, which serve as components of the latent variable. Values are indicating the strength of the relationship between the latent variable and its indicators. In red values for variables that loaded significantly in the model and black indicating non-significant loadings.

**Table 1.**
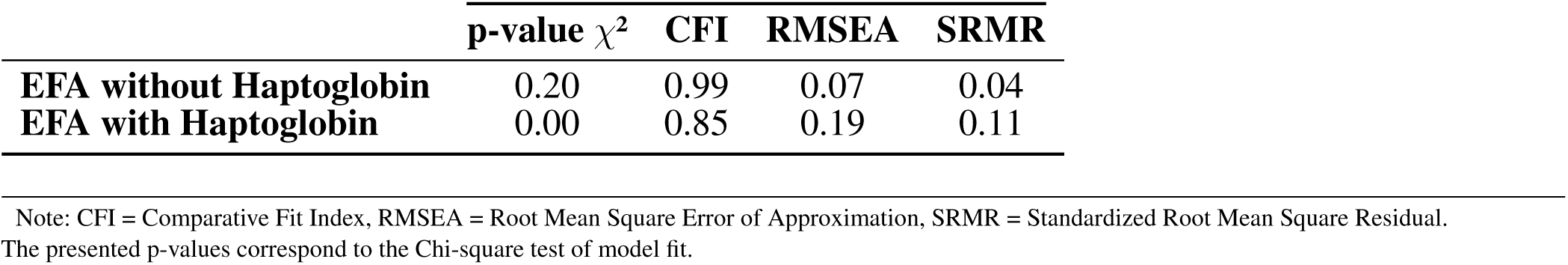
Model fit indices for exploratory factor analysis.

| | p-value $\chi^2$ | CFI | RMSEA | SRMR |
| --- | --- | --- | --- | --- |
| **EFA without Haptoglobin** | 0.20 | 0.99 | 0.07 | 0.04 |
| **EFA with Haptoglobin** | 0.00 | 0.85 | 0.19 | 0.11 |

Note: CFI = Comparative Fit Index, RMSEA = Root Mean Square Error of Approximation, SRMR = Standardized Root Mean Square Residual. The presented p-values correspond to the Chi-square test of model fit.

#### 2.5.3 Bayesian SEM

A Bayesian structural equation model (Kaplan and Depaoli, 2012) was fitted using the brms R package (Bürkner, 2017; Bürkner, 2018), which incorporates the probabilistic coding language Stan (Carpenter et al., 2017). Four distinct models were inputted into the multivariate analysis in order to test our main hypotheses:

**Path 1:**

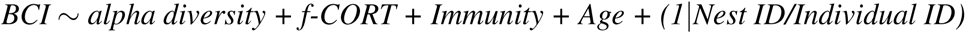

**Path 2:**

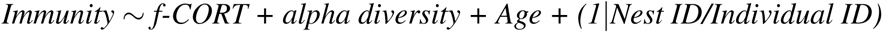

**Path 3:**

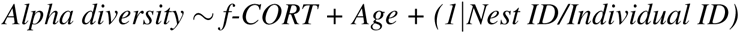

**Path 4:**

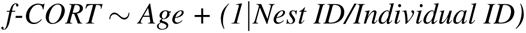

Path 1 allowed us to infer the contributions of the different components of the axis for nestling body condition, while the remaining paths were designed to capture the dynamics of the gut microbiota-immune-brain. Statistical constraints impose that we assume directionality *a priori*, although we are aware that these are bidirectional relationships. To account for repeated samples and nest sharing, a nested random effect (individual ID nested within nest ID) was incorporated into the model. Given our sampling design, where individuals of different ages were sampled at each time point (age as a continuous variable), the only way to account for age effects was to include age as a fixed effect. The model was run with four chains, each run with 100k iterations, a warm-up phase of 50k iterations and default priors. Model fit was assessed by examining the convergence of the runs, mixing of chains and performing posterior predictive checks (comparing predicted versus observed posterior distribution). Marginal and conditional Bayes *R*^2^ (Gelman et al., 2019) were calculated using the “bayes R2” function in brms. Scripts and intermediate results can be found in Appendix C & D.

### 2.6 Differential abundance analysis

Analysis of Compositions of Microbiomes with Bias Correction 2 (ANCOM-BC2) with default parameters was implemented in the R package ANCOMBC version 2.0.2 (Lin and Peddada, 2020; Lin et al., 2022). Body condition index, f-CORT, *Immunity* and age were specified as fixed effects. A nested random effect to account for Individual ID within Nest ID was fitted with the option “rand formula”. The Holm-Bonferroni method with a significance cutoff of padj *<* 0.05 was used to correct P values for multiple testing (Holm, 1979). Detailed scripts and results are presented in Appendix E.

## 3 RESULTS

We examined the potential contribution of sex, habitat and brood rank to the variation of BCI, f-CORT, and the various immune assays. The analysis revealed no substantial evidence linking body condition, corticosterone levels, or immune capacity to these variables. However, slight variations in lysozyme and haptoglobin levels were observed among different habitats. These patterns appear to be largely influenced by imbalances in the sample design and the presence of outliers (Appendix B; Fig. 9) Thus, we assume that the tested variables should not play a significant role in the SEM construct. For a comprehensive overview of the analysis pipeline and detailed results, see Appendix B.

### 3.1 Exploratory Factor Analysis

Exploratory factor analysis (EFA) derived a latent variable representing the underlying structure of immune parameter values. We synthesised the results of the immune assays into a composite variable denoted as *Immunity*, which included all of the immune parameters except haptoglobin concentration (see above). Components of the innate immune system (Hemolysis and Hemagglutination) displayed the strongest factor loadings ((*λ*_HA_ = 0.94*, λ*_HL_ = 0.90)), followed by IgY (*λ*_IgY_ = 0.50) representing the adaptive immune system (Fig.1).

### 3.2 Bayesian SEM

The Bayesian SEM revealed no evidence for a relationship between gut microbiota diversity and either *Immunity*/Haptoglobin or f-CORT (Fig.2, 3; Appendix A Fig. S1-S3). However, a negative correlation emerged between *Immunity* and f-CORT, showing a decrease in immune capacity with rising levels of f-CORT (Fig.2, 3, 4A; Appendix A Fig. S1, S2). Conversely, no evidence was found for a link between Haptoglobin and f-CORT (Fig.2, 3; Appendix A Fig. S3). A negative association was evident between f-CORT and BCI (Fig.2, 3, Fig. 4B; Appendix A Fig. S1-S3). Shannon bacterial diversity was associated with BCI, indicating that higher diversity levels corresponded to elevated BCI (µ = 0.18; CI [0.01, 0.35]; Fig. 2, 4C). Conversely, no evidence was found for the effects of *Immunity*/Haptoglobin on body condition (Fig. 2, 3; Appendix A Fig. S1-S3). Eukaryotic microbiota diversity exhibited no connection with BCI (Fig.2, 3; Appendix A Fig. S1-S3). We found no evidence for age-related effects on bacterial microbiota diversity (Fig. 2, 3; Appendix A Fig. S1, S2). However, there was credible support for a decrease of eukaryotic Faith PD and the number of observed ASV’s with age (Faith PD: µ = −0.28; CI[−0.55, −0.01]; Number of ASVs: µ = −0.28 CI[0.56, −0.01]; Appendix A Fig. S4). Additionally, f-CORT levels decreased as individuals matured, while *Immunity* increased with age (Fig. 2, 3; Appendix A Fig. S1-S4). As the results for the number of observed ASVs were similar to those for Faith PD, they are presented in the Supplementary Materials (Appendix A Fig. S1).

**Figure 2.**
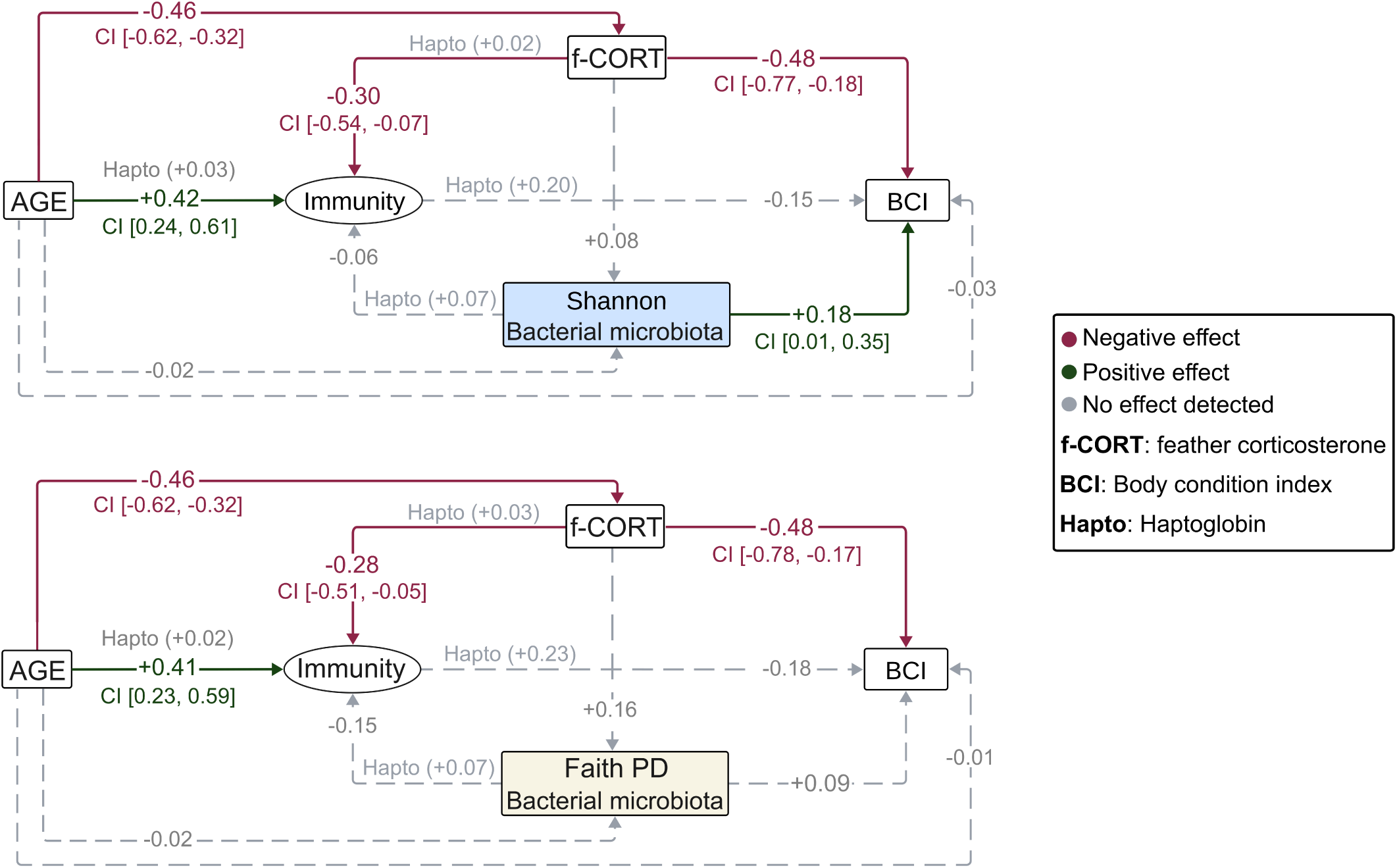
Diagrams representing the Bayesian structural equation models for the different bacterial microbiota diversity metrics. Illustrated are the results of the models incorporating the latent variable, with results for the Haptoglobin assay superimposed. Values represent the estimated mean effect of a predictor (*µ*) on the outcome variable. Credible intervals [95% CI] are provided for predictors that explain the outcome variable.

**Figure 3.**
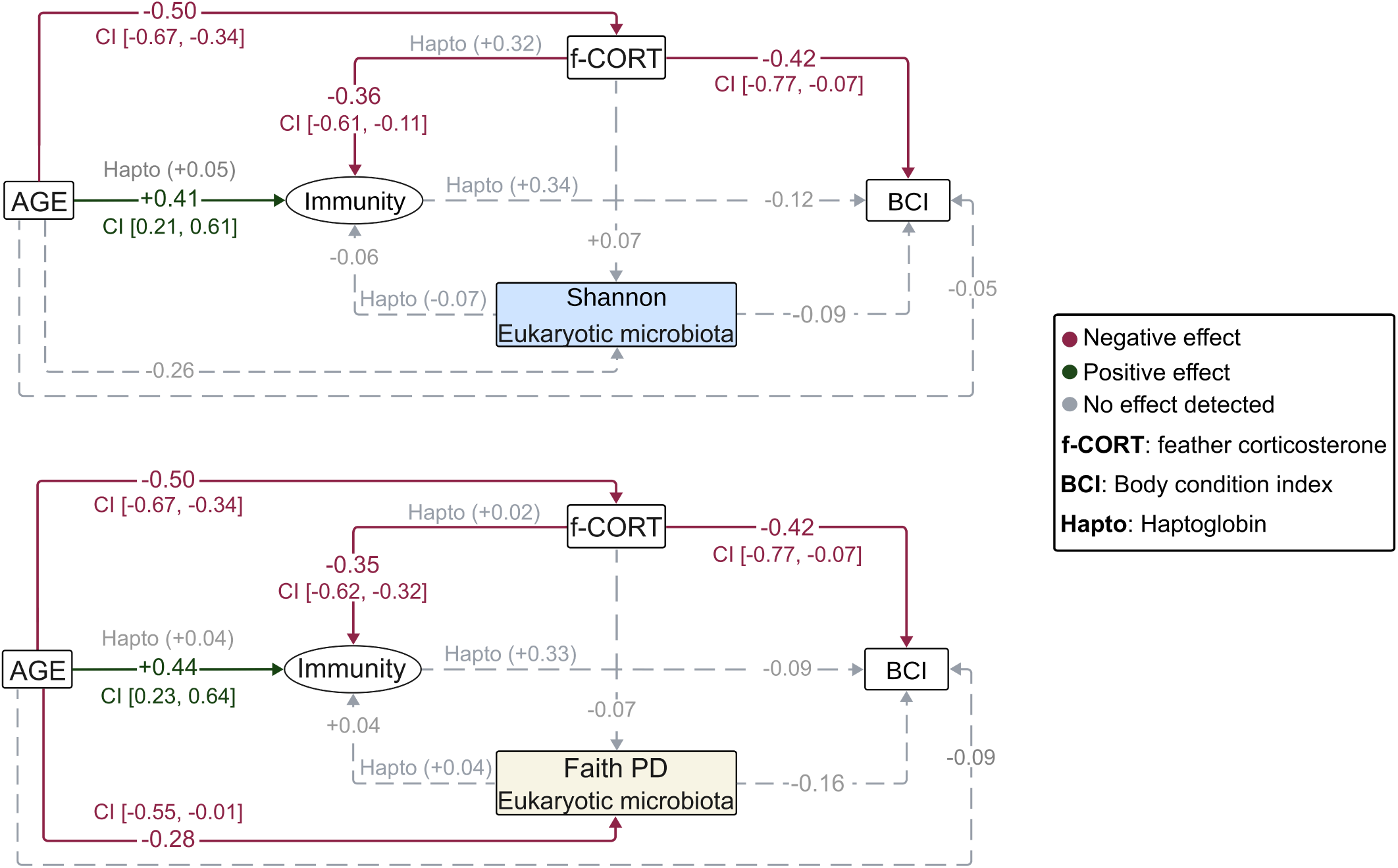
Diagrams representing the Bayesian structural equation models for the different eukaryotic microbiota diversity metrics. Illustrated are the results of the models incorporating the latent variable, with results for the Haptoglobin assay superimposed. Values represent the estimated mean effect of a predictor (µ) on the outcome variable. Credible intervals [95% CI] are provided for predictors that explain the outcome variable.

**Figure 4.**
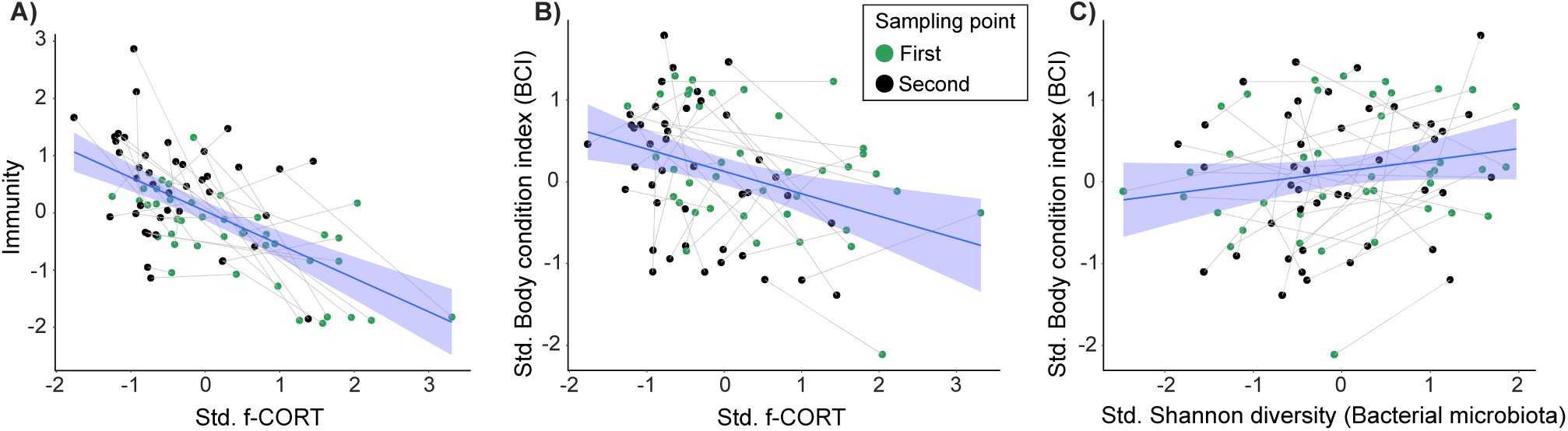
Regression plots illustrating the main results of the SEM model. **A)** Negative relationship between immunity and f-CORT; **B)** Negative correlation between body condition and f-CORT; **C)** Increase in bacterial microbiota Shannon diversity with body condition.

### 3.3 Deferentially abundant taxa

In the ANCOM-BC2 models incorporating the latent variable *Immunity*, two eukaryotic ASV’s exhibited differential abundance. Specifically, one ASV was linked to BCI (Genus: *Dothideales*), while another was associated with age (Phylum: *Phragmoplastophyta*) (see Appendix A, Fig. S5). However, the latter failed to retain statistical significance following sensitivity analysis (see Appendix A Table S22). The genus *Dothideales* demonstrated a log-fold decrease per unit of BCI (Appendix A Fig. S5). Notably, no bacterial taxa displayed differential abundance correlated with the studied variables (Appendix A Fig. S5). In contrast, when incorporating the Haptoglobin immune assay into the models, no deferentially abundant taxa were identified (Appendix A Fig S6, Table S23).

## 4 DISCUSSION

The gut microbiota-immune-brain axis shapes a variety of physiological responses through multidirectional communication among the gut microbiota, immune system, and the central nervous system (CNS) (Sylvia and Demas, 2018). This feedback system is known to influence immune function, neuroendocrine pathways, and behaviour (Martin et al., 2018; Cryan et al., 2019). Once established, disruptions to this axis may lead to the development of disorders (e.g. irritable bowel syndrome; ulcerative colitis; Alzheimer’s and Parkinson’s disease in humans) significantly impacting individual health (Rhee et al., 2009; Ghaisas et al., 2016). Here, we used gut microbial diversity measurements, various innate and adaptive immune markers and feather-corticosterone levels (a proxy for stress) to explore relationships among the three components of the axis and the repercussions to body condition in a wild population of common buzzard nestlings.

### 4.1 Gut microbial diversity, not associated with *Immunity* and stress

In contrast to our assumptions, our results revealed no association between microbial diversity, *Immunity* and stress (f-CORT) (Fig. 2, 3). However, several parts of the feedback cascade appeared to be functional (see below). Despite the well-established influence of the gut microbiota on immune system modulation and its impact on the CNS, these tripartite connections remain elusive in wild populations (Hird, 2017; Davidson et al., 2020; Madden et al., 2022; Pereira et al., 2023), particularly during the early stages of development, in contrast to the studies in humans and mice (Martin et al., 2018; Francella et al., 2022; Lynch et al., 2024).

Contrary to our findings, recent research by Francella et al. (2022) demonstrated clear links between gut microbiota, the immune system and stress in the early-life stages of laboratory-reared mice. Their study showed that immunocompromised mice had increased stress levels, decreased microbial diversity, and alterations in gut microbiome composition post-weaning; additionally, stress impacted the abundance of specific taxa that, in turn, were associated with specific behavioural traits. Furthermore, these behavioural changes were observed after just one week of age, demonstrating that stress can interact with host immunity during early development. Discrepancies between the results of the mice study and ours can be explained by various non-mutually exclusive factors. It is important to consider that their research was conducted under laboratory conditions, which lack ecologically relevant environmental factors, whereas our study was performed under ecologically realistic, natural circumstances. Indeed captivity is known to have an effect on each components of the axis (Slevin et al., 2020; Florkowski and Yorzinski, 2023), the studies also focused on different aspects of the immunity (cellular vs. humoral) and made use of immunocompromised/gene knockout mice. Furthermore species have different life-history strategies in terms of fast–slow life-history continuum (Réale et al., 2010; Bing et al., 2022).

Despite the scarcity of such distinct gut microbiota-immune-brain studies in other vertebrate species with lower degrees of experimental manipulation, studies on captive organisms have demonstrated a connection between the brain and the gut microbiota. For instance, higher gut microbiota alpha diversity in zebra finches (*Taeniopygia guttata*) has been correlated with elevated exploratory behaviour, and in house sparrows (*Passer domesticus*), beta diversity was associated with enhanced cognitive performance (Slevin et al., 2020; Florkowski and Yorzinski, 2023). It has been recognised that the gut microbiome modulates stress responses, particularly through the hypothalamic-pituitary-adrenal (HPA) axis (Foster and Neufeld, 2013); in general, higher stress levels tend to be associated with reduced microbiota diversity, specifically a reduction in the amount of rare and pathogenic taxa (Petrullo et al., 2022). While the notion of diverse microbiota being crucial for robust immunity is widely accepted (Hooper et al., 2012; Lozupone et al., 2012), extending these conclusions beyond a small number of experimental study systems is not straightforward, as several studies have found no association between microbiota diversity and immune indices. Instead, they observed that compositional differences or changes in specific taxa were more closely linked with variations in immune markers (Kreisinger et al., 2018; van Veelen et al., 2020; Fleischer et al., 2024). Our focus on microbial diversity, without considering composition (beta diversity), limits the depth of our analysis (Shade, 2017; Reese and Dunn, 2018). However, incorporating the complexities of multivariate metrics into structural equation modelling remains challenging. Nevertheless, we hope that ongoing advancements in methodologies for analysing compositional data (Sweeny et al., 2023; Fountain-Jones et al., 2024) will facilitate this approach in future studies. It is also important to consider timing in the context of stress responses, as short-term and prolonged stress can induce contrasting physiological reactions (LB, 2009). Evaluating different measures of HPA axis activity would offer a more comprehensive understanding of stress response dynamics (Stothart et al., 2019). Short-term acute stress triggers an organism’s immune response (LB, 2009) and enhances intestinal mucus secretion (Castagliuolo et al., 1996). Conversely, prolonged stress typically suppresses immunity and reduces mucus production, impacting the microbiota differently (Estienne et al., 2010; Shigeshiro et al., 2012).

Several studies of wild populations have delved into some components of the gut microbiota-immune-brain axis (Noguera et al., 2018; Petrullo et al., 2022; Berman et al., 2023), yet few have comprehensively addressed the entire system. An exception is a recent study on eastern newts (*Notophthalmus viridescens*), which explored the effects of experimentally elevated CORT levels on various immune indices and skin microbiome but did not find any clear evidence for a relationship among the three components (Pereira et al., 2023).

The examples provided demonstrate the challenges encountered in establishing associations among the components of the gut microbiome-immune-brain axis, either due to the complexities of wild settings or the specific time window investigated, or indeed, because there are no discernible associations in the corresponding systems. These challenges increase in less controlled study systems (further aggravated by the fact that studies of completely wild populations are almost non-existent), particularly when dealing with systems where certain components of the axis lack experimental challenge, alteration, or complete knockout. Given this, and considering the absence of an evident connection with the microbiome axis, we propose the following possible, not mutually exclusive mechanisms to be at play: **1.** The gut microbiota may not exert a significant influence in the initial stages of life, especially when compared to the impacts of stress and the immune system. This is supported by the robust link between f-CORT and *Immunity*, which we find despite the correlative nature of our study system (Fig. 2, 3, 4A). Consequently, we suggest that stress and immune regulation may hold greater importance for maintaining homeostasis during early development. **2.** The complexity of wild environments introduces many background effects, potentially including diverse diets, exposure to various pathogens, and the unpredictable nature of ecological interactions. These factors might overshadow the subtle and context-dependent relationships observed in more controlled settings. In reality, the connections found in these controlled environments may not be as crucial or representative of the complete picture encountered in the more realistic complexities of a wild setting. **3.** Direct comparisons between species and generalisations across species might be challenging due to ecological differences between the systems.

### 4.2 Stress, *Immunity* and body condition

The relationship between stress, immunity, and body condition is a strong feedback mechanism in animal physiology (Vagasi et al., 2018). Baseline levels of corticosterone play important roles in metabolism, development, reproduction, behaviour, and immunity (Sapolsky et al., 2000). While beneficial in the short term for resolving inflammation and preventing an overshoot of the immune responses (Dhabhar, 2018), prolonged stress-induced elevation of corticosterone compromises the immune system over time (Dhabhar and Mcewen, 1997). A study of house sparrows revealed that prolonged activation of the stress response inhibits components of the innate immune system, such as complement-mediated lysis, bacteria-killing ability, and agglutination. Indeed, numerous studies have demonstrated the detrimental effects of high stress levels, for prolonged periodes of time, on overall body condition, survival, and reproductive success (Angelier et al., 2010; Mikkelsen et al., 2023; Quirici et al., 2021).

We used feather CORT as an indicator of long-term stress. In birds, circulating CORT accumulates in developing feathers, serving as a cumulative gauge of hormone concentrations during feather growth (Jenni-Eiermann et al., 2015; Romero and Fairhurst, 2016). Our results show a negative association between *Immunity* and f-CORT, supporting the established concept that immunosuppression is expected in the face of allostatic overload (chronic stress) (Romero et al., 2009). Additionally, prolonged stress can incur costs for developing individuals, may divert resources away from essential physiological processes, and impair an individual’s ability to fend off infections and maintain overall well-being, as evidenced here by a decline in body condition (Fig. 2, 3, 4A-B).

### 4.3 Bacterial microbiota diversity and body condition

Our results show that nestlings with higher Shannon diversity have better body condition (Fig. 2, 4C). This positive association can be attributed to the enhanced resistance of more diverse gut communities against pathogenic invasion, increased stability and resilience to disturbance (Buffie and Pamer, 2013). Additionally, a more diverse microbiota can potentially offer a broader range of functions performed by various bacterial taxa, leading to benefits for the host (Heiman and Greenway, 2016). Conversely, lower microbiota diversity is typically viewed as being detrimental to hosts (Le Chatelier et al., 2013), as it implies a loss of essential functions that could result in reduced nutrient assimilation or immunodeficiency (Round and Mazmanian, 2009; Hanning and Diaz-Sanchez, 2015). However, it has also been shown that high diversity might be associated with a state of dysbiosis, so a reduction in diversity could signal a return to homeostasis (Johnson and Burnet, 2016; Kohl et al., 2018; Coyte and Rakoff-Nahoum, 2019). Our results align with the Anna Karenina principle, which states that changes in microbiota due to disturbances lead to shifts from stable to unstable community states (Zaneveld et al., 2017). We propose that the increase in Shannon diversity in buzzard nestlings indicates an increase in stable and abundant taxa, which may offer greater benefits to the host (Hanning and Diaz-Sanchez, 2015). Metrics like Faith’s PD and the number of ASVs treat all taxa equally (Faith and Baker, 2006), potentially explaining why they do not show associations with body condition (Fig. 3; Appendix A Fig. S1).

The absence of clear links between eukaryotic microbiota diversity and body condition may indicate a slower rate of change in the eukaryotic microbiota. Bacterial and eukaryotic taxa are likely to have distinct roles within the gut ecosystem (Oever and Netea, 2014; Vemuri et al., 2020). Fast-changing bacterial taxa may experience stronger competition, leading to positive selection on the core functionally-relevant taxa (Abt and Pamer, 2014; Coyte and Rakoff-Nahoum, 2019), whereas changes in the more complex and less abundant eukaryotic taxa (Chin et al., 2020) may occur at a slower pace and therefore not be observed during early stages of host development.

### 4.4 Age effects on stress, *Immunity* and eukaryotic microbiota diversity

The positive relationship between age and *Immunity*, and the negative association of age and stress levels (f-CORT; Fig. 2, 3), most likely reflect nestling development and the maturation of the immune system. Initially, nestlings rely on innate and maternal transferred immunity (Klasing and Leshchinsky, 1999; Palacios et al., 2009). As they mature, exposure to pathogens and foreign microbes through the environment and diet, contact with nest material and siblings, challenges and stimulates the immune system, thus aiding its maturation (Lochmiller and Deerenberg, 2000; Morais et al., 2020; Oldereid et al., 2023). During the ontogenetic development of birds, exposure to corticosterone occurs in both the prenatal (embryonic) and postnatal (nestling and fledgling) periods (Henriksen et al., 2011; Strange et al., 2016). In the prenatal stage, corticosterone is transferred from the mother to the embryo through the egg yolk, and is influenced by the maternal environment, e.g. exposure to predators, competitors and other stress-inducing cues during egg production, and differences in environmental quality (Hayward and Wingfield, 2004; Saino et al., 2005; Love et al., 2008). In the postnatal stage, an initial surge in corticosterone levels may serve as an adaptive response to stressors associated with hatching, exposure to a new environment, and nutritional demands, being beneficial in the short term (Chin et al., 2009; Spencer et al., 2009; Strange et al., 2016). As nestlings mature, the HPA axis undergoes maturation, leading to improved stress response regulation. Initially, nestlings invest significant energy in growth and development. As they progress through early life stages, maturation enables more efficient allocation and prioritisation of resources (Smulders, 2021; Spencer et al., 2009). As mentioned earlier, the maturation of both physiological systems is not independent; instead, a bidirectional interaction regulates both the immune system and the HPA axis (Francella et al., 2022).

We find a decline in eukaryotic microbiota phylogenetic diversity and the number of ASV’s with age. Dominant taxa show higher resilience to disturbance, securing their positions via selective filtering and the occupation of core niches (Costello et al., 2012; Abt and Pamer, 2014; Coyte and Rakoff-Nahoum, 2019). Rare taxa are primarily acquired through stochastic processes but still significantly contribute to certain community alpha diversity measures (like Faith PD and n° of observed ASVs). Over time, these rare taxa are progressively replaced by the more dominant taxa (Shade et al., 2014). The dominant taxa also show closer phylogenetic relationships to one another, suggesting potential specialisation or competitive advantages driving their prevalence (Janiak et al., 2021; West et al., 2022; Davies et al., 2022). This hints at an initial diverse gut microbiota in newborns due to rapid post-hatching colonisation (Trevelline et al., 2018; Pereira et al., 2024).

## 5 CONCLUSION

We investigated the interactions of the gut microbiota-immune-brain axis in raptor nestlings. As far as we are aware, this represents one of few studies exploring this axis in a wild vertebrate population and incorporating different time points during the nestling phase in a longitudinal study design. While there was no conclusive evidence for the microbiota-immune-brain axis in our study system, we did find evidence for the hypothesised relationships among stress, *Immunity*, and body condition. Elevated f-CORT levels were linked to immunosuppression and adverse effects on overall body condition, suggesting that immune and stress regulation play a dominant role in early nestling development. Shannon bacterial microbiota diversity was positively correlated with nestling body condition, suggesting a potential benefit of a diverse and stable gut microbiota. Age plays a crucial role in influencing immune development, stress responses, and eukaryotic microbial diversity. The decline in eukaryotic microbial diversity with age implies an early uncontrolled gut colonisation, followed by selective removal of non-relevant taxa. Our study thereby contributes to a growing body of knowledge on the dynamics of the gut microbiota-immune-brain axis in wild populations.

## Supporting information

Associations-sex-habitat-rank

Bayesian-SEM-immunity

Bayesian-SEM-Haptoglobin

DAA

f-CORT-assay-validation

## 6 ETHICS STATMENT

Sampling of common buzzards nestlings followed the ARRIVE guidelines and were approved by the Animal Ethics Committee at Bielefeld University and permitted by the local authority Kreis Gütersloh, permit number: 4.5.2-723-Bussard and by the ethics committee of the Animal Care and Use Committee of the German North Rhine-Westphalia State Office for Nature, Environment and Consumer Protection (Landesamt für Nature, Umwelt und Verbraucherschutz Nordrhein-Westfalen) under permit numbers: 84-02.04.2014.A091, 84-02-04.2017.A147.

## AUTHOR CONTRIBUTIONS

The research project was conceptualised by HP, NC, JIH, and OK. HP participated in data collection, conducted bioinformatic and statistical analyses, and drafted the manuscript with input from NC, JIH, OK, and GÁC. Fieldwork and sample collection were carried out by NC, MO, TR, KPG, and OK. GÁC conducted the immune assays, while SK oversaw the corticosterone determination procedures. BC and ÖM provided scientific advice and contributed to the development of the methodology. All authors reviewed and approved the final manuscript.

## ACKNOWLEDGMENTS

We express our gratitude to Thomas Grünkorn and Justine Léauté for their dedicated data collection efforts during the field season. Special thanks to Rebecca Chen for her valuable guidance and assistance in statistical analysis. We acknowledge the contributions of Sabine Kruse and Edith Ossendorf in the corticosterone determination process and Katja Pohle during the immunological analyses. We acknowledge the financial support of the German Research Foundation (DFG). The funding body had no involvement in the study design, data collection, analysis, interpretation, or manuscript writing.

## SUPPLEMENTARY MATERIAL

Appendix A - Supplementary tables & figures

Appendix B - Testing associations with sex, habitat and rank

Appendix C - Bayesian SEM for the latent variable *Immunity*

Appendix D - Bayesian SEM for the Haptoglobin immune assay

Appendix E - Differential abundance analysis

Appendix F - f-CORT assay validation

## DATA AVAILABILITY STATEMENT

All 16S and 28S rRNA raw reads have been submitted to the European Nucleotide Archive repository, Project ID: PRJEB70791. The scripts and metadata to reproduce all analyses can be accessed via the GitHub repository: Gut-immune-brain-axis-common-buzzard-nestlings

