## Supplementary materials for "The gut microbiota-immune-brain axis in a wild vertebrate: dynamic interactions and health impacts"

### Supplementary Material

#### Table of Contents

**Supplementary Material**

- Table of Contents
- Table S1. Summary of number of individuals per marker
- Table S2. Summary of final ASV table: 16s rRNA
- Table S3. Summary of final ASV table: 28s rRNA
- 1. Exploratory factor analysis for the latent variable "Immunity"
  - Table S4. Latent variable model results excluding Haptoglobin assay
  - Table S5. Latent variable model resullts incorporating Haptoglobin assay
- Figure S1. Diagrams illustrating the Bayesian structural equation models for the number of bacterial and eukaryotic ASVs.
- Figure S2. Posterior estimates (points) and 95% credible intervals (horizontal lines) from Bayesian structural equation models that include the latent variable "Immunity"
- Figure S3. Posterior estimates (points) and 95% credible intervals (horizontal lines) from Bayesian structural equation model for Haptoglobin immune assay
- Figure S4. Regression plots depicting the age effects obtained from brmsSEM: age effects on *Immunity* and f-CORT, eukaryotic Faith PD and n° of observed ASVs
- 2. Bayesian Structural Equation Modeling (SEM) diagnostics of 16S rRNA diversity measurements - models incorporating the latent variable "Immunity"
  - Table S6. Shannon diversity index
  - Table S7. Faith PD
  - Table S8. N° of observed ASV's
  - Table S9. Bayes R2 for each diversity measurement
- 3. Bayesian Structural Equation Modeling (SEM) diagnostics of 28S rRNA diversity measurements - models incorporating the latent variable "Immunity"
  - Table S10. Shannon diversity index
  - Table S11. Faith PD
  - Table S12. N° of observed ASV's
  - Table S13. Bayes R2 for each diversity measurement
- 4. Bayesian Structural Equation Modeling (SEM) diagnostics of 16S rRNA diversity measurements - model results for Haptoglobin immune assay
  - Table S14. Shannon diversity index
  - Table S15. Faith PD
  - Table S16. N° of Observed ASV's
  - Table S17. Bayes R2 for each diversity measurement
- 5. Bayesian Structural Equation Modeling (SEM) diagnostics of 28S rRNA diversity measurements - model results for Haptoglobin immune assay
  - Table S18. Shannon diversity index
  - Table S19. Faith PD
  - Table S20. N° of observed ASV's
  - Table S21. Bayes R2 for each diversity measurement
- Figure S5. Differential abundance analysis results for each of the variables in study - ANCOM-BC2 model incorporating the latent variable *Immunity*
  - Table S22. Sensitivity analysis for the two deferentially abundant taxa
- Figure S6. Differential abundance analysis results for each of the variables in study - ANCOM-BC2 model incorporating Haptoglobin immune assay
  - Table S23. Sensitivity analysis for the deferentially abundant taxa
- Figure S7. Bayesian structural equation models for the different bacterial diversity measures with results from each immune assay superimposed onto each diagram

Figure S8. Bayesian structural equation models for the different eukaryotic diversity measures with results from each immune assay superimposed onto each diagram

#### Table S1. Summary of number of individuals per marker

| | N individuals | N samples | Males | Females | Nests |
|---|---|---|---|---|---|
| **16s rRNA dataset** | 43 | 86 | 24 | 19 | 23 |
| **28s rRNA dataset** | 42 | 72 | 24 | 18 | 23 |

#### Table S2. Summary of final ASV table: 16s rRNA

| | Sample |
|---|---|
| **Number of samples** | 86 |
| **Number of features (ASVs)** | 2,072 |
| **Total frequency** | 20,336 |

#### Table S3. Summary of final ASV table: 28s rRNA

| | Sample |
|---|---|
| **Number of samples** | 72 |
| **Number of features (ASVs)** | 1768 |
| **Total frequency** | 31,502 |

### 1. Exploratory factor analysis for the latent variable "Immunity"

##### Table S4. Latent variable model results excluding Haptoglobin assay

| | | | | | | |
|---|---|---|---|---|---|---|
| **Latent variables:** | | | | | | |
| | **Estimate** | **Std.Err** | **z-value** | **P(>\|z\|)** | **Std.lv** | **Std.all** |
| immunity =~ | | | | | | |
| std_ha | 0.94 | 0.09 | 10.01 | 0.00 | 0.94 | 0.94 |
| std_hl | 0.90 | 0.10 | 8.76 | 0.00 | 0.90 | 0.90 |

| | | | | | | |
|---|---|---|---|---|---|---|
| std_bka | 0.31 | 0.12 | 2.58 | 0.01 | 0.31 | 0.31 |
| std_lyso | 0.31 | 0.12 | 2.69 | 0.01 | 0.31 | 0.31 |
| std_igy | 0.53 | 0.11 | 4.78 | 0.00 | 0.53 | 0.53 |
| **Intercepts:** | | | | | | |
| | **Estimate** | **Std.Err** | **z-value** | **P(>\|z\|)** | **Std.lv** | **Std.all** |
| std_ha | 0.00 | 0.10 | 0.00 | 1.00 | 0.00 | 0.00 |
| std_hl | 0.00 | 0.11 | 0.00 | 1.00 | 0.00 | 0.00 |
| std_bka | 0.00 | 0.10 | 0.00 | 1.00 | 0.00 | 0.00 |
| std_lyso | -0.01 | 0.11 | -0.08 | 0.94 | -0.01 | -0.01 |
| std_igy | 0.00 | 0.11 | 0.00 | 1.00 | 0.00 | 0.00 |
| immunity | 0.00 | | | | 0.00 | 0.00 |
| **Variances:** | | | | | | |
| | **Estimate** | **Std.Err** | **z-value** | **P(>\|z\|)** | **Std.lv** | **Std.all** |
| std_ha | 0.112 | 0.089 | 1.261 | 0.207 | 0.112 | 0.113 |
| std_hl | 0.181 | 0.089 | 2.032 | 0.042 | 0.181 | 0.183 |
| std_bka | 0.895 | 0.164 | 5.456 | 0.000 | 0.895 | 0.906 |
| std_lyso | 0.894 | 0.223 | 4.010 | 0.000 | 0.894 | 0.903 |
| std_igy | 0.709 | 0.116 | 6.092 | 0.000 | 0.709 | 0.717 |
| immunity | 1.000 | | | | 1.000 | 1.000 |
| **R-Square:** | **Estimate** | | | | | |
| std_ha | 0.887 | | | | | |
| std_hl | 0.817 | | | | | |
| std_bka | 0.094 | | | | | |
| std_lyso | 0.097 | | | | | |
| std_igy | 0.283 | | | | | |

#### Table S5. Latent variable model resullts incorporating Haptoglobin assay

| | | | | | | |
|---|---|---|---|---|---|---|
| **Latent Variables:** | | | | | | |
| | **Estimate** | **Std.Err** | **z-value** | **P(>\|z\|)** | **Std.lv** | **Std.all** |
| **immunity =~** | | | | | | |
| std_ha | 0.93 | 0.09 | 10.20 | 0.00 | 0.93 | 0.93 |
| std_hl | 0.91 | 0.10 | 9.20 | 0.00 | 0.91 | 0.91 |

| | | | | | | |
|---|---|---|---|---|---|---|
| std_bka | 0.31 | 0.11 | 2.77 | 0.01 | 0.31 | 0.31 |
| std_lyso | 0.32 | 0.12 | 2.64 | 0.01 | 0.32 | 0.32 |
| std_hapto | 0.26 | 0.15 | 1.76 | 0.08 | 0.26 | 0.26 |
| std_igy | 0.53 | 0.11 | 4.73 | 0.00 | 0.53 | 0.53 |
| **Intercepts:** | | | | | | |
| | **Estimate** | **Std.Err** | **z-value** | **P(>\|z\|)** | **Std.lv** | **Std.all** |
| .std_ha | 0.00 | 0.10 | 0.00 | 1.00 | 0.00 | 0.00 |
| .std_hl | 0.00 | 0.11 | 0.00 | 1.00 | 0.00 | 0.00 |
| .std_bka | 0.00 | 0.10 | 0.00 | 1.00 | 0.00 | 0.00 |
| .std_lyso | -0.01 | 0.11 | -0.08 | 0.94 | -0.01 | -0.01 |
| .std_hapto | -0.01 | 0.11 | -0.09 | 0.93 | -0.01 | -0.01 |
| .std_igy | 0.00 | 0.11 | 0.00 | 1.00 | 0.00 | 0.00 |
| immunity | 0.00 | | | | 0.00 | 0.00 |
| **Variances:** | | | | | | |
| | **Estimate** | **Std.Err** | **z-value** | **P(>\|z\|)** | **Std.lv** | **Std.all** |
| .std_ha | 0.13 | 0.07 | 1.79 | 0.07 | 0.13 | 0.13 |
| .std_hl | 0.17 | 0.07 | 2.31 | 0.02 | 0.17 | 0.17 |
| .std_bka | 0.89 | 0.16 | 5.53 | 0.00 | 0.89 | 0.90 |
| .std_lyso | 0.89 | 0.22 | 4.09 | 0.00 | 0.89 | 0.90 |
| .std_hapto | 0.92 | 0.44 | 2.10 | 0.04 | 0.92 | 0.93 |
| .std_igy | 0.71 | 0.12 | 6.11 | 0.00 | 0.71 | 0.72 |
| immunity | 1.00 | | | | 1.00 | 1.00 |
| **R-Square:** | **Estimate** | | | | | |
| std_ha | 0.87 | | | | | |
| std_hl | 0.83 | | | | | |
| std_bka | 0.10 | | | | | |
| std_lyso | 0.10 | | | | | |
| std_hapto | 0.07 | | | | | |
| std_igy | 0.28 | | | | | |

#### Figure S1. Diagrams illustrating the Bayesian structural equation models for the number of bacterial and eukaryotic ASVs.

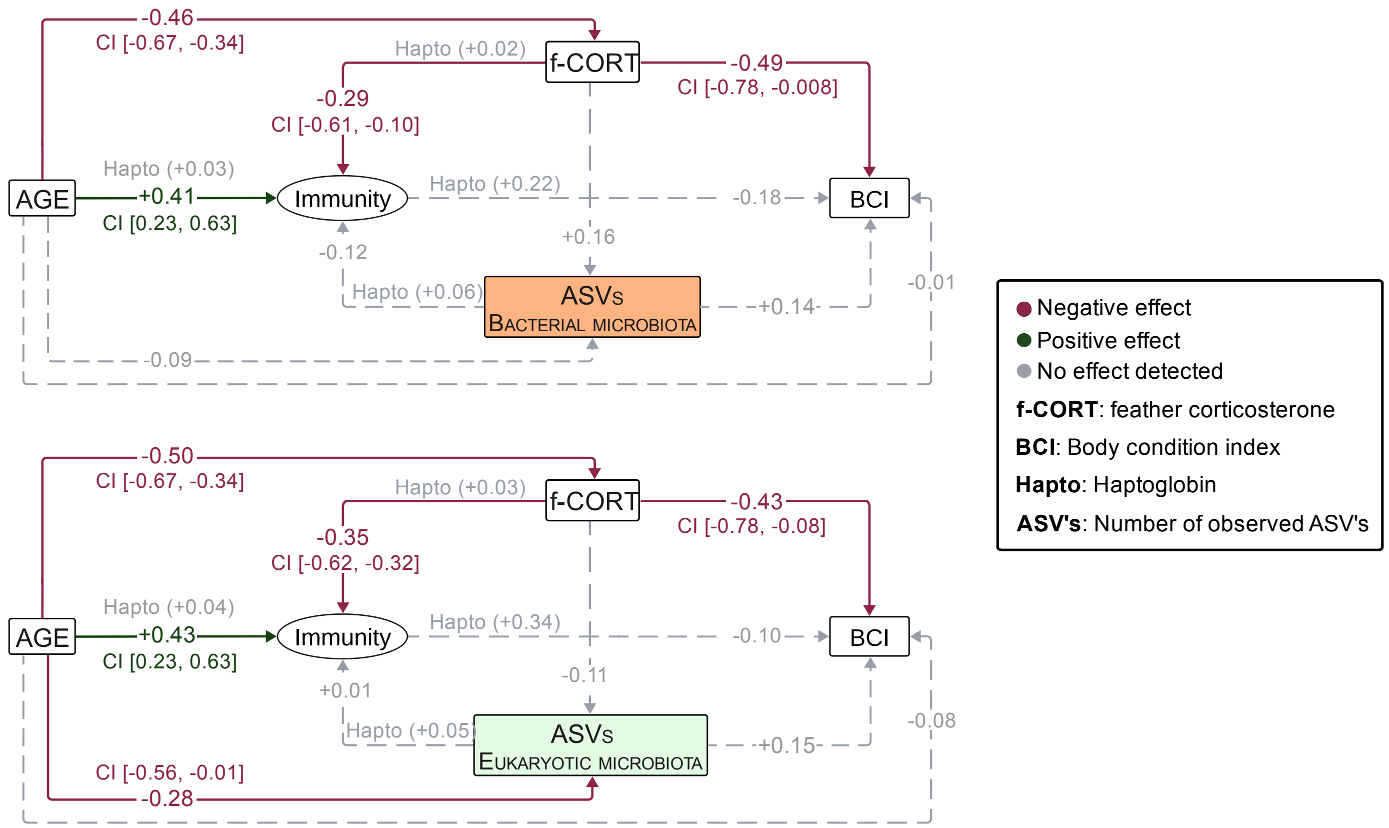

### Figure S2. Posterior estimates (points) and 95% credible intervals (horizontal lines) from Bayesian structural equation models that include the latent variable "Immunity"

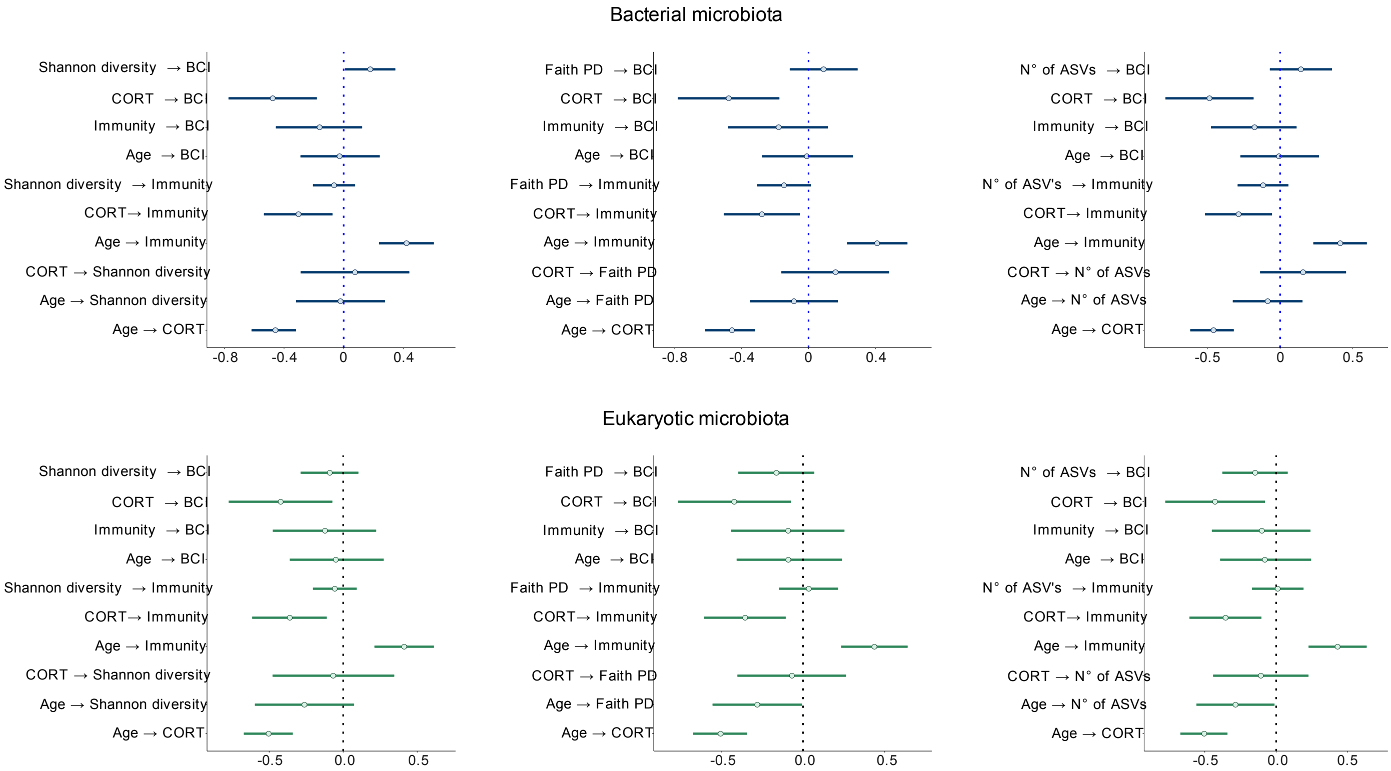

**Note:** A predictor confidently explains the outcome variable when the 95% credible intervals do not overlap 0.

### Figure S3. Posterior estimates (points) and 95% credible intervals (horizontal lines) from Bayesian structural equation model for Haptoglobin immune assay

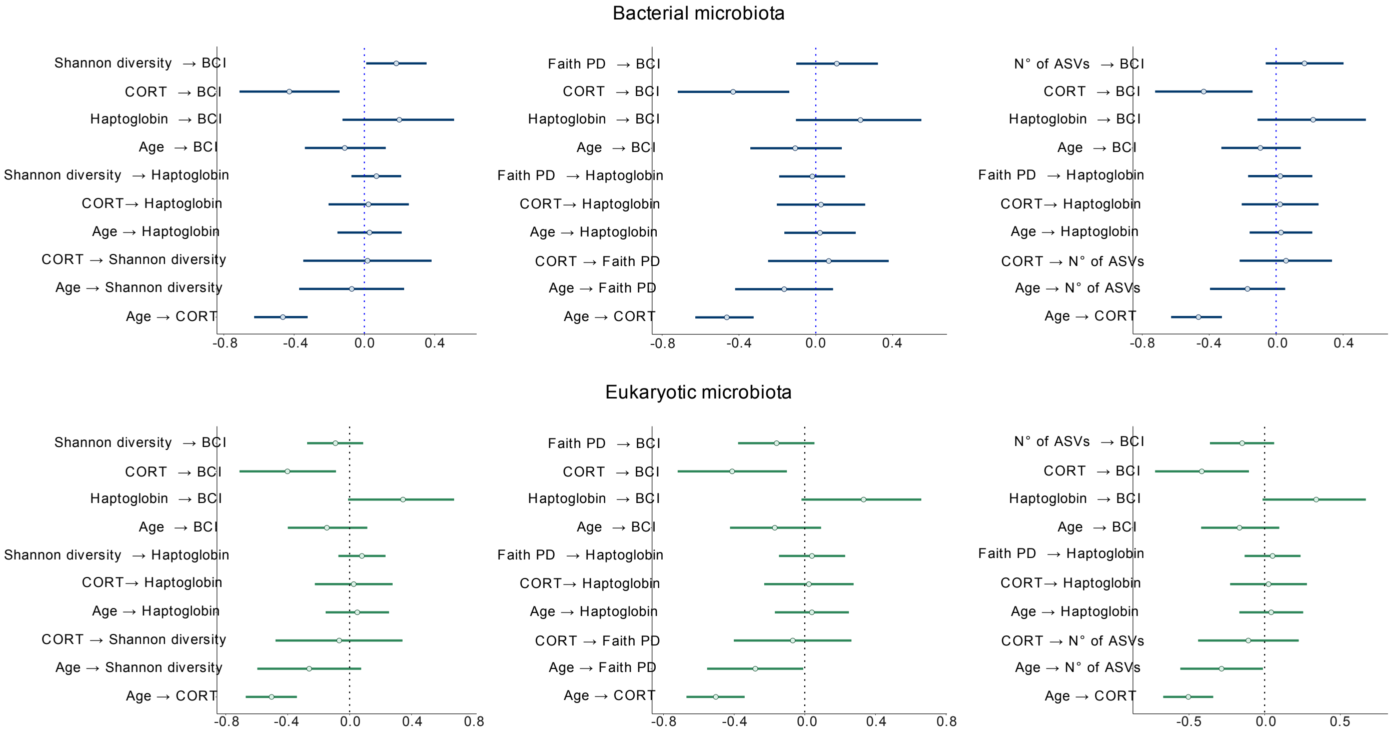

**Note:** A predictor confidently explains the outcome variable when the 95% credible intervals do not overlap 0.

### Figure S4. Regression plots depicting the age effects obtained from brmsSEM: age effects on *Immunity* and f-CORT, eukaryotic Faith PD and n° of observed ASVs

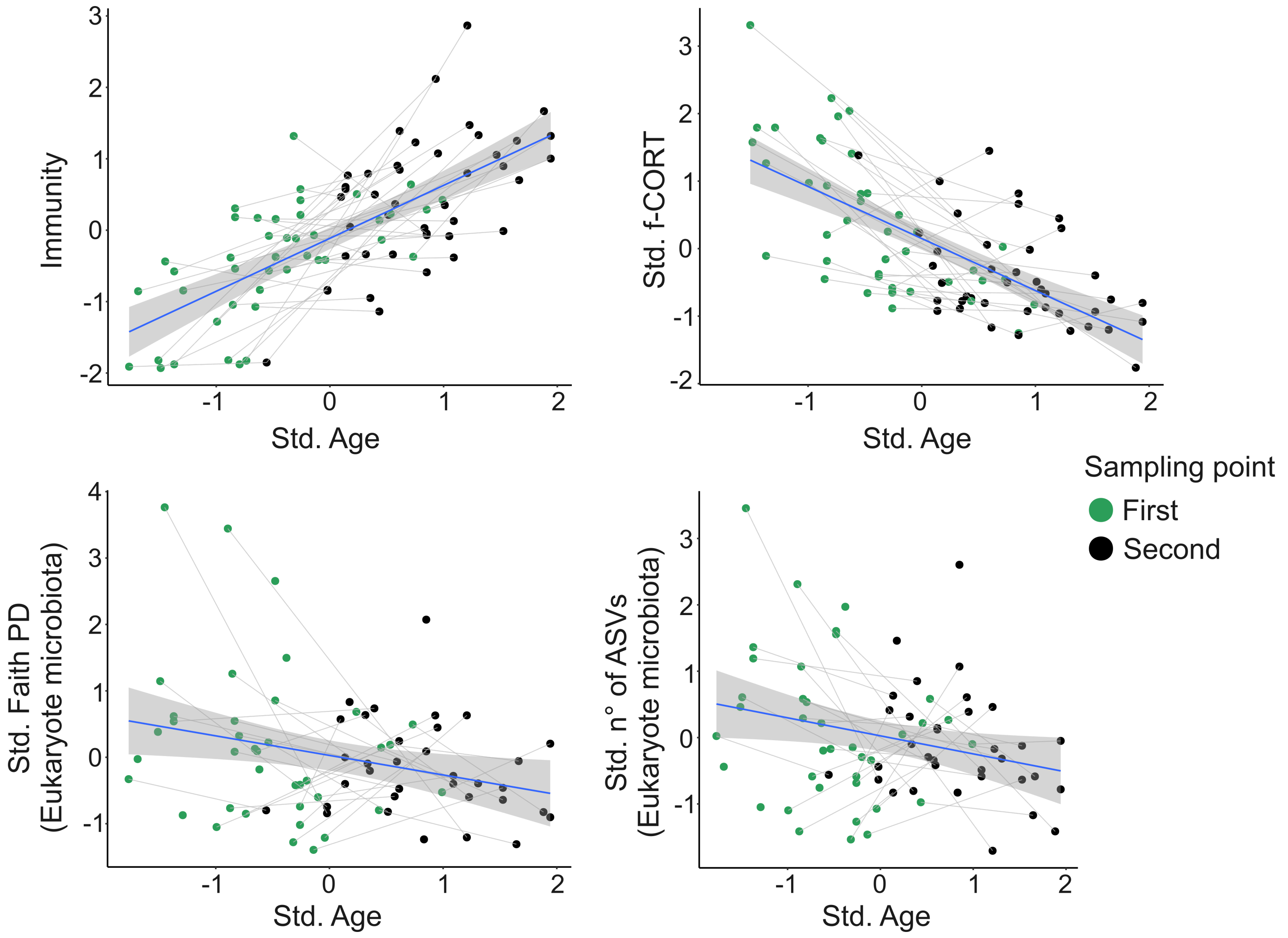

**Note:** Samples from the same individual at two different time points are linked by gray lines.

#### 2. Bayesian Structural Equation Modeling (SEM) diagnostics of 16S rRNA diversity measurements - models incorporating the latent variable "Immunity"

##### Table S6. Shannon diversity index

| | | | | | | | |
|---|---|---|---|---|---|---|---|
| **Group-Level Effects:** | | | | | | | |
| **nest (Number of levels: 23)** | | | | | | | |
| | **Estimate** | **Est.Error** | **l-95% CI** | **u-95% CI** | **Rhat** | **Bulk_ESS** | **Tail_ESS** |
| sd(BCI_Intercept) | 0.10 | 0.03 | 0.03 | 0.17 | 1.00 | 40193 | 29967 |
| sd(Immunity_Intercept) | 0.04 | 0.02 | 0.00 | 0.09 | 1.00 | 53009 | 75934 |
| sd(Shannon_Intercept) | 0.05 | 0.03 | 0.00 | 0.12 | 1.00 | 73171 | 83559 |
| sd(cort_Intercept) | 0.04 | 0.03 | 0.00 | 0.10 | 1.00 | 42186 | 67727 |
| **nest:ring_number (Number of levels: 43)** | | | | | | | |
| | **Estimate** | **Est.Error** | **l-95% CI** | **u-95% CI** | **Rhat** | **Bulk_ESS** | **Tail_ESS** |

| | | | | | | | |
|---|---|---|---|---|---|---|---|
| sd(BCI_Intercept) | 0.05 | 0.03 | 0.00 | 0.12 | 1.00 | 34067 | 63400 |
| sd(Immunity_Intercept) | 0.03 | 0.02 | 0.00 | 0.08 | 1.00 | 55421 | 79441 |
| sd(Shannon_Intercept) | 0.04 | 0.03 | 0.00 | 0.10 | 1.00 | 85824 | 85327 |
| sd(CORT_Intercept) | 0.03 | 0.02 | 0.00 | 0.09 | 1.00 | 42011 | 76190 |
| **Population-Level Effects:** | | | | | | | |
| | **Estimate** | **Est.Error** | **l-95% CI** | **u-95% CI** | **Rhat** | **Bulk_ESS** | **Tail_ESS** |
| BCI_Intercept | 0.72 | 0.12 | 0.48 | 0.96 | 1.00 | 171059 | 152106 |
| Immunity_Intercept | 0.33 | 0.09 | 0.15 | 0.51 | 1.00 | 164599 | 148738 |
| shannon_Intercept | 0.54 | 0.14 | 0.27 | 0.80 | 1.00 | 152069 | 142448 |
| CORT_Intercept | 0.59 | 0.04 | 0.51 | 0.68 | 1.00 | 136837 | 147761 |
| BCI_Shannon | 0.18 | 0.09 | 0.01 | 0.35 | 1.00 | 233151 | 157272 |
| BCI_CORT | -0.48 | 0.15 | -0.77 | -0.18 | 1.00 | 168613 | 149179 |
| BCI_Immunity | -0.16 | 0.15 | -0.45 | 0.12 | 1.00 | 135230 | 140900 |
| BCI_Age | -0.03 | 0.14 | -0.29 | 0.24 | 1.00 | 116247 | 130704 |
| Immunity_Shannon | -0.06 | 0.07 | -0.21 | 0.08 | 1.00 | 261721 | 154686 |
| Immunity_CORT | -0.30 | 0.12 | -0.54 | -0.07 | 1.00 | 151185 | 148688 |
| Immunity_Age | 0.42 | 0.09 | 0.24 | 0.61 | 1.00 | 167508 | 151875 |
| Shannon_CORT | 0.08 | 0.19 | -0.29 | 0.44 | 1.00 | 160936 | 152577 |
| Shannon_Age | -0.02 | 0.15 | -0.32 | 0.28 | 1.00 | 165292 | 153792 |
| CORT_Age | -0.46 | 0.08 | -0.62 | -0.32 | 1.00 | 127711 | 150763 |
| **Family Specific Parameters:** | | | | | | | |
| | **Estimate** | **Est.Error** | **l-95% CI** | **u-95% CI** | **Rhat** | **Bulk_ESS** | **Tail_ESS** |
| sigma_BCI | 0.16 | 0.02 | 0.13 | 0.19 | 1.00 | 87455 | 122865 |
| sigma_Immunity | 0.14 | 0.01 | 0.11 | 0.17 | 1.00 | 137061 | 142505 |
| sigma_Shannon | 0.23 | 0.02 | 0.19 | 0.27 | 1.00 | 219034 | 145803 |
| sigma_CORT | 0.14 | 0.01 | 0.11 | 0.17 | 1.00 | 86466 | 132787 |
| alpha_CORT | 4.51 | 2.72 | -0.57 | 10.31 | 1.00 | 98159 | 125900 |

#### Table S7. Faith PD

| | | | | | | | |
|---|---|---|---|---|---|---|---|
| **Group-Level Effects:** | | | | | | | |
| **nest (Number of levels: 23)** | | | | | | | |
| | **Estimate** | **Est.Error** | **l-95% CI** | **u-95% CI** | **Rhat** | **Bulk_ESS** | **Tail_ESS** |
| sd(BCI_Intercept) | 0.09 | 0.04 | 0.02 | 0.17 | 1.00 | 41835 | 38783 |

| | | | | | | | |
|---|---|---|---|---|---|---|---|
| sd(Immunity_Intercept) | 0.03 | 0.02 | 0.00 | 0.08 | 1.00 | 66785 | 90315 |
| sd(FaithPD_Intercept) | 0.04 | 0.03 | 0.00 | 0.11 | 1.00 | 78895 | 93112 |
| sd(cort_Intercept) | 0.04 | 0.03 | 0.00 | 0.10 | 1.00 | 46183 | 76222 |
| **nest:ring_number (Number of levels: 43)** | | | | | | | |
| | **Estimate** | **Est.Error** | **l-95% CI** | **u-95% CI** | **Rhat** | **Bulk_ESS** | **Tail_ESS** |
| sd(BCI_Intercept) | 0.05 | 0.03 | 0.00 | 0.13 | 1.00 | 36858 | 69470 |
| sd(Immunity_Intercept) | 0.03 | 0.02 | 0.00 | 0.08 | 1.00 | 72728 | 93078 |
| sd(FaithPD_Intercept) | 0.04 | 0.03 | 0.00 | 0.10 | 1.00 | 86094 | 94672 |
| sd(CORT_Intercept) | 0.03 | 0.02 | 0.00 | 0.09 | 1.00 | 45689 | 86842 |
| **Population-Level Effects:** | | | | | | | |
| | **Estimate** | **Est.Error** | **l-95% CI** | **u-95% CI** | **Rhat** | **Bulk_ESS** | **Tail_ESS** |
| BCI_Intercept | 0.78 | 0.13 | 0.53 | 1.03 | 1.00 | 227253 | 158046 |
| Immunity_Intercept | 0.35 | 0.09 | 0.18 | 0.53 | 1.00 | 217297 | 152915 |
| FaithPD_Intercept | 0.41 | 0.12 | 0.18 | 0.65 | 1.00 | 201266 | 150828 |
| CORT_Intercept | 0.59 | 0.04 | 0.51 | 0.68 | 1.00 | 163851 | 158106 |
| BCI_FaithPD | 0.09 | 0.10 | -0.11 | 0.29 | 1.00 | 241820 | 154659 |
| BCI_CORT | -0.48 | 0.16 | -0.78 | -0.17 | 1.00 | 221946 | 161268 |
| BCI_Immunity | -0.18 | 0.15 | -0.48 | 0.12 | 1.00 | 176801 | 155537 |
| BCI_Age | -0.01 | 0.14 | -0.28 | 0.27 | 1.00 | 148433 | 151120 |
| Immunity_FaithPD | -0.15 | 0.08 | -0.31 | 0.01 | 1.00 | 281668 | 150324 |
| Immunity_CORT | -0.28 | 0.12 | -0.51 | -0.05 | 1.00 | 192357 | 158346 |
| Immunity_Age | 0.41 | 0.09 | 0.23 | 0.59 | 1.00 | 214708 | 162819 |
| FaithPD_CORT | 0.16 | 0.16 | -0.16 | 0.48 | 1.00 | 204099 | 161576 |
| FaithPD_Age | -0.09 | 0.13 | -0.35 | 0.18 | 1.00 | 212116 | 161601 |
| CORT_Age | -0.46 | 0.08 | -0.62 | -0.32 | 1.00 | 153,019.85 | 159,412.54 |
| **Family Specific Parameters:** | | | | | | | |
| | **Estimate** | **Est.Error** | **l-95% CI** | **u-95% CI** | **Rhat** | **Bulk_ESS** | **Tail_ESS** |
| sigma_BCI | 0.16 | 0.02 | 0.13 | 0.20 | 1.00 | 92480 | 124753 |
| sigma_Immunity | 0.14 | 0.01 | 0.12 | 0.16 | 1.00 | 195882 | 149848 |
| sigma_FaithPD | 0.20 | 0.02 | 0.17 | 0.24 | 1.00 | 241036 | 152012 |
| sigma_CORT | 0.14 | 0.01 | 0.11 | 0.17 | 1.00 | 97127 | 139061 |
| alpha_CORT | 4.53 | 2.72 | -0.56 | 10.33 | 1.00 | 114529 | 138100 |

#### Table S8. N° of observed ASV's

| | | | | | | | |
|---|---|---|---|---|---|---|---|
| **Group-Level Effects:** | | | | | | | |
| **nest (Number of levels: 23)** | | | | | | | |
| | **Estimate** | **Est.Error** | **l-95% CI** | **u-95% CI** | **Rhat** | **Bulk_ESS** | **Tail_ESS** |
| sd(BCI_Intercept) | 0.09 | 0.04 | 0.02 | 0.17 | 1.00 | 42963 | 38463 |
| sd(Immunity_Intercept) | 0.03 | 0.02 | 0.00 | 0.09 | 1.00 | 63091 | 90077 |
| sd(ASV_Intercept) | 0.04 | 0.03 | 0.00 | 0.10 | 1.00 | 81158 | 94442 |
| sd(cort_Intercept) | 0.04 | 0.03 | 0.00 | 0.10 | 1.00 | 46623 | 73961 |
| **nest:ring_number (Number of levels: 43)** | | | | | | | |
| | **Estimate** | **Est.Error** | **l-95% CI** | **u-95% CI** | **Rhat** | **Bulk_ESS** | **Tail_ESS** |
| sd(BCI_Intercept) | 0.05 | 0.03 | 0.00 | 0.13 | 1.00 | 38749 | 74160 |
| sd(Immunity_Intercept) | 0.03 | 0.02 | 0.00 | 0.08 | 1.00 | 68082 | 90282 |
| sd(ASV_Intercept) | 0.03 | 0.02 | 0.00 | 0.08 | 1.00 | 101607 | 98436 |
| sd(CORT_Intercept) | 0.03 | 0.02 | 0.00 | 0.09 | 1.00 | 45851 | 87339 |
| **Population-Level Effects:** | | | | | | | |
| | **Estimate** | **Est.Error** | **l-95% CI** | **u-95% CI** | **Rhat** | **Bulk_ESS** | **Tail_ESS** |
| BCI_Intercept | 0.77 | 0.12 | 0.52 | 1.01 | 1.00 | 221975 | 158089 |
| Immunity_Intercept | 0.33 | 0.09 | 0.16 | 0.51 | 1.00 | 209396 | 146766 |
| ASV_Intercept | 0.33 | 0.11 | 0.12 | 0.55 | 1.00 | 204922 | 146073 |
| CORT_Intercept | 0.59 | 0.04 | 0.51 | 0.68 | 1.00 | 164348 | 158239 |
| BCI_ASV | 0.14 | 0.11 | -0.07 | 0.36 | 1.00 | 278133 | 153518 |
| BCI_CORT | -0.49 | 0.15 | -0.79 | -0.18 | 1.00 | 216771 | 153897 |
| BCI_Immunity | -0.18 | 0.15 | -0.48 | 0.11 | 1.00 | 175698 | 144150 |
| BCI_Age | -0.01 | 0.14 | -0.27 | 0.27 | 1.00 | 149260 | 143820 |
| Immunity_ASV | -0.12 | 0.09 | -0.29 | 0.06 | 1.00 | 273104 | 155751 |
| Immunity_CORT | -0.29 | 0.12 | -0.52 | -0.06 | 1.00 | 187571 | 151720 |
| Immunity_Age | 0.41 | 0.09 | 0.23 | 0.60 | 1.00 | 213409 | 156429 |
| ASV_CORT | 0.16 | 0.15 | -0.14 | 0.45 | 1.00 | 209768 | 159805 |
| ASV_Age | -0.09 | 0.12 | -0.33 | 0.15 | 1.00 | 213539 | 159926 |
| CORT_Age | -0.46 | 0.08 | -0.62 | -0.32 | 1.00 | 153667 | 162653 |
| **Family Specific Parameters:** | | | | | | | |
| | **Estimate** | **Est.Error** | **l-95% CI** | **u-95% CI** | **Rhat** | **Bulk_ESS** | **Tail_ESS** |
| sigma_BCI | 0.16 | 0.02 | 0.13 | 0.20 | 1.00 | 94855 | 130217 |
| sigma_Immunity | 0.14 | 0.01 | 0.12 | 0.16 | 1.00 | 183033 | 152022 |
| sigma_ASV | 0.19 | 0.02 | 0.16 | 0.22 | 1.00 | 261966 | 150278 |

| | | | | | | | |
|---|---|---|---|---|---|---|---|
| sigma_CORT | 0.14 | 0.01 | 0.11 | 0.17 | 1.00 | 101905 | 141945 |
| alpha_CORT | 4.52 | 2.72 | -0.58 | 10.33 | 1.00 | 113172 | 139327 |

##### Table S9. Bayes R2 for each diversity measurement

| | R2m | Est.Error | Q2.5 | Q97.5 | R2c | Est.Error | Q2.5 | Q97.5 |
|---|---|---|---|---|---|---|---|---|
| **BCI (Path1)** | 0.19 | 0.07 | 0.07 | 0.32 | 0.47 | 0.08 | 0.28 | 0.61 |
| **Immunity (Path2)** | 0.53 | 0.05 | 0.41 | 0.61 | 0.57 | 0.06 | 0.44 | 0.67 |
| **Shanonn (Path3)** | 0.03 | 0.03 | 0.00 | 0.11 | 0.10 | 0.06 | 0.01 | 0.25 |
| **CORT (Path4)** | 0.35 | 0.08 | 0.19 | 0.51 | 0.46 | 0.11 | 0.24 | 0.65 |
| | **R2m** | **Est.Error** | **Q2.5** | **Q97.5** | **R2c** | **Est.Error** | **Q2.5** | **Q97.5** |
| **BCI (Path1)** | 0.17 | 0.07 | 0.05 | 0.30 | 0.44 | 0.09 | 0.24 | 0.59 |
| **Immunity (Path2)** | 0.54 | 0.05 | 0.42 | 0.63 | 0.57 | 0.05 | 0.45 | 0.67 |
| **Faith PD (Path3)** | 0.08 | 0.05 | 0.00 | 0.19 | 0.15 | 0.07 | 0.03 | 0.31 |
| **CORT (Path4)** | 0.35 | 0.08 | 0.19 | 0.51 | 0.46 | 0.11 | 0.24 | 0.65 |
| | **R2m** | **Est.Error** | **Q2.5** | **Q97.5** | **R2c** | **Est.Error** | **Q2.5** | **Q97.5** |
| **BCI (Path1)** | 0.18 | 0.07 | 0.06 | 0.31 | 0.45 | 0.09 | 0.25 | 0.60 |
| **Immunity (Path2)** | 0.53 | 0.05 | 0.41 | 0.62 | 0.57 | 0.06 | 0.44 | 0.67 |
| **ASV (Path3)** | 0.08 | 0.05 | 0.01 | 0.20 | 0.15 | 0.07 | 0.03 | 0.29 |
| **CORT (Path4)** | 0.35 | 0.08 | 0.19 | 0.51 | 0.46 | 0.11 | 0.24 | 0.65 |

#### 3. Bayesian Structural Equation Modeling (SEM) diagnostics of 28S rRNA diversity measurements - models incorporating the latent variable "Immunity"

##### Table S10. Shannon diversity index

| | | | | | | | |
|---|---|---|---|---|---|---|---|
| **Group-Level Effects:** | | | | | | | |
| **nest (Number of levels: 23)** | | | | | | | |
| | **Estimate** | **Est.Error** | **l-95% CI** | **u-95% CI** | **Rhat** | **Bulk_ESS** | **Tail_ESS** |
| sd(BCI_Intercept) | 0.10 | 0.04 | 0.01 | 0.18 | 1.00 | 43440 | 47029 |
| sd(Immunity_Intercept) | 0.05 | 0.03 | 0.00 | 0.12 | 1.00 | 46497 | 80602 |

| | | | | | | | |
|---|---|---|---|---|---|---|---|
| sd(Shannon_Intercept) | 0.05 | 0.04 | 0.00 | 0.13 | 1.00 | 88109 | 100503 |
| sd(CORT_Intercept) | 0.05 | 0.03 | 0.00 | 0.12 | 1.00 | 44588 | 73408 |
| **nest:ring_number (Number of levels: 41)** | | | | | | | |
| | **Estimate** | **Est.Error** | **l-95% CI** | **u-95% CI** | **Rhat** | **Bulk_ESS** | **Tail_ESS** |
| sd(BCI_Intercept) | 0.05 | 0.04 | 0.00 | 0.13 | 1.00 | 51191 | 82715 |
| sd(Immunity_Intercept) | 0.04 | 0.03 | 0.00 | 0.09 | 1.00 | 57078 | 97148 |
| sd(Shannon_Intercept) | 0.05 | 0.04 | 0.00 | 0.13 | 1.00 | 86785 | 101805 |
| sd(CORT_Intercept) | 0.04 | 0.03 | 0.00 | 0.09 | 1.00 | 52082 | 94318 |
| **Population-Level Effects:** | | | | | | | |
| | **Estimate** | **Est.Error** | **l-95% CI** | **u-95% CI** | **Rhat** | **Bulk_ESS** | **Tail_ESS** |
| BCI_Intercept | 0.84 | 0.16 | 0.53 | 1.15 | 1.00 | 257492 | 157983 |
| Immunity_Intercept | 0.35 | 0.11 | 0.14 | 0.56 | 1.00 | 210031 | 153012 |
| shannon_Intercept | 0.77 | 0.15 | 0.47 | 1.07 | 1.00 | 234697 | 151070 |
| CORT_Intercept | 0.62 | 0.05 | 0.53 | 0.72 | 1.00 | 235559 | 171146 |
| BCI_Shannon | -0.09 | 0.10 | -0.29 | 0.10 | 1.00 | 299121 | 151667 |
| BCI_CORT | -0.42 | 0.18 | -0.77 | -0.07 | 1.00 | 241261 | 161921 |
| BCI_Immunity | -0.12 | 0.18 | -0.48 | 0.22 | 1.00 | 177201 | 151056 |
| BCI_Age | -0.05 | 0.16 | -0.36 | 0.27 | 1.00 | 159380 | 146270 |
| Immunity_Shannon | -0.06 | 0.07 | -0.20 | 0.09 | 1.00 | 323971 | 160165 |
| Immunity_CORT | -0.36 | 0.13 | -0.61 | -0.11 | 1.00 | 183336 | 158445 |
| Immunity_Age | 0.41 | 0.10 | 0.21 | 0.61 | 1.00 | 226141 | 162064 |
| Shannon_CORT | -0.07 | 0.21 | -0.48 | 0.35 | 1.00 | 227697 | 159630 |
| Shannon_Age | -0.26 | 0.17 | -0.60 | 0.07 | 1.00 | 237093 | 162077 |
| CORT_Age | -0.50 | 0.08 | -0.67 | -0.34 | 1.00 | 205533 | 170103 |
| **Family Specific Parameters:** | | | | | | | |
| | **Estimate** | **Est.Error** | **l-95% CI** | **u-95% CI** | **Rhat** | **Bulk_ESS** | **Tail_ESS** |
| sigma_BCI | 0.17 | 0.02 | 0.13 | 0.21 | 1.00 | 94141 | 129645 |
| sigma_Immunity | 0.13 | 0.01 | 0.11 | 0.16 | 1.00 | 125995 | 148081 |
| sigma_Shannon | 0.24 | 0.02 | 0.19 | 0.28 | 1.00 | 250653 | 151912 |
| sigma_CORT | 0.14 | 0.02 | 0.11 | 0.17 | 1.00 | 115629 | 143287 |
| alpha_CORT | 3.11 | 2.71 | -1.78 | 8.93 | 1.00 | 105352 | 121969 |

#### Table S11. Faith PD

| | | | | | | | |
|---|---|---|---|---|---|---|---|
| **Group-Level Effects:** | | | | | | | |
| **nest (Number of levels: 23)** | | | | | | | |
| | **Estimate** | **Est.Error** | **l-95% CI** | **u-95% CI** | **Rhat** | **Bulk_ESS** | **Tail_ESS** |
| sd(BCI_Intercept) | 0.10 | 0.04 | 0.01 | 0.18 | 1.00 | 44108 | 50748 |
| sd(Immunity_Intercept) | 0.05 | 0.03 | 0.00 | 0.11 | 1.00 | 48959 | 85818 |
| sd(FaithPD_Intercept) | 0.03 | 0.03 | 0.00 | 0.10 | 1.00 | 107439 | 111926 |
| sd(CORT_Intercept) | 0.05 | 0.03 | 0.00 | 0.12 | 1.00 | 47224 | 78299 |
| **nest:ring_number (Number of levels: 41)** | | | | | | | |
| | **Estimate** | **Est.Error** | **l-95% CI** | **u-95% CI** | **Rhat** | **Bulk_ESS** | **Tail_ESS** |
| sd(BCI_Intercept) | 0.05 | 0.03 | 0.00 | 0.13 | 1.00 | 54540 | 88979 |
| sd(Immunity_Intercept) | 0.04 | 0.03 | 0.00 | 0.10 | 1.00 | 56684 | 99592 |
| sd(FaithPD_Intercept) | 0.04 | 0.03 | 0.00 | 0.10 | 1.00 | 104537 | 108735 |
| sd(CORT_Intercept) | 0.04 | 0.03 | 0.00 | 0.09 | 1.00 | 55175 | 95121 |
| **Population-Level Effects:** | | | | | | | |
| | **Estimate** | **Est.Error** | **l-95% CI** | **u-95% CI** | **Rhat** | **Bulk_ESS** | **Tail_ESS** |
| BCI_Intercept | 0.84 | 0.14 | 0.56 | 1.11 | 1.00 | 278684 | 160729 |
| Immunity_Intercept | 0.29 | 0.10 | 0.09 | 0.49 | 1.00 | 243151 | 154233 |
| FaithPD_Intercept | 0.45 | 0.13 | 0.20 | 0.69 | 1.00 | 271811 | 154780 |
| CORT_Intercept | 0.62 | 0.05 | 0.53 | 0.72 | 1.00 | 250796 | 171645 |
| BCI_FaithPD | -0.16 | 0.12 | -0.40 | 0.07 | 1.00 | 340822 | 161568 |
| BCI_CORT | -0.42 | 0.18 | -0.77 | -0.07 | 1.00 | 273914 | 160657 |
| BCI_Immunity | -0.09 | 0.18 | -0.44 | 0.25 | 1.00 | 192620 | 149586 |
| BCI_Age | -0.09 | 0.16 | -0.41 | 0.24 | 1.00 | 165692 | 146731 |
| Immunity_FaithPD | 0.04 | 0.09 | -0.15 | 0.22 | 1.00 | 341797 | 155995 |
| Immunity_CORT | -0.35 | 0.13 | -0.60 | -0.11 | 1.00 | 204113 | 163518 |
| Immunity_Age | 0.44 | 0.10 | 0.23 | 0.64 | 1.00 | 251992 | 163363 |
| FaithPD_CORT | -0.07 | 0.17 | -0.40 | 0.26 | 1.00 | 264045 | 164815 |
| FaithPD_Age | -0.28 | 0.14 | -0.55 | -0.01 | 1.00 | 264698 | 163463 |
| CORT_Age | -0.50 | 0.08 | -0.67 | -0.34 | 1.00 | 217734 | 170138 |
| **Family Specific Parameters:** | | | | | | | |
| | **Estimate** | **Est.Error** | **l-95% CI** | **u-95% CI** | **Rhat** | **Bulk_ESS** | **Tail_ESS** |
| sigma_BCI | 0.17 | 0.02 | 0.13 | 0.21 | 1.00 | 98802 | 132296 |
| sigma_Immunity | 0.13 | 0.01 | 0.11 | 0.16 | 1.00 | 122836 | 149890 |
| sigma_FaithPD | 0.19 | 0.02 | 0.16 | 0.23 | 1.00 | 316900 | 148549 |

| | | | | | | | |
|---|---|---|---|---|---|---|---|
| sigma_CORT | 0.14 | 0.02 | 0.11 | 0.17 | 1.00 | 122290 | 142987 |
| alpha_CORT | 3.11 | 2.72 | -1.78 | 8.96 | 1.00 | 114427 | 131169 |

#### Table S12. N° of observed ASV's

| | | | | | | | |
|---|---|---|---|---|---|---|---|
| **Group-Level Effects:** | | | | | | | |
| **nest (Number of levels: 23)** | | | | | | | |
| | **Estimate** | **Est.Error** | **l-95% CI** | **u-95% CI** | **Rhat** | **Bulk_ESS** | **Tail_ESS** |
| sd(BCI_Intercept) | 0.10 | 0.04 | 0.01 | 0.18 | 1.00 | 41708 | 43096 |
| sd(Immunity_Intercept) | 0.05 | 0.03 | 0.00 | 0.11 | 1.00 | 46701 | 75331 |
| sd(ASV_Intercept) | 0.03 | 0.02 | 0.00 | 0.09 | 1.00 | 111097 | 94406 |
| sd(CORT_Intercept) | 0.05 | 0.03 | 0.00 | 0.12 | 1.00 | 42664 | 69630 |
| **nest:ring_number (Number of levels: 41)** | | | | | | | |
| | **Estimate** | **Est.Error** | **l-95% CI** | **u-95% CI** | **Rhat** | **Bulk_ESS** | **Tail_ESS** |
| sd(BCI_Intercept) | 0.05 | 0.03 | 0.00 | 0.13 | 1.00 | 49548 | 82494 |
| sd(Immunity_Intercept) | 0.04 | 0.03 | 0.00 | 0.10 | 1.00 | 52291 | 87013 |
| sd(ASV_Intercept) | 0.04 | 0.03 | 0.00 | 0.11 | 1.00 | 85524 | 92005 |
| sd(CORT_Intercept) | 0.04 | 0.03 | 0.00 | 0.09 | 1.00 | 46236 | 90325 |
| **Population-Level Effects:** | | | | | | | |
| | **Estimate** | **Est.Error** | **l-95% CI** | **u-95% CI** | **Rhat** | **Bulk_ESS** | **Tail_ESS** |
| BCI_Intercept | 0.84 | 0.14 | 0.56 | 1.13 | 1.00 | 193481 | 152637 |
| Immunity_Intercept | 0.30 | 0.10 | 0.09 | 0.50 | 1.00 | 169514 | 144525 |
| ASV_Intercept | 0.52 | 0.13 | 0.27 | 0.77 | 1.00 | 182509 | 140896 |
| CORT_Intercept | 0.62 | 0.05 | 0.53 | 0.72 | 1.00 | 186895 | 161248 |
| BCI_ASV | -0.15 | 0.12 | -0.38 | 0.08 | 1.00 | 254699 | 156980 |
| BCI_CORT | -0.43 | 0.18 | -0.78 | -0.08 | 1.00 | 189445 | 155658 |
| BCI_Immunity | -0.10 | 0.18 | -0.45 | 0.24 | 1.00 | 159502 | 150111 |
| BCI_Age | -0.08 | 0.16 | -0.39 | 0.24 | 1.00 | 136058 | 137626 |
| Immunity_ASV | 0.01 | 0.09 | -0.17 | 0.19 | 1.00 | 245230 | 156711 |
| Immunity_CORT | -0.35 | 0.13 | -0.61 | -0.10 | 1.00 | 157148 | 154855 |
| Immunity_Age | 0.43 | 0.10 | 0.23 | 0.63 | 1.00 | 185126 | 157534 |
| ASV_CORT | -0.11 | 0.17 | -0.44 | 0.23 | 1.00 | 189017 | 151191 |
| ASV_Age | -0.28 | 0.14 | -0.56 | -0.01 | 1.00 | 190401 | 153832 |
| CORT_Age | -0.50 | 0.08 | -0.67 | -0.34 | 1.00 | 168839 | 164270 |

| | | | | | | | |
|---|---|---|---|---|---|---|---|
| **Family Specific Parameters:** | | | | | | | |
| | **Estimate** | **Est.Error** | **l-95% CI** | **u-95% CI** | **Rhat** | **Bulk_ESS** | **Tail_ESS** |
| sigma_BCI | 0.17 | 0.02 | 0.13 | 0.21 | 1.00 | 89208 | 125835 |
| sigma_Immunity | 0.13 | 0.01 | 0.11 | 0.16 | 1.00 | 110137 | 141585 |
| sigma_ASV | 0.19 | 0.02 | 0.16 | 0.23 | 1.00 | 215140 | 149817 |
| sigma_CORT | 0.14 | 0.02 | 0.11 | 0.17 | 1.00 | 105791 | 134827 |
| alpha_CORT | 3.13 | 2.72 | -1.73 | 9.00 | 1.00 | 92657 | 111289 |

#### Table S13. Bayes R2 for each diversity measurement

| | **R2m** | **Est.Error** | **Q2.5** | **Q97.5** | **R2c** | **Est.Error** | **Q2.5** | **Q97.5** |
|---|---|---|---|---|---|---|---|---|
| **BCI (Path1)** | 0.16 | 0.07 | 0.04 | 0.30 | 0.41 | 0.10 | 0.19 | 0.58 |
| **Immunity (Path2)** | 0.58 | 0.05 | 0.46 | 0.66 | 0.64 | 0.06 | 0.50 | 0.74 |
| **Shannon (Path3)** | 0.08 | 0.05 | 0.00 | 0.20 | 0.16 | 0.08 | 0.03 | 0.33 |
| **CORT (Path4)** | 0.41 | 0.09 | 0.22 | 0.56 | 0.54 | 0.10 | 0.30 | 0.70 |
| | **R2m** | **Est.Error** | **Q2.5** | **Q97.5** | **R2c** | **Est.Error** | **Q2.5** | **Q97.5** |
| **BCI (Path1)** | 0.17 | 0.07 | 0.04 | 0.30 | 0.42 | 0.10 | 0.20 | 0.59 |
| **Immunity (Path2)** | 0.58 | 0.05 | 0.45 | 0.66 | 0.64 | 0.06 | 0.50 | 0.74 |
| **Faith PD (Path3)** | 0.11 | 0.06 | 0.01 | 0.25 | 0.17 | 0.07 | 0.04 | 0.33 |
| **CORT (Path4)** | 0.41 | 0.09 | 0.22 | 0.56 | 0.54 | 0.10 | 0.30 | 0.70 |
| | **R2m** | **Est.Error** | **Q2.5** | **Q97.5** | **R2c** | **Est.Error** | **Q2.5** | **Q97.5** |
| **BCI (Path1)** | 0.16 | 0.07 | 0.04 | 0.30 | 0.42 | 0.10 | 0.20 | 0.59 |
| **Immunity (Path2)** | 0.58 | 0.05 | 0.45 | 0.65 | 0.63 | 0.06 | 0.50 | 0.74 |
| **ASV (Path3)** | 0.10 | 0.06 | 0.01 | 0.24 | 0.17 | 0.08 | 0.04 | 0.33 |
| **CORT (Path4)** | 0.41 | 0.09 | 0.22 | 0.56 | 0.54 | 0.10 | 0.30 | 0.70 |

### 4. Bayesian Structural Equation Modeling (SEM) diagnostics of 16S rRNA diversity measurements - model results for Haptoglobin immune assay

#### Table S14. Shannon diversity index

| | | | | | | | |
|---|---|---|---|---|---|---|---|
| **Group-Level Effects:** | | | | | | | |
| **nest (Number of levels: 23)** | | | | | | | |
| | **Estimate** | **Est.Error** | **l-95% CI** | **u-95% CI** | **Rhat** | **Bulk_ESS** | **Tail_ESS** |
| sd(BCI_Intercept) | 0.12 | 0.04 | 0.05 | 0.20 | 1.00 | 46577 | 40288 |
| sd(Haptoglobin_Intercept) | 0.06 | 0.03 | 0.00 | 0.12 | 1.00 | 37280 | 61394 |
| sd(Shannon_Intercept) | 0.05 | 0.04 | 0.00 | 0.13 | 1.00 | 70193 | 93652 |
| sd(CORT_Intercept) | 0.04 | 0.02 | 0.00 | 0.09 | 1.00 | 49165 | 80164 |
| **nest:ring_number (Number of levels: 43)** | | | | | | | |
| | **Estimate** | **Est.Error** | **l-95% CI** | **u-95% CI** | **Rhat** | **Bulk_ESS** | **Tail_ESS** |
| sd(BCI_Intercept) | 0.05 | 0.04 | 0.00 | 0.13 | 1.00 | 34914 | 59787 |
| sd(Haptoglobin_Intercept) | 0.04 | 0.02 | 0.00 | 0.09 | 1.00 | 52859 | 91654 |
| sd(Shannon_Intercept) | 0.04 | 0.03 | 0.00 | 0.11 | 1.00 | 90472 | 99033 |
| sd(CORT_Intercept) | 0.03 | 0.02 | 0.00 | 0.08 | 1.00 | 49587 | 92268 |
| **Population-Level Effects:** | | | | | | | |
| | **Estimate** | **Est.Error** | **l-95% CI** | **u-95% CI** | **Rhat** | **Bulk_ESS** | **Tail_ESS** |
| BCI_Intercept | 0.65 | 0.12 | 0.42 | 0.88 | 1.00 | 166370 | 152064 |
| Haptoglobin_Intercept | 0.07 | 0.09 | -0.12 | 0.25 | 1.00 | 207964 | 151016 |
| shannon_Intercept | 0.58 | 0.14 | 0.31 | 0.84 | 1.00 | 214929 | 148041 |
| CORT_Intercept | 0.59 | 0.04 | 0.51 | 0.69 | 1.00 | 170215 | 163200 |
| BCI_Shannon | 0.18 | 0.09 | 0.01 | 0.35 | 1.00 | 279225 | 157629 |
| BCI_CORT | -0.43 | 0.15 | -0.71 | -0.14 | 1.00 | 176535 | 155057 |
| BCI_Haptoglobin | 0.20 | 0.16 | -0.12 | 0.51 | 1.00 | 146181 | 141461 |
| BCI_Age | -0.11 | 0.12 | -0.34 | 0.12 | 1.00 | 150130 | 154761 |
| Haptoglobin_Shannon | 0.07 | 0.07 | -0.07 | 0.21 | 1.00 | 306014 | 156540 |
| Haptoglobin_CORT | 0.02 | 0.12 | -0.20 | 0.25 | 1.00 | 196783 | 158952 |
| Haptoglobin_Age | 0.03 | 0.09 | -0.15 | 0.21 | 1.00 | 210761 | 157411 |
| Shannon_CORT | 0.02 | 0.19 | -0.35 | 0.38 | 1.00 | 212876 | 160860 |
| Shannon_Age | -0.07 | 0.15 | -0.37 | 0.23 | 1.00 | 220302 | 164092 |
| CORT_Age | -0.47 | 0.08 | -0.63 | -0.32 | 1.00 | 151514 | 161846 |
| **Family Specific Parameters:** | | | | | | | |
| | **Estimate** | **Est.Error** | **l-95% CI** | **u-95% CI** | **Rhat** | **Bulk_ESS** | **Tail_ESS** |
| sigma_BCI | 0.15 | 0.02 | 0.12 | 0.18 | 1.00 | 83069 | 120711 |
| sigma_Haptoglobin | 0.13 | 0.01 | 0.11 | 0.16 | 1.00 | 107411 | 142534 |
| sigma_Shannon | 0.22 | 0.02 | 0.19 | 0.27 | 1.00 | 237812 | 149810 |

| | | | | | | | |
|---|---|---|---|---|---|---|---|
| sigma_CORT | 0.14 | 0.01 | 0.11 | 0.17 | 1.00 | 107827 | 137420 |
| alpha_CORT | 4.49 | 2.73 | -0.60 | 10.26 | 1.00 | 120661 | 143627 |

#### Table S15. Faith PD

| | | | | | | | |
|---|---|---|---|---|---|---|---|
| **Group-Level Effects:** | | | | | | | |
| **nest (Number of levels: 23)** | | | | | | | |
| | **Estimate** | **Est.Error** | **l-95% CI** | **u-95% CI** | **Rhat** | **Bulk_ESS** | **Tail_ESS** |
| sd(BCI_Intercept) | 0.12 | 0.04 | 0.04 | 0.20 | 1.00 | 38682 | 31898 |
| sd(Haptoglobin_Intercept) | 0.05 | 0.03 | 0.00 | 0.12 | 1.00 | 38548 | 61800 |
| sd(FaithPD_Intercept) | 0.05 | 0.03 | 0.00 | 0.12 | 1.00 | 60992 | 88316 |
| sd(CORT_Intercept) | 0.04 | 0.02 | 0.00 | 0.09 | 1.00 | 47206 | 77017 |
| **nest:ring_number (Number of levels: 43)** | | | | | | | |
| | **Estimate** | **Est.Error** | **l-95% CI** | **u-95% CI** | **Rhat** | **Bulk_ESS** | **Tail_ESS** |
| sd(BCI_Intercept) | 0.06 | 0.04 | 0.00 | 0.14 | 1.00 | 29673 | 54217 |
| sd(Haptoglobin_Intercept) | 0.03 | 0.02 | 0.00 | 0.09 | 1.00 | 56438 | 92926 |
| sd(FaithPD_Intercept) | 0.04 | 0.03 | 0.00 | 0.11 | 1.00 | 63496 | 88641 |
| sd(CORT_Intercept) | 0.03 | 0.02 | 0.00 | 0.08 | 1.00 | 46184 | 90112 |
| **Population-Level Effects:** | | | | | | | |
| | **Estimate** | **Est.Error** | **l-95% CI** | **u-95% CI** | **Rhat** | **Bulk_ESS** | **Tail_ESS** |
| BCI_Intercept | 0.70 | 0.12 | 0.46 | 0.94 | 1.00 | 160066 | 150464 |
| Haptoglobin_Intercept | 0.12 | 0.09 | -0.07 | 0.30 | 1.00 | 183369 | 150088 |
| FaithPD_Intercept | 0.48 | 0.12 | 0.25 | 0.70 | 1.00 | 183336 | 149744 |
| CORT_Intercept | 0.59 | 0.04 | 0.51 | 0.69 | 1.00 | 160318 | 156040 |
| BCI_FaithPD | 0.11 | 0.11 | -0.10 | 0.32 | 1.00 | 195841 | 156807 |
| BCI_CORT | -0.43 | 0.15 | -0.72 | -0.14 | 1.00 | 166981 | 154805 |
| BCI_Haptoglobin | 0.23 | 0.17 | -0.10 | 0.55 | 1.00 | 127842 | 133346 |
| BCI_Age | -0.11 | 0.12 | -0.34 | 0.14 | 1.00 | 136464 | 149465 |
| Haptoglobin_FaithPD | -0.02 | 0.09 | -0.19 | 0.15 | 1.00 | 230578 | 153645 |
| Haptoglobin_CORT | 0.03 | 0.12 | -0.20 | 0.26 | 1.00 | 185160 | 157469 |
| Haptoglobin_Age | 0.02 | 0.09 | -0.16 | 0.21 | 1.00 | 193232 | 157016 |
| FaithPD_CORT | 0.07 | 0.16 | -0.25 | 0.38 | 1.00 | 183577 | 158376 |
| FaithPD_Age | -0.16 | 0.13 | -0.42 | 0.09 | 1.00 | 195686 | 159848 |
| CORT_Age | -0.47 | 0.08 | -0.63 | -0.32 | 1.00 | 143907 | 154942 |

| | | | | | | | |
|---|---|---|---|---|---|---|---|
| **Family Specific Parameters:** | | | | | | | |
| | **Estimate** | **Est.Error** | **l-95% CI** | **u-95% CI** | **Rhat** | **Bulk_ESS** | **Tail_ESS** |
| sigma_BCI | 0.15 | 0.02 | 0.12 | 0.19 | 1.00 | 69276 | 114960 |
| sigma_Haptoglobin | 0.13 | 0.01 | 0.11 | 0.16 | 1.00 | 112819 | 138462 |
| sigma_FaithPD | 0.19 | 0.02 | 0.16 | 0.22 | 1.00 | 172827 | 151782 |
| sigma_CORT | 0.14 | 0.01 | 0.11 | 0.17 | 1.00 | 99573 | 138620 |
| alpha_CORT | 4.51 | 2.71 | -0.57 | 10.29 | 1.00 | 110645 | 132494 |

#### Table S16. N° of Observed ASV's

| **Group-Level Effects:** | | | | | | | |
|---|---|---|---|---|---|---|---|
| **nest (Number of levels: 23)** | | | | | | | |
| | **Estimate** | **Est.Error** | **l-95% CI** | **u-95% CI** | **Rhat** | **Bulk_ESS** | **Tail_ESS** |
| sd(BCI_Intercept) | 0.12 | 0.04 | 0.04 | 0.20 | 1.00 | 49217 | 43984 |
| sd(Haptoglobin_Intercept) | 0.06 | 0.03 | 0.00 | 0.12 | 1.00 | 43823 | 78048 |
| sd(ASV_Intercept) | 0.04 | 0.03 | 0.00 | 0.10 | 1.00 | 73534 | 108245 |
| sd(CORT_Intercept) | 0.04 | 0.02 | 0.00 | 0.09 | 1.00 | 55507 | 95386 |
| **nest:ring_number (Number of levels: 43)** | | | | | | | |
| | **Estimate** | **Est.Error** | **l-95% CI** | **u-95% CI** | **Rhat** | **Bulk_ESS** | **Tail_ESS** |
| sd(BCI_Intercept) | 0.06 | 0.04 | 0.00 | 0.14 | 1.00 | 36515 | 64074 |
| sd(Haptoglobin_Intercept) | 0.03 | 0.02 | 0.00 | 0.09 | 1.00 | 64382 | 105246 |
| sd(ASV_Intercept) | 0.03 | 0.02 | 0.00 | 0.08 | 1.00 | 98168 | 110066 |
| sd(CORT_Intercept) | 0.03 | 0.02 | 0.00 | 0.08 | 1.00 | 58612 | 101913 |
| **Population-Level Effects:** | | | | | | | |
| | **Estimate** | **Est.Error** | **l-95% CI** | **u-95% CI** | **Rhat** | **Bulk_ESS** | **Tail_ESS** |
| BCI_Intercept | 0.69 | 0.12 | 0.45 | 0.92 | 1.00 | 249579 | 154242 |
| Haptoglobin_Intercept | 0.10 | 0.09 | -0.09 | 0.28 | 1.00 | 296794 | 155887 |
| ASV_Intercept | 0.40 | 0.10 | 0.20 | 0.60 | 1.00 | 318155 | 154810 |
| CORT_Intercept | 0.59 | 0.04 | 0.51 | 0.69 | 1.00 | 224297 | 172718 |
| BCI_ASV | 0.17 | 0.12 | -0.06 | 0.40 | 1.00 | 320964 | 155417 |
| BCI_CORT | -0.43 | 0.15 | -0.72 | -0.14 | 1.00 | 257536 | 161108 |
| BCI_Haptoglobin | 0.22 | 0.16 | -0.11 | 0.54 | 1.00 | 170007 | 151221 |
| BCI_Age | -0.09 | 0.12 | -0.33 | 0.15 | 1.00 | 202081 | 155126 |
| Haptoglobin_ASV | 0.02 | 0.10 | -0.17 | 0.22 | 1.00 | 336553 | 157222 |

| **Group-Level Effects:** | | | | | | | |
|---|---|---|---|---|---|---|---|
| Haptoglobin_CORT | 0.02 | 0.12 | -0.21 | 0.25 | 1.00 | 285478 | 164665 |
| Haptoglobin_Age | 0.03 | 0.10 | -0.16 | 0.22 | 1.00 | 297820 | 167261 |
| ASV_CORT | 0.06 | 0.14 | -0.22 | 0.33 | 1.00 | 289168 | 164095 |
| ASV_Age | -0.17 | 0.11 | -0.40 | 0.05 | 1.00 | 304712 | 167802 |
| CORT_Age | -0.47 | 0.08 | -0.63 | -0.32 | 1.00 | 198304 | 169954 |
| **Family Specific Parameters:** | | | | | | | |
| | **Estimate** | **Est.Error** | **l-95% CI** | **u-95% CI** | **Rhat** | **Bulk_ESS** | **Tail_ESS** |
| sigma_BCI | 0.15 | 0.02 | 0.12 | 0.19 | 1.00 | 80124 | 128574 |
| sigma_Haptoglobin | 0.13 | 0.01 | 0.11 | 0.16 | 1.00 | 130042 | 145658 |
| sigma_ASV | 0.17 | 0.01 | 0.14 | 0.20 | 1.00 | 279694 | 157778 |
| sigma_CORT | 0.14 | 0.01 | 0.11 | 0.17 | 1.00 | 129980 | 148975 |
| alpha_CORT | 4.51 | 2.73 | -0.59 | 10.29 | 1.00 | 146675 | 154517 |

#### Table S17. Bayes R2 for each diversity measurement

| | **R2m** | **Est.Error** | **Q2.5** | **Q97.5** | **R2c** | **Est.Error** | **Q2.5** | **Q97.5** |
|---|---|---|---|---|---|---|---|---|
| **BCI (Path1)** | 0.18 | 0.06 | 0.07 | 0.29 | 0.52 | 0.08 | 0.33 | 0.66 |
| **Haptoglobin (Path2)** | 0.05 | 0.03 | 0.00 | 0.13 | 0.24 | 0.10 | 0.06 | 0.43 |
| **Shannon (Path3)** | 0.03 | 0.03 | 0.00 | 0.11 | 0.12 | 0.07 | 0.02 | 0.28 |
| **CORT (Path4)** | 0.36 | 0.08 | 0.20 | 0.52 | 0.47 | 0.11 | 0.25 | 0.66 |
| | **R2m** | **Est.Error** | **Q2.5** | **Q97.5** | **R2c** | **Est.Error** | **Q2.5** | **Q97.5** |
| **BCI (Path1)** | 0.16 | 0.06 | 0.05 | 0.27 | 0.50 | 0.09 | 0.30 | 0.65 |
| **Haptoglobin (Path2)** | 0.04 | 0.03 | 0.00 | 0.11 | 0.22 | 0.10 | 0.05 | 0.41 |
| **Faith PD (Path3)** | 0.08 | 0.05 | 0.01 | 0.20 | 0.19 | 0.09 | 0.05 | 0.37 |
| **CORT (Path4)** | 0.36 | 0.08 | 0.20 | 0.52 | 0.47 | 0.11 | 0.25 | 0.66 |
| | **R2m** | **Est.Error** | **Q2.5** | **Q97.5** | **R2c** | **Est.Error** | **Q2.5** | **Q97.5** |
| **BCI (Path1)** | 0.16 | 0.06 | 0.06 | 0.28 | 0.50 | 0.09 | 0.31 | 0.65 |
| **Haptoglobin (Path2)** | 0.04 | 0.03 | 0.00 | 0.11 | 0.23 | 0.10 | 0.05 | 0.42 |
| **ASV (Path3)** | 0.10 | 0.06 | 0.01 | 0.22 | 0.18 | 0.08 | 0.05 | 0.34 |
| **CORT (Path4)** | 0.36 | 0.08 | 0.20 | 0.52 | 0.47 | 0.11 | 0.25 | 0.66 |

### 5. Bayesian Structural Equation Modeling (SEM) diagnostics of 28S rRNA diversity measurements - model results for Haptoglobin immune assay

#### Table S18. Shannon diversity index

| | | | | | | | |
|---|---|---|---|---|---|---|---|
| **Group-Level Effects:** | | | | | | | |
| **nest (Number of levels: 23)** | | | | | | | |
| | **Estimate** | **Est.Error** | **l-95% CI** | **u-95% CI** | **Rhat** | **Bulk_ESS** | **Tail_ESS** |
| sd(BCI_Intercept) | 0.13 | 0.04 | 0.05 | 0.22 | 1.00 | 42951 | 37948 |
| sd(Haptoglobin_Intercept) | 0.05 | 0.03 | 0.00 | 0.11 | 1.00 | 50319 | 76803 |
| sd(Shannon_Intercept) | 0.05 | 0.04 | 0.00 | 0.14 | 1.00 | 85975 | 99903 |
| sd(CORT_Intercept) | 0.05 | 0.03 | 0.00 | 0.12 | 1.00 | 43080 | 63799 |
| **nest:ring_number (Number of levels: 41)** | | | | | | | |
| | **Estimate** | **Est.Error** | **l-95% CI** | **u-95% CI** | **Rhat** | **Bulk_ESS** | **Tail_ESS** |
| sd(BCI_Intercept) | 0.05 | 0.04 | 0.00 | 0.13 | 1.00 | 42050 | 65893 |
| sd(Haptoglobin_Intercept) | 0.04 | 0.03 | 0.00 | 0.09 | 1.00 | 61510 | 96100 |
| sd(Shannon_Intercept) | 0.05 | 0.04 | 0.00 | 0.13 | 1.00 | 80886 | 90962 |
| sd(CORT_Intercept) | 0.04 | 0.03 | 0.00 | 0.09 | 1.00 | 49853 | 91266 |
| **Population-Level Effects:** | | | | | | | |
| | **Estimate** | **Est.Error** | **l-95% CI** | **u-95% CI** | **Rhat** | **Bulk_ESS** | **Tail_ESS** |
| BCI_Intercept | 0.79 | 0.14 | 0.52 | 1.05 | 1.00 | 177015 | 149832 |
| Haptoglobin_Intercept | 0.04 | 0.11 | -0.17 | 0.26 | 1.00 | 178238 | 149776 |
| shannon_Intercept | 0.77 | 0.15 | 0.47 | 1.07 | 1.00 | 193181 | 148698 |
| CORT_Intercept | 0.62 | 0.05 | 0.53 | 0.72 | 1.00 | 202412 | 169172 |
| BCI_Shannon | -0.09 | 0.09 | -0.27 | 0.09 | 1.00 | 256177 | 154358 |
| BCI_CORT | -0.40 | 0.16 | -0.71 | -0.09 | 1.00 | 166515 | 152065 |
| BCI_Haptoglobin | 0.34 | 0.17 | -0.01 | 0.67 | 1.00 | 131149 | 122600 |
| BCI_Age | -0.15 | 0.13 | -0.40 | 0.11 | 1.00 | 154134 | 145350 |
| Haptoglobin_Shannon | 0.08 | 0.08 | -0.07 | 0.23 | 1.00 | 266918 | 156618 |
| Haptoglobin_CORT | 0.03 | 0.13 | -0.22 | 0.28 | 1.00 | 185576 | 160109 |
| Haptoglobin_Age | 0.05 | 0.10 | -0.16 | 0.25 | 1.00 | 193838 | 161802 |
| Shannon_CORT | -0.07 | 0.21 | -0.48 | 0.34 | 1.00 | 196076 | 154250 |
| Shannon_Age | -0.26 | 0.17 | -0.60 | 0.07 | 1.00 | 202369 | 161208 |
| CORT_Age | -0.50 | 0.08 | -0.67 | -0.34 | 1.00 | 178286 | 162816 |
| **Family Specific Parameters:** | | | | | | | |

| | Estimate | Est.Error | l-95% CI | u-95% CI | Rhat | Bulk_ESS | Tail_ESS |
|---|---|---|---|---|---|---|---|
| sigma_BCI | 0.15 | 0.02 | 0.12 | 0.19 | 1.00 | 80726 | 110024 |
| sigma_Haptoglobin | 0.14 | 0.01 | 0.11 | 0.17 | 1.00 | 130548 | 148475 |
| sigma_Shannon | 0.24 | 0.02 | 0.19 | 0.28 | 1.00 | 221222 | 146208 |
| sigma_CORT | 0.14 | 0.02 | 0.11 | 0.17 | 1.00 | 107657 | 138376 |
| alpha_CORT | 3.12 | 2.72 | -1.74 | 8.98 | 1.00 | 95525 | 115777 |

#### Table S19. Faith PD

| | | | | | | | |
|---|---|---|---|---|---|---|---|
| **Group-Level Effects:** | | | | | | | |
| **nest (Number of levels: 23)** | | | | | | | |
| | **Estimate** | **Est.Error** | **l-95% CI** | **u-95% CI** | **Rhat** | **Bulk_ESS** | **Tail_ESS** |
| sd(BCI_Intercept) | 0.13 | 0.04 | 0.05 | 0.22 | 1.00 | 45351 | 40452 |
| sd(Haptoglobin_Intercept) | 0.05 | 0.03 | 0.00 | 0.11 | 1.00 | 51291 | 79207 |
| sd(FaithPD_Intercept) | 0.03 | 0.03 | 0.00 | 0.10 | 1.00 | 101230 | 101146 |
| sd(CORT_Intercept) | 0.05 | 0.03 | 0.00 | 0.12 | 1.00 | 41187 | 66489 |
| **nest:ring_number (Number of levels: 41)** | | | | | | | |
| | **Estimate** | **Est.Error** | **l-95% CI** | **u-95% CI** | **Rhat** | **Bulk_ESS** | **Tail_ESS** |
| sd(BCI_Intercept) | 0.05 | 0.03 | 0.00 | 0.13 | 1.00 | 42197 | 63897 |
| sd(Haptoglobin_Intercept) | 0.04 | 0.03 | 0.00 | 0.10 | 1.00 | 60837 | 92942 |
| sd(FaithPD_Intercept) | 0.04 | 0.03 | 0.00 | 0.10 | 1.00 | 95660 | 94232 |
| sd(CORT_Intercept) | 0.04 | 0.03 | 0.00 | 0.09 | 1.00 | 50249 | 93337 |
| **Population-Level Effects:** | | | | | | | |
| | **Estimate** | **Est.Error** | **l-95% CI** | **u-95% CI** | **Rhat** | **Bulk_ESS** | **Tail_ESS** |
| BCI_Intercept | 0.79 | 0.13 | 0.54 | 1.04 | 1.00 | 164304 | 146946 |
| Haptoglobin_Intercept | 0.09 | 0.10 | -0.12 | 0.29 | 1.00 | 180969 | 143777 |
| FaithPD_Intercept | 0.45 | 0.12 | 0.20 | 0.69 | 1.00 | 208782 | 149273 |
| CORT_Intercept | 0.62 | 0.05 | 0.53 | 0.72 | 1.00 | 208633 | 166641 |
| BCI_FaithPD | -0.16 | 0.11 | -0.38 | 0.05 | 1.00 | 257166 | 152653 |
| BCI_CORT | -0.41 | 0.16 | -0.72 | -0.10 | 1.00 | 171235 | 151359 |
| BCI_Haptoglobin | 0.33 | 0.17 | -0.02 | 0.66 | 1.00 | 137582 | 124258 |
| BCI_Age | -0.17 | 0.13 | -0.42 | 0.09 | 1.00 | 155648 | 144723 |
| Haptoglobin_FaithPD | 0.04 | 0.10 | -0.15 | 0.23 | 1.00 | 269140 | 154322 |
| Haptoglobin_CORT | 0.02 | 0.13 | -0.23 | 0.28 | 1.00 | 189700 | 156382 |

| | | | | | | | |
|---|---|---|---|---|---|---|---|
| Haptoglobin_Age | 0.04 | 0.11 | -0.17 | 0.25 | 1.00 | 198629 | 154843 |
| FaithPD_CORT | -0.07 | 0.17 | -0.40 | 0.26 | 1.00 | 210016 | 159100 |
| FaithPD_Age | -0.28 | 0.14 | -0.55 | -0.01 | 1.00 | 214852 | 156644 |
| CORT_Age | -0.50 | 0.08 | -0.67 | -0.34 | 1.00 | 177992 | 158833 |
| **Family Specific Parameters:** | | | | | | | |
| | **Estimate** | **Est.Error** | **l-95% CI** | **u-95% CI** | **Rhat** | **Bulk_ESS** | **Tail_ESS** |
| sigma_BCI | 0.15 | 0.02 | 0.12 | 0.19 | 1.00 | 81762 | 109599 |
| sigma_Haptoglobin | 0.14 | 0.01 | 0.11 | 0.17 | 1.00 | 129246 | 148539 |
| sigma_FaithPD | 0.19 | 0.02 | 0.16 | 0.23 | 1.00 | 266445 | 144974 |
| sigma_CORT | 0.14 | 0.02 | 0.11 | 0.17 | 1.00 | 109605 | 136275 |
| alpha_CORT | 3.13 | 2.71 | -1.74 | 8.96 | 1.00 | 97554 | 117030 |

#### Table S20. N° of observed ASV's

| | | | | | | | |
|---|---|---|---|---|---|---|---|
| **Group-Level Effects:** | | | | | | | |
| **~nest (Number of levels: 23)** | | | | | | | |
| | **Estimate** | **Est.Error** | **l-95% CI** | **u-95% CI** | **Rhat** | **Bulk_ESS** | **Tail_ESS** |
| sd(BCI_Intercept) | 0.13 | 0.04 | 0.05 | 0.22 | 1.00 | 48956 | 43678 |
| sd(Haptoglobin_Intercept) | 0.05 | 0.03 | 0.00 | 0.11 | 1.00 | 51356 | 83029 |
| sd(ASV_Intercept) | 0.03 | 0.02 | 0.00 | 0.09 | 1.00 | 121055 | 106693 |
| sd(CORT_Intercept) | 0.05 | 0.03 | 0.00 | 0.12 | 1.00 | 46631 | 76304 |
| **nest:ring_number (Number of levels: 41)** | | | | | | | |
| | **Estimate** | **Est.Error** | **l-95% CI** | **u-95% CI** | **Rhat** | **Bulk_ESS** | **Tail_ESS** |
| sd(BCI_Intercept) | 0.05 | 0.03 | 0.00 | 0.13 | 1.00 | 45517 | 68906 |
| sd(Haptoglobin_Intercept) | 0.04 | 0.03 | 0.00 | 0.10 | 1.00 | 62403 | 101415 |
| sd(ASV_Intercept) | 0.04 | 0.03 | 0.00 | 0.11 | 1.00 | 92477 | 104228 |
| sd(CORT_Intercept) | 0.04 | 0.03 | 0.00 | 0.09 | 1.00 | 52226 | 93595 |
| **Population-Level Effects:** | | | | | | | |
| | **Estimate** | **Est.Error** | **l-95% CI** | **u-95% CI** | **Rhat** | **Bulk_ESS** | **Tail_ESS** |
| BCI_Intercept | 0.80 | 0.13 | 0.54 | 1.05 | 1.00 | 207560 | 156305 |
| Haptoglobin_Intercept | 0.08 | 0.11 | -0.13 | 0.29 | 1.00 | 225466 | 154892 |
| ASV_Intercept | 0.52 | 0.12 | 0.27 | 0.77 | 1.00 | 259536 | 150683 |
| CORT_Intercept | 0.62 | 0.05 | 0.53 | 0.72 | 1.00 | 247458 | 171992 |

| | | | | | | | |
|---|---|---|---|---|---|---|---|
| BCI_ASV | -0.15 | 0.11 | -0.36 | 0.06 | 1.00 | 326297 | 160195 |
| BCI_CORT | -0.42 | 0.16 | -0.72 | -0.10 | 1.00 | 206788 | 157745 |
| BCI_Haptoglobin | 0.34 | 0.17 | -0.01 | 0.67 | 1.00 | 149067 | 124855 |
| BCI_Age | -0.17 | 0.13 | -0.42 | 0.10 | 1.00 | 182753 | 148280 |
| Haptoglobin_ASV | 0.05 | 0.09 | -0.13 | 0.24 | 1.00 | 339562 | 157186 |
| Haptoglobin_CORT | 0.03 | 0.13 | -0.23 | 0.28 | 1.00 | 227703 | 164385 |
| Haptoglobin_Age | 0.04 | 0.11 | -0.17 | 0.25 | 1.00 | 236824 | 164963 |
| ASV_CORT | -0.11 | 0.17 | -0.44 | 0.23 | 1.00 | 250166 | 163417 |
| ASV_Age | -0.28 | 0.14 | -0.56 | -0.01 | 1.00 | 253690 | 160598 |
| CORT_Age | -0.50 | 0.08 | -0.67 | -0.34 | 1.00 | 211111 | 163889 |
| **Family Specific Parameters:** | | | | | | | |
| | **Estimate** | **Est.Error** | **l-95% CI** | **u-95% CI** | **Rhat** | **Bulk_ESS** | **Tail_ESS** |
| sigma_BCI | 0.15 | 0.02 | 0.12 | 0.19 | 1.00 | 85554 | 115319 |
| sigma_Haptoglobin | 0.14 | 0.01 | 0.11 | 0.17 | 1.00 | 135435 | 151746 |
| sigma_ASV | 0.19 | 0.02 | 0.16 | 0.23 | 1.00 | 292603 | 149156 |
| sigma_CORT | 0.14 | 0.02 | 0.11 | 0.17 | 1.00 | 120035 | 144255 |
| alpha_CORT | 3.12 | 2.72 | -1.74 | 8.94 | 1.00 | 108540 | 126059 |

#### Table S21. Bayes R2 for each diversity measurement

| | R2m | Est.Error | Q2.5 | Q97.5 | R2c | Est.Error | Q2.5 | Q97.5 |
|---|---|---|---|---|---|---|---|---|
| **BCI (Path1)** | 0.17 | 0.06 | 0.06 | 0.28 | 0.51 | 0.09 | 0.29 | 0.66 |
| **Haptoglobin (Path2)** | 0.06 | 0.04 | 0.01 | 0.16 | 0.23 | 0.10 | 0.05 | 0.43 |
| **Shannon (Path3)** | 0.08 | 0.05 | 0.00 | 0.20 | 0.16 | 0.08 | 0.03 | 0.33 |
| **CORT (Path4)** | 0.41 | 0.09 | 0.22 | 0.56 | 0.54 | 0.10 | 0.30 | 0.70 |
| | **R2m** | **Est.Error** | **Q2.5** | **Q97.5** | **R2c** | **Est.Error** | **Q2.5** | **Q97.5** |
| **BCI (Path1)** | 0.18 | 0.06 | 0.06 | 0.29 | 0.52 | 0.09 | 0.30 | 0.66 |
| **Haptoglobin (Path2)** | 0.05 | 0.03 | 0.00 | 0.13 | 0.22 | 0.10 | 0.05 | 0.43 |
| **Faith PD (Path3)** | 0.11 | 0.06 | 0.01 | 0.25 | 0.17 | 0.07 | 0.04 | 0.33 |
| **CORT (Path4)** | 0.41 | 0.09 | 0.22 | 0.56 | 0.54 | 0.10 | 0.30 | 0.70 |
| | **R2m** | **Est.Error** | **Q2.5** | **Q97.5** | **R2c** | **Est.Error** | **Q2.5** | **Q97.5** |
| **BCI (Path1)** | 0.17 | 0.06 | 0.06 | 0.29 | 0.52 | 0.09 | 0.30 | 0.67 |
| **Haptoglobin (Path2)** | 0.05 | 0.04 | 0.00 | 0.14 | 0.22 | 0.10 | 0.05 | 0.43 |
| **ASV (Path3)** | 0.10 | 0.06 | 0.01 | 0.24 | 0.16 | 0.08 | 0.04 | 0.33 |

| | R2m | Est.Error | Q2.5 | Q97.5 | R2c | Est.Error | Q2.5 | Q97.5 |
|---|---|---|---|---|---|---|---|---|
| **CORT (Path4)** | 0.41 | 0.09 | 0.22 | 0.56 | 0.54 | 0.10 | 0.30 | 0.70 |

### Figure S5. Differential abundance analysis results for each of the variables in study - ANCOM-BC2 model incorporating the latent variable *Immunity*

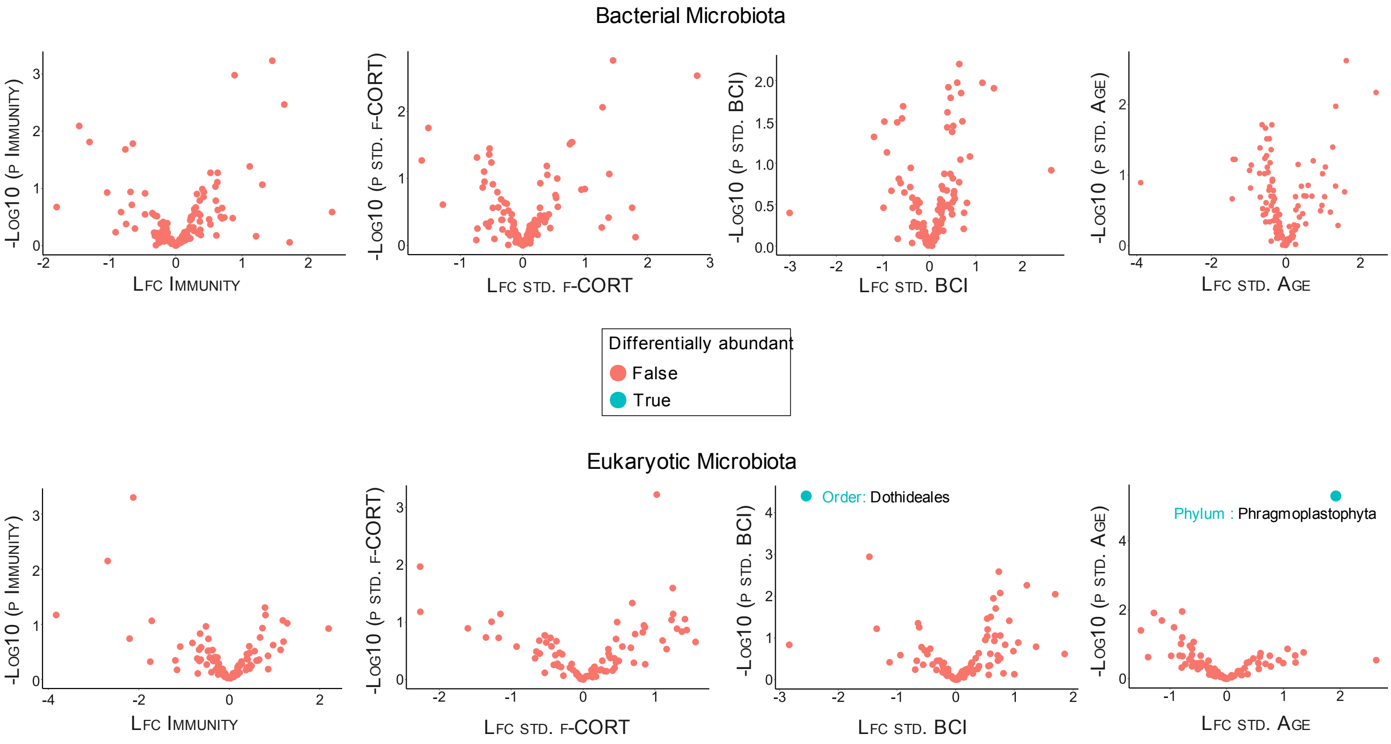

### Table S22. Sensitivity analysis for the two deferentially abundant taxa

| **Taxon** | **lfc (std.BCI)** | **se (std.BCI)** | **W (std.BCI)** | **p (std.BCI)** | **q (std.BCI)** | **diff (std.BCI)** | **passed_ss (std.BCI)** |
|---|---|---|---|---|---|---|---|
| d__Eukaryota;p_Ascomycota; c_Dothideomycetes;o_Dothideales; f_Dothideales;g_Dothideales; s_Hormonema_carpetanum | -2.60 | 0.18 | -14.12 | 0.00 | 0.01 | TRUE | TRUE |
| **Taxon** | **lfc (std.Age)** | **se (std.Age)** | **W (std.Age)** | **p (std.Age)** | **q (std.Age)** | **diff (std.Age)** | **passed_ss (std.Age)** |
| d__Eukaryota;p_Phragmoplastophyta; c_Phragmoplastophyta;o_Phragmoplastophyta; f_Phragmoplastophyta; g_Phragmoplastophyta;s_Pinus_taeda | 1.90 | 0.23 | 8.27 | 0.00 | 0.00 | TRUE | FALSE |

### Figure S6. Differential abundance analysis results for each of the variables in study - ANCOM-BC2 model incorporating Haptoglobin immune assay

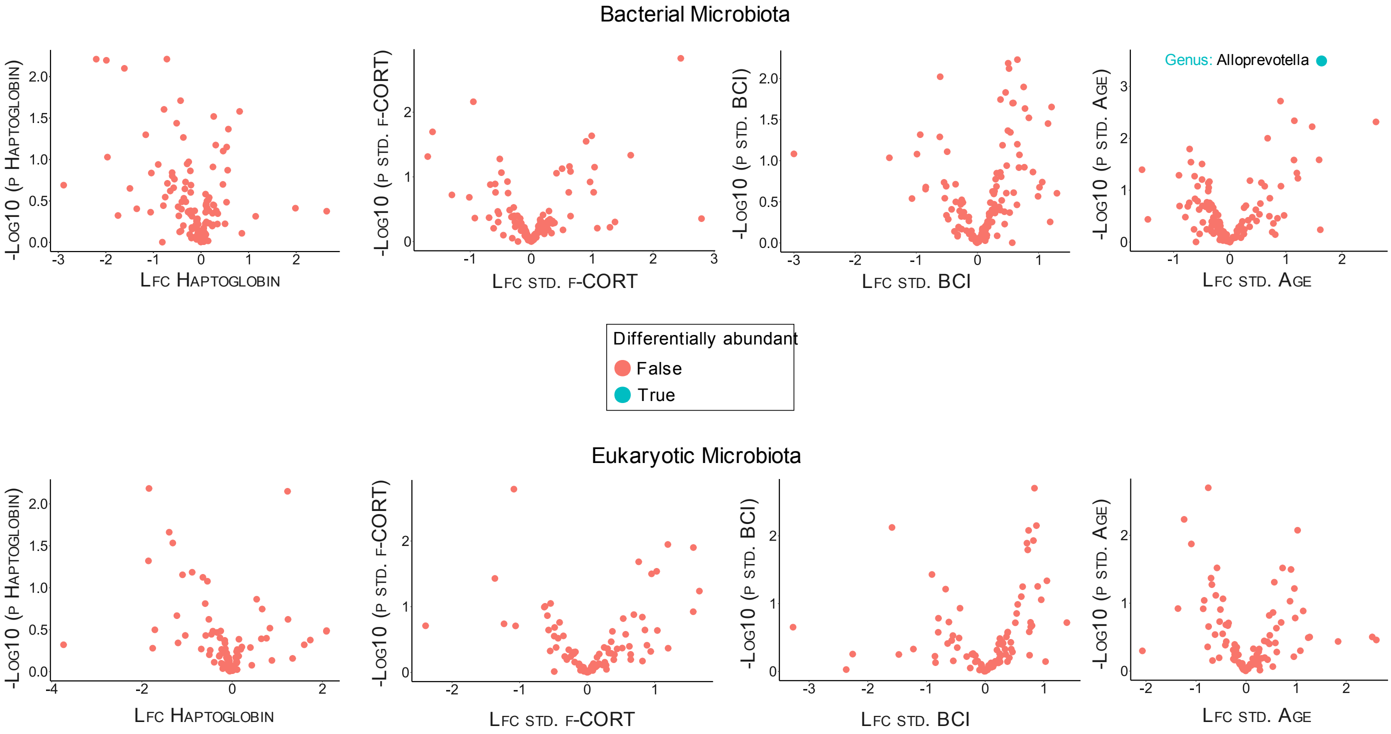

### Table S23. Sensitivity analysis for the deferentially abundant taxa

| Taxon | lfc (std.Age) | se (std.Age) | W (std.Age) | p (std.Age) | q (std.Age) | diff (std.Age) | passed_ss (std.Age) |
|---|---|---|---|---|---|---|---|
| d__Bacteria;p_Bacteroidota; c_Bacteroidia;o_Bacteroidales; f_Prevotellaceae;g_Alloprevotella;s_Alloprevotella_rava | 1.60 | 0.18 | 8.92 | 0.00 | 0.05 | TRUE | FALSE |

### Figure S7. Bayesian structural equation models for the different bacterial diversity measures with results from each immune assay superimposed onto each diagram

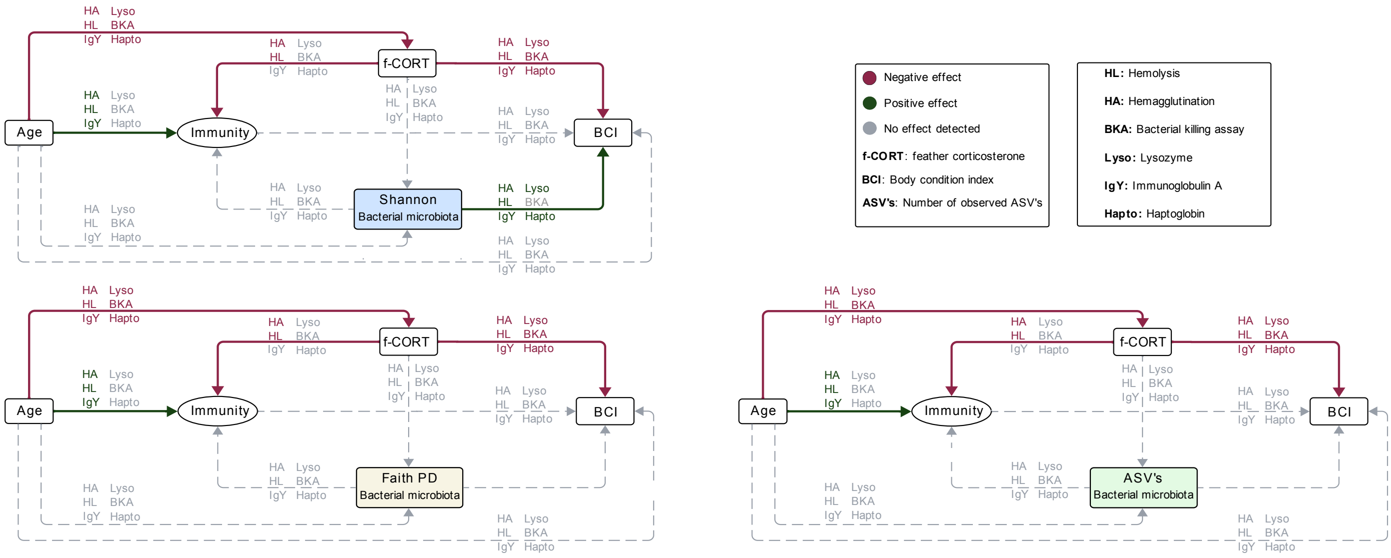

### Figure S8. Bayesian structural equation models for the different eukaryotic diversity measures with results from each immune assay superimposed onto each diagram

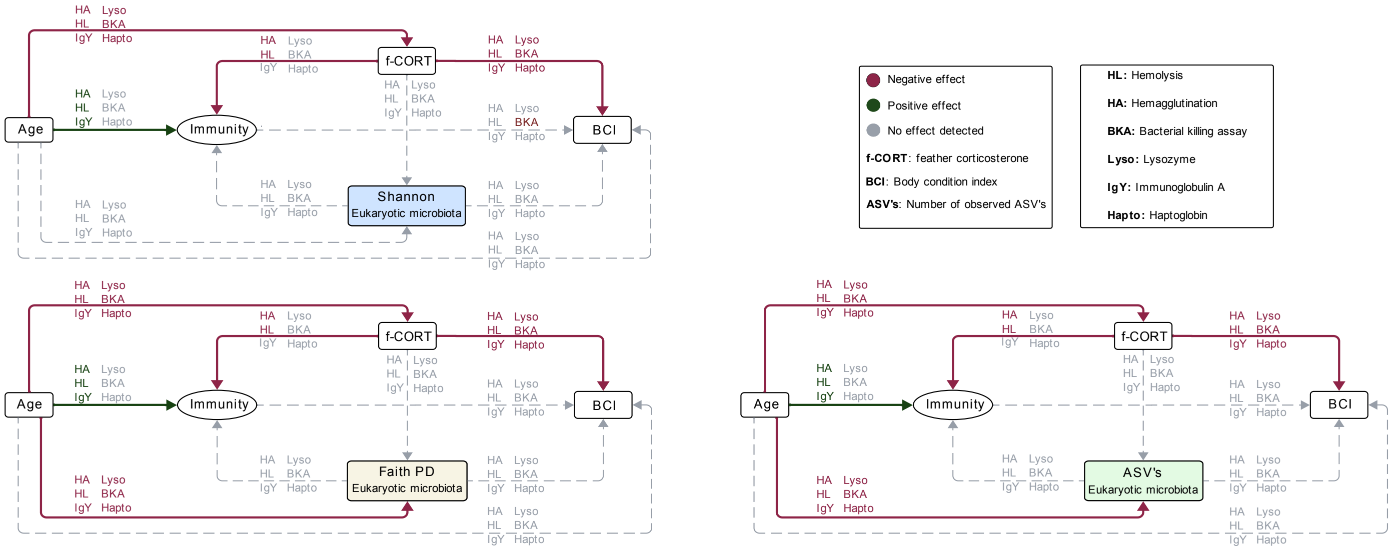
