## Supplementary material for "The gut microbiota-immune-brain axis in a wild vertebrate: dynamic interactions and health impacts": Associations-sex-habitat-rank

### Testing for associations of the variables of interest with : sex, habitat and rank

#### Table of Contents

**Testing for associations of the variables of interest with : sex, habitat and rank**
- Table of Contents
- 1. Read in the data
- 2. Body condition index
  - 2.1 Sex
  - 2.2 Habitat
  - 2.3 Rank
  - 2.4 Adjust p values
- 3. Corticosterone (CORT)
  - 3.1 Sex
  - 3.2 Habitat
  - 3.3 Rank
  - 3.4 Adjust p values
- 4. Hemagglutination
  - 4.1 Sex
  - 4.2 Habitat
  - 4.3 Rank
  - 4.4 Adjust p values
- 5. Hemolysis
  - 5.1 Sex
  - 5.2 Habitat
  - 5.3 Rank
  - 5.4 Adjust p values
- 6. Bacteria killing assay
  - 6.1 Sex
  - 6.2 Habitat
  - 6.3 Rank
  - 6.4 Adjust p values
- 7. Lysozyme
  - 7.1 Sex
  - 7.2 Habitat
  - 7.3 Rank
  - 7.4 Adjust p values
- 8. Immunoglobulin Y
  - 8.1 Sex
  - 8.2 Habitat
  - 8.3 Rank
  - 8.4 Adjust p values
- 9. Associations between the variables of interest and: sex; habitat and rank
  - 9.1 Body condition index (BCI)
  - 9.2 Feather corticosterone (f-CORT)
  - 9.3 Immune assays

### 1. Read in the data

```
#Load libraries
library(tidyverse)
library(lme4)
library(MuMIn)
library (performance)
library(car)
library(effects)
library(ggplot2)
library(openxlsx)
library(multcomp)

#Load the dataset
metadata <- readRDS("metadata_immune.rds")

#Scale all variables
metadata$std_ha <- as.numeric (scale(metadata$ha))
metadata$std_hl <- as.numeric(scale(metadata$hl))
metadata$std_bka <- as.numeric(scale(metadata$bka))
metadata$std_lyso <-as.numeric(scale(metadata$lyso))
metadata$std_igy <- as.numeric(scale(metadata$igy))
metadata$std_hapto <- as.numeric(scale(metadata$hapto))
metadata$std_shannon <- as.numeric(scale(metadata$shannon_entropy))
```

### 2. Body condition index

#### 2.1 Sex

```
# Fit LMM - Sex
bci_sex <- lmer(std_bci_two ~ sex + (1|nest/ring_number), data = metadata)

check_normality(bci_sex)
check_model(bci_sex)

# Model summary
summary(bci_sex)

# Test of significance
p1_bci <- Anova(bci_sex)
p1_bci

# Create a box plot
bci_sex_plot <- ggplot(metadata, aes(x = sex, y = std_bci_two, fill = sex)) +
  geom_boxplot() +
  geom_jitter(position = position_jitter(0.2), alpha = 0.5) +
  labs(x = "Sex", y = "Std BCI") +
  theme_classic()+
  scale_x_discrete(labels = c("F" = "Female", "M" = "Male")) +
  theme(axis.text.x = element_text(size = 16),  # Adjust the size as needed
        axis.text.y = element_text(size = 16))+
  theme(axis.title.x = element_text(size = 16),  # Adjust the size as needed
        axis.title.y = element_text(size = 16))+
  theme(axis.title.x = element_text(margin = margin(t = 13)))+
  theme(axis.title.y = element_text(margin = margin(r = 14)))+
  theme(legend.position = "none")+
```

```
  theme(text = element_text(family = "Arial"))
```

#### 2.2 Habitat

```
# Fit LMM - habitat
bci_habitat <- lmer(std_bci_two ~ habitat + (1|nest/ring_number), data = metadata)

check_normality(bci_habitat)
check_model(bci_habitat)

# Model summary
summary(bci_habitat)

# Test of significance
p3_bci <- Anova(bci_habitat)
p3_bci

# Create a box plot
bci_habitat_plot <- ggplot(metadata, aes(x = habitat, y = std_bci_two, fill = habitat)) +
  geom_boxplot() +
  geom_jitter(position = position_jitter(0.2), alpha = 0.5) +
  #stat_summary(fun.data="mean_sdl", mult=1, geom="crossbar", width=0.03) +
  labs(x = "Habitat", y = "Std BCI") +
  theme_classic()+
  scale_x_discrete(labels = c("north" = "North", "south" = "South")) +
  theme(axis.text.x = element_text(size = 16),  # Adjust the size as needed
        axis.text.y = element_text(size = 16))+
  theme(axis.title.x = element_text(size = 16),  # Adjust the size as needed
        axis.title.y = element_text(size = 16))+
  theme(axis.title.x = element_text(margin = margin(t = 13)))+
  theme(axis.title.y = element_text(margin = margin(r = 14)))+
  scale_fill_manual(values = c("#8BC34A", "#FF5722")) +
  theme(legend.position = "none")+
  theme(text = element_text(family = "Arial"))
```

#### 2.3 Rank

```
# Fit LMM - rank

bci_rank <- lmer(std_bci_two ~ rank + (1|nest/ring_number), data = metadata)

check_normality(bci_rank)
check_model(bci_rank)

# Model summary

summary(bci_rank)

# Test of significance

p4_bci <- Anova(bci_rank)
p4_bci

# Create a box plot
```

```
bci_rank_plot <- ggplot(metadata, aes(x = rank, y = std_bci_two, fill = rank)) +
  geom_boxplot() +
  geom_jitter(position = position_jitter(0.2), alpha = 0.5) +
  labs(x = "Rank", y = "Std BCI") +
  theme_classic()+
  theme(axis.text.x = element_text(size = 16),  # Adjust the size as needed
        axis.text.y = element_text(size = 16))+
  theme(axis.title.x = element_text(size = 16),  # Adjust the size as needed
        axis.title.y = element_text(size = 16))+
  theme(axis.title.x = element_text(margin = margin(t = 13)))+
  theme(axis.title.y = element_text(margin = margin(r = 14)))+
  scale_fill_manual(values = c("#FFC107", "#FF5733","#FF0000")) +
  theme(legend.position = "none")+
  theme(text = element_text(family = "Arial"))
```

#### 2.4 Adjust p values

```
#Adjust p values BH correction
p_values_bci <- c(p1_bci$`Pr(>Chisq)`,
                    p2_bci$`Pr(>Chisq)`,
                    p3_bci$`Pr(>Chisq)`,
                    p4_bci$`Pr(>Chisq))
adjusted_p_values_bci <- p.adjust(p_values_bci, method = "BH")
adjusted_p_values_bci <- format(round(adjusted_p_values_bci, digits = 3), scientific = FALSE)
adjusted_p_values_bci

#       sex     lbinom    habitat     rank
[1]   "0.367"  "0.421"   "0.348"    "0.207"
```

### 3. Corticosterone (CORT)

#### 3.1 Sex

```
# Fit LMM - Sex
cort_sex <- lmer(std_cort ~ sex + (1|nest/ring_number), data = metadata)

check_normality(cort_sex)
check_model(cort_sex)

# Model summary
summary(cort_sex)

# Test of significance
p1_cort <- Anova(cort_sex)
p1_cort

# Create a box plot
cort_sex_plot <- ggplot(metadata, aes(x = sex, y = std_cort, fill = sex)) +
  geom_boxplot() +
  geom_jitter(position = position_jitter(0.2), alpha = 0.5) +
  labs(x = "Sex", y = "Std CORT") +
```

```
  theme_classic()+
  scale_x_discrete(labels = c("F" = "Female", "M" = "Male")) +
  theme(axis.text.x = element_text(size = 16),  # Adjust the size as needed
        axis.text.y = element_text(size = 16))+
  theme(axis.title.x = element_text(size = 16),  # Adjust the size as needed
        axis.title.y = element_text(size = 16))+
  theme(axis.title.x = element_text(margin = margin(t = 13)))+
  theme(axis.title.y = element_text(margin = margin(r = 14)))+
  theme(legend.position = "none")+
  theme(text = element_text(family = "Arial"))
```

#### 3.2 Habitat

```
# Fit LMM - habitat
cort_habitat <- lmer(std_cort ~ habitat + (1|nest/ring_number), data = metadata)

check_normality(cort_habitat)
check_model(cort_habitat)

# Model summary
summary(cort_habitat)

# Test of significance
p3_cort <- Anova(cort_habitat)
p3_cort

# Create a box plot
cort_habitat_plot <- ggplot(metadata, aes(x = habitat, y = std_cort, fill = habitat)) +
  geom_boxplot() +
  geom_jitter(position = position_jitter(0.2), alpha = 0.5) +
  #stat_summary(fun.data="mean_sdl", mult=1, geom="crossbar", width=0.03) +
  labs(x = "Habitat", y = "Std CORT") +
  theme_classic()+
  scale_x_discrete(labels = c("north" = "North", "south" = "South")) +
  theme(axis.text.x = element_text(size = 16),  # Adjust the size as needed
        axis.text.y = element_text(size = 16))+
  theme(axis.title.x = element_text(size = 16),  # Adjust the size as needed
        axis.title.y = element_text(size = 16))+
  theme(axis.title.x = element_text(margin = margin(t = 13)))+
  theme(axis.title.y = element_text(margin = margin(r = 14)))+
  scale_fill_manual(values = c("#8BC34A", "#FF5722")) +
  theme(legend.position = "none")+
  theme(text = element_text(family = "Arial"))
```

#### 3.3 Rank

```
# Fit LMM - rank
cort_rank <- lmer(std_cort ~ rank + (1|nest/ring_number), data = metadata)

check_normality(cort_rank)
check_model(cort_rank)

# Model summary
summary(cort_rank)
```

```r
# Test of significance
p4_cort <- Anova(cort_rank)
p4_cort

# Create a box plot
cort_rank_plot <- ggplot(metadata, aes(x = rank, y = std_cort, fill = rank)) +
  geom_boxplot() +
  geom_jitter(position = position_jitter(0.2), alpha = 0.5) +
  labs(x = "Rank", y = "Std CORT") +
  theme_classic()+
  theme(axis.text.x = element_text(size = 16),  # Adjust the size as needed
        axis.text.y = element_text(size = 16))+
  theme(axis.title.x = element_text(size = 16),  # Adjust the size as needed
        axis.title.y = element_text(size = 16))+
  theme(axis.title.x = element_text(margin = margin(t = 13)))+
  theme(axis.title.y = element_text(margin = margin(r = 14)))+
  scale_fill_manual(values = c("#FFC107", "#FF5733","#FF0000")) +
  theme(legend.position = "none")+
  theme(text = element_text(family = "Arial"))
```

#### 3.4 Adjust p values

```r
#Adjust p values BH correction
p_values_cort <- c(p1_cort$`Pr(>Chisq)`,
                    p2_cort$`Pr(>Chisq)`,
                    p3_cort$`Pr(>Chisq)`,
                    p4_cort$`Pr(>Chisq))
adjusted_p_values_cort <- p.adjust(p_values_cort, method = "BH")
adjusted_p_values_cort <- format(round(adjusted_p_values_cort, digits = 3), scientific = FALSE)
adjusted_p_values_cort

#       sex      lbinom    habitat     rank
[1]    "0.767"  "0.371"    "0.951"    "0.951"
```

### 4. Hemagglutination

#### 4.1 Sex

```r
# Fit LMM - Sex
ha_sex <- lmer(std_ha ~ sex + (1|nest/ring_number), data = metadata)

check_normality(ha_sex)
check_model(ha_sex)

# Model summary
summary(ha_sex)

# Test of significance
p1_ha <- Anova(ha_sex)
p1_ha
```

```r
# Create a box plot
HA_sex_plot <- ggplot(metadata, aes(x = sex, y = std_ha, fill = sex)) +
  geom_boxplot() +
  geom_jitter(position = position_jitter(0.2), alpha = 0.5) +
  labs(x = "Sex", y = "Std Hemagglutination") +
  theme_classic()+
  scale_x_discrete(labels = c("F" = "Female", "M" = "Male")) +
  theme(axis.text.x = element_text(size = 16),  # Adjust the size as needed
        axis.text.y = element_text(size = 16))+
  theme(axis.title.x = element_text(size = 16),  # Adjust the size as needed
        axis.title.y = element_text(size = 16))+
  theme(axis.title.x = element_text(margin = margin(t = 13)))+
  theme(axis.title.y = element_text(margin = margin(r = 14)))+
  theme(legend.position = "none")+
  theme(text = element_text(family = "Arial"))
```

#### 4.2 Habitat

```r
# Fit LMM - habitat
ha_habitat <- lmer(std_ha ~ habitat + (1|nest/ring_number), data = metadata)

check_normality(ha_habitat)
check_model(ha_habitat)

# Model summary
summary(ha_habitat)

# Test of significance
p3_ha <- Anova(ha_habitat)
p3_ha

# Create a box plot
HA_habitat_plot <- ggplot(metadata, aes(x = habitat, y = std_ha, fill = habitat)) +
  geom_boxplot() +
  geom_jitter(position = position_jitter(0.2), alpha = 0.5) +
  #stat_summary(fun.data="mean_sdl", mult=1, geom="crossbar", width=0.03) +
  labs(x = "Habitat", y = "Std Hemagglutination") +
  theme_classic()+
  scale_x_discrete(labels = c("north" = "North", "south" = "South")) +
  theme(axis.text.x = element_text(size = 16),  # Adjust the size as needed
        axis.text.y = element_text(size = 16))+
  theme(axis.title.x = element_text(size = 16),  # Adjust the size as needed
        axis.title.y = element_text(size = 16))+
  theme(axis.title.x = element_text(margin = margin(t = 13)))+
  theme(axis.title.y = element_text(margin = margin(r = 14)))+
  scale_fill_manual(values = c("#8BC34A", "#FF5722")) +
  theme(legend.position = "none")+
  theme(text = element_text(family = "Arial"))
```

#### 4.3 Rank

```
# Fit LMM - rank
ha_rank <- lmer(std_ha ~ rank + (1|nest/ring_number), data = metadata)

check_normality(ha_rank)
check_model(ha_rank)

# Model summary
summary(ha_rank)

# Test of significance
p4_ha <- Anova(ha_rank)
p4_ha

# Create a box plot
HA_rank_plot <- ggplot(metadata, aes(x = rank, y = std_ha, fill = rank)) +
  geom_boxplot() +
  geom_jitter(position = position_jitter(0.2), alpha = 0.5) +
  labs(x = "Rank", y = "Std Hemagglutination") +
  theme_classic()+
  theme(axis.text.x = element_text(size = 16),  # Adjust the size as needed
        axis.text.y = element_text(size = 16))+
  theme(axis.title.x = element_text(size = 16),  # Adjust the size as needed
        axis.title.y = element_text(size = 16))+
  theme(axis.title.x = element_text(margin = margin(t = 13)))+
  theme(axis.title.y = element_text(margin = margin(r = 14)))+
  scale_fill_manual(values = c("#FFC107", "#FF5733","#FF0000")) +
  theme(legend.position = "none")+
  theme(text = element_text(family = "Arial"))
```

#### 4.4 Adjust p values

```
#Adjust p values BH correction
p_values_ha <- c(p1_ha$`Pr(>Chisq)`,
                    p2_ha$`Pr(>Chisq)`,
                    p3_ha$`Pr(>Chisq)`,
                    p4_ha$`Pr(>Chisq))
adjusted_p_values_ha <- p.adjust(p_values_ha, method = "BH")
adjusted_p_values_ha <- format(round(adjusted_p_values_ha, digits = 3), scientific = FALSE)
adjusted_p_values_ha

#       sex      lbinom    habitat     rank
[1]    "0.128"   "0.128"   "0.384"    "0.560"
```

### 5. Hemolysis

#### 5.1 Sex

```
# Fit LMM - Sex
hl_sex <- lmer(std_hl ~ sex + (1|nest/ring_number), data = metadata)

check_normality(hl_sex)
check_model(hl_sex)

# Model summary
summary(hl_sex)

# Test of significance
p1_hl <- Anova(hl_sex)
p1_hl

# Create a box plot
HL_sex_plot <- ggplot(metadata, aes(x = sex, y = std_hl, fill = sex)) +
  geom_boxplot() +
  geom_jitter(position = position_jitter(0.2), alpha = 0.5) +
  labs(x = "Sex", y = "Std Hemolysis") +
  theme_classic()+
  scale_x_discrete(labels = c("F" = "Female", "M" = "Male")) +
  theme(axis.text.x = element_text(size = 16),  # Adjust the size as needed
        axis.text.y = element_text(size = 16))+
  theme(axis.title.x = element_text(size = 16),  # Adjust the size as needed
        axis.title.y = element_text(size = 16))+
  theme(axis.title.x = element_text(margin = margin(t = 13)))+
  theme(axis.title.y = element_text(margin = margin(r = 14)))+
  theme(legend.position = "none")+
  theme(text = element_text(family = "Arial"))
```

#### 5.2 Habitat

```
# Fit LMM - habitat
hl_habitat <- lmer(std_hl ~ habitat + (1|nest/ring_number), data = metadata)

check_normality(hl_habitat)
check_model(hl_habitat)

# Model summary
summary(hl_habitat)

# Test of significance
p3_hl <- Anova(hl_habitat)
p3_hl

# Create a box plot
HL_habitat_plot <- ggplot(metadata, aes(x = habitat, y = std_hl, fill = habitat)) +
  geom_boxplot() +
  geom_jitter(position = position_jitter(0.2), alpha = 0.5) +
  labs(x = "Habitat", y = "Std Hemolysis") +
  theme_classic()+
  scale_x_discrete(labels = c("north" = "North", "south" = "South")) +
  theme(axis.text.x = element_text(size = 16),  # Adjust the size as needed
        axis.text.y = element_text(size = 16))+
  theme(axis.title.x = element_text(size = 16),  # Adjust the size as needed
```

```
        axis.title.y = element_text(size = 16))+
  theme(axis.title.x = element_text(margin = margin(t = 13)))+
  theme(axis.title.y = element_text(margin = margin(r = 14)))+
  scale_fill_manual(values = c("#8BC34A", "#FF5722")) +
  theme(legend.position = "none")+
  theme(text = element_text(family = "Arial"))
```

#### 5.3 Rank

```
# Fit LMM - rank
hl_rank <- lmer(std_hl ~ rank + (1|nest/ring_number), data = metadata)

check_normality(hl_rank)
check_model(hl_rank)

# Model summary
summary(hl_rank)

# Test of significance
p4_hl <- Anova(hl_rank)
p4_hl

# Create a box plot
HL_rank_plot <- ggplot(metadata, aes(x = rank, y = std_hl, fill = rank)) +
  geom_boxplot() +
  geom_jitter(position = position_jitter(0.2), alpha = 0.5) +
  labs(x = "Rank", y = "Std Hemolysis") +
  theme_classic()+
  theme(axis.text.x = element_text(size = 16),  # Adjust the size as needed
        axis.text.y = element_text(size = 16))+
  theme(axis.title.x = element_text(size = 16),  # Adjust the size as needed
        axis.title.y = element_text(size = 16))+
  theme(axis.title.x = element_text(margin = margin(t = 13)))+
  theme(axis.title.y = element_text(margin = margin(r = 14)))+
  scale_fill_manual(values = c("#FFC107", "#FF5733","#FF0000")) +
  theme(legend.position = "none")+
  theme(text = element_text(family = "Arial"))
```

#### 5.4 Adjust p values

```
#Adjust p values BH correction
p_values_hl <- c(p1_hl$`Pr(>Chisq)`,
                    p2_hl$`Pr(>Chisq)`,
                    p3_hl$`Pr(>Chisq)`,
                    p4_hl$`Pr(>Chisq)`)
adjusted_p_values_hl <- p.adjust(p_values_hl, method = "BH")
adjusted_p_values_hl <- format(round(adjusted_p_values_hl, digits = 3), scientific = FALSE)
adjusted_p_values_hl

#       sex      lbinom    habitat     rank
[1]    "0.260"   "0.428"   "0.713"   "0.260"
```

### 6. Bacteria killing assay

#### 6.1 Sex

```
# Fit LMM - Sex
bka_sex <- lmer(std_bka ~ sex + (1|nest/ring_number), data = metadata)

check_normality(bka_sex)
check_model(bka_sex)

# Model summary
summary(bka_sex)

# Test of significance
p1_bka <- Anova(bka_sex)
p1_bka

# Create a box plot
BKA_sex_plot <- ggplot(metadata, aes(x = sex, y = std_bka, fill = sex)) +
  geom_boxplot() +
  geom_jitter(position = position_jitter(0.2), alpha = 0.5) +
  labs(x = "Sex", y = "Std % E.coli killed") +
  theme_classic()+
  scale_x_discrete(labels = c("F" = "Female", "M" = "Male")) +
  theme(axis.text.x = element_text(size = 16),  # Adjust the size as needed
        axis.text.y = element_text(size = 16))+
  theme(axis.title.x = element_text(size = 16),  # Adjust the size as needed
        axis.title.y = element_text(size = 16))+
  theme(axis.title.x = element_text(margin = margin(t = 13)))+
  theme(axis.title.y = element_text(margin = margin(r = 14)))+
  theme(legend.position = "none")+
  theme(text = element_text(family = "Arial"))
```

#### 6.2 Habitat

```
# Fit LMM - habitat
bka_habitat <- lmer(std_bka ~ habitat + (1|nest/ring_number), data = metadata)

check_normality(bka_habitat)
check_model(bka_habitat)

# Model summary
summary(bka_habitat)

# Test of significance
p3_bka <- Anova(bka_habitat)
p3_bka

# Create a box plot
BKA_habitat_plot <- ggplot(metadata, aes(x = habitat, y = std_bka, fill = habitat)) +
  geom_boxplot() +
  geom_jitter(position = position_jitter(0.2), alpha = 0.5) +
```

```
  labs(x = "Habitat", y = "Std % E.coli killed") +
  theme_classic()+
  scale_x_discrete(labels = c("north" = "North", "south" = "South")) +
  theme(axis.text.x = element_text(size = 16),  # Adjust the size as needed
        axis.text.y = element_text(size = 16))+
  theme(axis.title.x = element_text(size = 16),  # Adjust the size as needed
        axis.title.y = element_text(size = 16))+
  theme(axis.title.x = element_text(margin = margin(t = 13)))+
  theme(axis.title.y = element_text(margin = margin(r = 14)))+
  scale_fill_manual(values = c("#8BC34A", "#FF5722")) +
  theme(legend.position = "none")+
  theme(text = element_text(family = "Arial"))
```

#### 6.3 Rank

```
# Fit LMM - rank
bka_rank <- lmer(std_bka ~ rank + (1|nest/ring_number), data = metadata)

check_normality(bka_rank)
check_model(bka_rank)

# Model summary
summary(bka_rank)

# Test of significance
p4_bka <- Anova(bka_rank)
p4_bka

# Create a box plot
BKA_rank_plot <- ggplot(metadata, aes(x = rank, y = std_bka, fill = rank)) +
  geom_boxplot() +
  geom_jitter(position = position_jitter(0.2), alpha = 0.5) +
  labs(x = "Rank", y = "Std % E.coli killed") +
  theme_classic()+
  theme(axis.text.x = element_text(size = 16),  # Adjust the size as needed
        axis.text.y = element_text(size = 16))+
  theme(axis.title.x = element_text(size = 16),  # Adjust the size as needed
        axis.title.y = element_text(size = 16))+
  theme(axis.title.x = element_text(margin = margin(t = 13)))+
  theme(axis.title.y = element_text(margin = margin(r = 14)))+
  scale_fill_manual(values = c("#FFC107", "#FF5733","#FF0000")) +
  theme(legend.position = "none")+
  theme(text = element_text(family = "Arial"))
```

#### 6.4 Adjust p values

```
#Adjust p values BH correction
p_values_bka <- c(p1_bka$`Pr(>Chisq)`,
                  p2_bka$`Pr(>Chisq)`,
                  p3_bka$`Pr(>Chisq)`,
                  p4_bka$`Pr(>Chisq)`)
adjusted_p_values_bka <- p.adjust(p_values_bka, method = "BH")
adjusted_p_values_bka <- format(round(adjusted_p_values_bka, digits = 3), scientific = FALSE)
adjusted_p_values_bka

#       sex      lbinom    habitat     rank
[1]    "0.930"   "0.600"   "0.798"    "0.600"
```

### 7. Lysozyme

#### 7.1 Sex

```
# Fit LMM - Sex
lyso_sex <- lmer(std_lyso ~ sex + (1|nest/ring_number), data = metadata)

check_normality(lyso_sex)
check_model(lyso_sex)

# Model summary
summary(lyso_sex)

# Test of significance
p1_lyso <- Anova(lyso_sex)
p1_lyso

# Create a box plot
lyso_sex_plot <- ggplot(metadata, aes(x = sex, y = std_lyso, fill = sex)) +
  geom_boxplot() +
  geom_jitter(position = position_jitter(0.2), alpha = 0.5) +
  labs(x = "Sex", y = "Std Lysozyme (mg/ml)") +
  theme_classic()+
  scale_x_discrete(labels = c("F" = "Female", "M" = "Male")) +
  theme(axis.text.x = element_text(size = 16),  # Adjust the size as needed
        axis.text.y = element_text(size = 16))+
  theme(axis.title.x = element_text(size = 16),  # Adjust the size as needed
        axis.title.y = element_text(size = 16))+
  theme(axis.title.x = element_text(margin = margin(t = 13)))+
  theme(axis.title.y = element_text(margin = margin(r = 14)))+
  theme(legend.position = "none")+
  theme(text = element_text(family = "Arial"))
```

#### 7.2 Habitat

```
# Fit LMM - habitat
lyso_habitat <- lmer(std_lyso ~ habitat + (1|nest/ring_number), data = metadata)

check_normality(lyso_habitat)
check_model(lyso_habitat)

# Model summary
summary(lyso_habitat)

# Test of significance
p3_lyso <- Anova(lyso_habitat)
p3_lyso

# Create a box plot
lyso_habitat_plot <- ggplot(metadata, aes(x = habitat, y = std_lyso, fill = habitat)) +
  geom_boxplot() +
  geom_jitter(position = position_jitter(0.2), alpha = 0.5) +
  labs(x = "Habitat", y = "Std Lysozyme (mg/ml)") +
  theme_classic()+
  scale_x_discrete(labels = c("north" = "North", "south" = "South")) +
  theme(axis.text.x = element_text(size = 16),  # Adjust the size as needed
        axis.text.y = element_text(size = 16))+
  theme(axis.title.x = element_text(size = 16),  # Adjust the size as needed
        axis.title.y = element_text(size = 16))+
  theme(axis.title.x = element_text(margin = margin(t = 13)))+
  theme(axis.title.y = element_text(margin = margin(r = 14)))+
  scale_fill_manual(values = c("#8BC34A", "#FF5722")) +
  theme(legend.position = "none")+
  theme(text = element_text(family = "Arial"))
```

#### 7.3 Rank

```
# Fit LMM - rank
lyso_rank <- lmer(std_lyso ~ rank + (1|nest/ring_number), data = metadata)

check_normality(lyso_rank)
check_model(lyso_rank)

# Model summary
summary(lyso_rank)

# Test of significance
p4_lyso <- Anova(lyso_rank)
p4_lyso

# Create a box plot
lyso_rank_plot <- ggplot(metadata, aes(x = rank, y = std_lyso, fill = rank)) +
  geom_boxplot() +
  geom_jitter(position = position_jitter(0.2), alpha = 0.5) +
  labs(x = "Rank", y = "Std Lysozyme (mg/ml)") +
  theme_classic()+
  theme(axis.text.x = element_text(size = 16),  # Adjust the size as needed
        axis.text.y = element_text(size = 16))+
  theme(axis.title.x = element_text(size = 16),  # Adjust the size as needed
```

```
        axis.title.y = element_text(size = 16))+
  theme(axis.title.x = element_text(margin = margin(t = 13)))+
  theme(axis.title.y = element_text(margin = margin(r = 14)))+
  scale_fill_manual(values = c("#FFC107", "#FF5733","#FF0000")) +
  theme(legend.position = "none")+
  theme(text = element_text(family = "Arial"))
```

#### 7.4 Adjust p values

```
#Adjust p values BH correction
p_values_lyso <- c(p1_lyso$`Pr(>Chisq)`,
                   p2_lyso$`Pr(>Chisq)`,
                   p3_lyso$`Pr(>Chisq)`,
                   p4_lyso$`Pr(>Chisq)`)
adjusted_p_values_lyso <- p.adjust(p_values_lyso, method = "BH")
adjusted_p_values_lyso <- format(round(adjusted_p_values_lyso, digits = 3), scientific = FALSE)
adjusted_p_values_lyso

#       sex      lbinom    habitat     rank
[1]   "0.520"   "0.149"   "0.002"    "0.520"
```

### 8. Immunoglobulin Y

#### 8.1 Sex

```
# Log transform igy
metadata$log_igy <- log(metadata$igy)

# Fit LMM - Sex
igy_sex <- lmer(log_igy ~ sex + (1|nest/ring_number), data = metadata)

check_normality(igy_sex)
check_model(igy_sex)

# Model summary
summary(igy_sex)

# Test of significance
p1_igy <- Anova(igy_sex)
p1_igy

# Create a box plot
igy_sex_plot <- ggplot(metadata, aes(x = sex, y = std_igy, fill = sex)) +
  geom_boxplot() +
  geom_jitter(position = position_jitter(0.2), alpha = 0.5) +
  labs(x = "Sex", y = "Log [IgY] (absorvance)") +
  theme_classic()+
  scale_x_discrete(labels = c("F" = "Female", "M" = "Male")) +
  theme(axis.text.x = element_text(size = 16),  # Adjust the size as needed
        axis.text.y = element_text(size = 16))+
  theme(axis.title.x = element_text(size = 16),  # Adjust the size as needed
```

```
        axis.title.y = element_text(size = 16))+
  theme(axis.title.x = element_text(margin = margin(t = 13)))+
  theme(axis.title.y = element_text(margin = margin(r = 14)))+
  theme(legend.position = "none")+
  theme(text = element_text(family = "Arial"))
```

#### 8.2 Habitat

```
# Fit LMM - habitat
igy_habitat <- lmer(std_igy ~ habitat + (1|nest/ring_number), data = metadata)

check_normality(igy_habitat)
check_model(igy_habitat)

# Model summary
summary(igy_habitat)

# Test of significance
p3_igy <- Anova(igy_habitat)
p3_igy

# Create a box plot
igy_habitat_plot <- ggplot(metadata, aes(x = habitat, y = std_igy, fill = habitat)) +
  geom_boxplot() +
  geom_jitter(position = position_jitter(0.2), alpha = 0.5) +
  #stat_summary(fun.data="mean_sdl", mult=1, geom="crossbar", width=0.03) +
  labs(x = "habitat", y = "Log [IgY] (absorvance)") +
  theme_classic()+
  scale_x_discrete(labels = c("north" = "North", "south" = "South")) +
  theme(axis.text.x = element_text(size = 16),  # Adjust the size as needed
        axis.text.y = element_text(size = 16))+
  theme(axis.title.x = element_text(size = 16),  # Adjust the size as needed
        axis.title.y = element_text(size = 16))+
  theme(axis.title.x = element_text(margin = margin(t = 13)))+
  theme(axis.title.y = element_text(margin = margin(r = 14)))+
  scale_fill_manual(values = c("#8BC34A", "#FF5722")) +
  theme(legend.position = "none")+
  theme(text = element_text(family = "Arial"))
```

#### 8.3 Rank

```
# Fit LMM - rank
igy_rank <- lmer(std_igy ~ rank + (1|nest/ring_number), data = metadata)

check_normality(igy_rank)
check_model(igy_rank)

# Model summary
summary(igy_rank)

# Test of significance
p4_igy <- Anova(igy_rank)
p4_igy
```

```r
# Create a box plot
igy_rank_plot <- ggplot(metadata, aes(x = rank, y = std_igy, fill = rank)) +
  geom_boxplot() +
  geom_jitter(position = position_jitter(0.2), alpha = 0.5) +
  labs(x = "Rank", y = "Log [IgY] (absorvance)") +
  theme_classic()+
  theme(axis.text.x = element_text(size = 16),  # Adjust the size as needed
        axis.text.y = element_text(size = 16))+
  theme(axis.title.x = element_text(size = 16),  # Adjust the size as needed
        axis.title.y = element_text(size = 16))+
  theme(axis.title.x = element_text(margin = margin(t = 13)))+
  theme(axis.title.y = element_text(margin = margin(r = 14)))+
  scale_fill_manual(values = c("#FFC107", "#FF5733","#FF0000")) +
  theme(legend.position = "none")+
  theme(text = element_text(family = "Arial"))
```

#### 8.4 Adjust p values

```r
#Adjust p values BH correction
p_values_igy <- c(p1_igy$`Pr(>Chisq)`,
                  p2_igy$`Pr(>Chisq)`,
                  p3_igy$`Pr(>Chisq)`,
                  p4_igy$`Pr(>Chisq)`)
adjusted_p_values_igy <- p.adjust(p_values_igy, method = "BH")
adjusted_p_values_igy <- format(round(adjusted_p_values_igy, digits = 3), scientific = FALSE)
adjusted_p_values_igy

#       sex     lbinom    habitat      rank
[1]  "0.111"   "0.002"    "0.584"   "0.950"
```

### 9. Associations between the variables of interest and: sex; habitat and rank

#### 9.1 Body condition index (BCI)

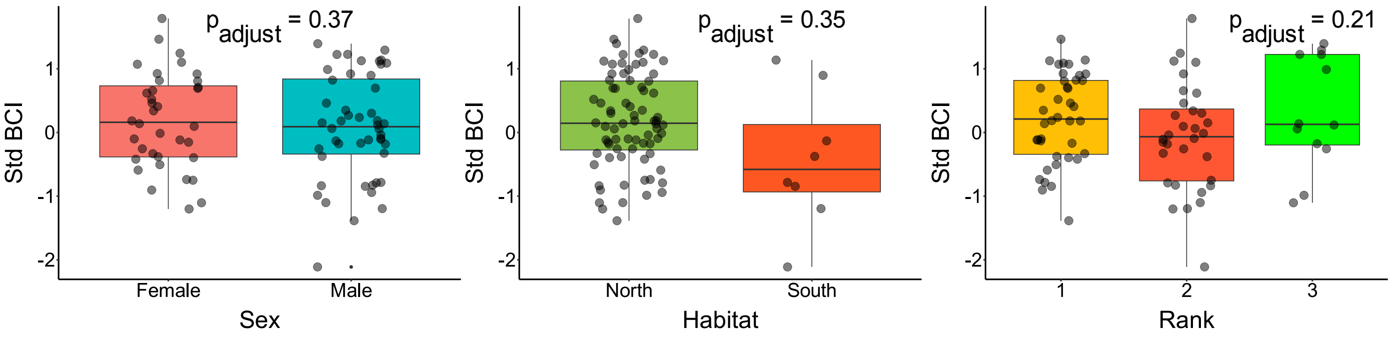

#### 9.2 Feather corticosterone (f-CORT)

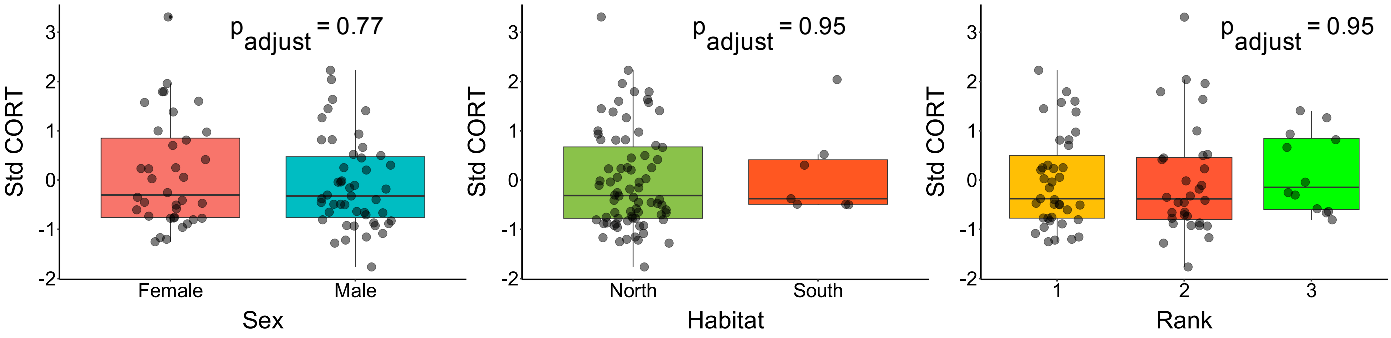


#### 9.3 Immune assays

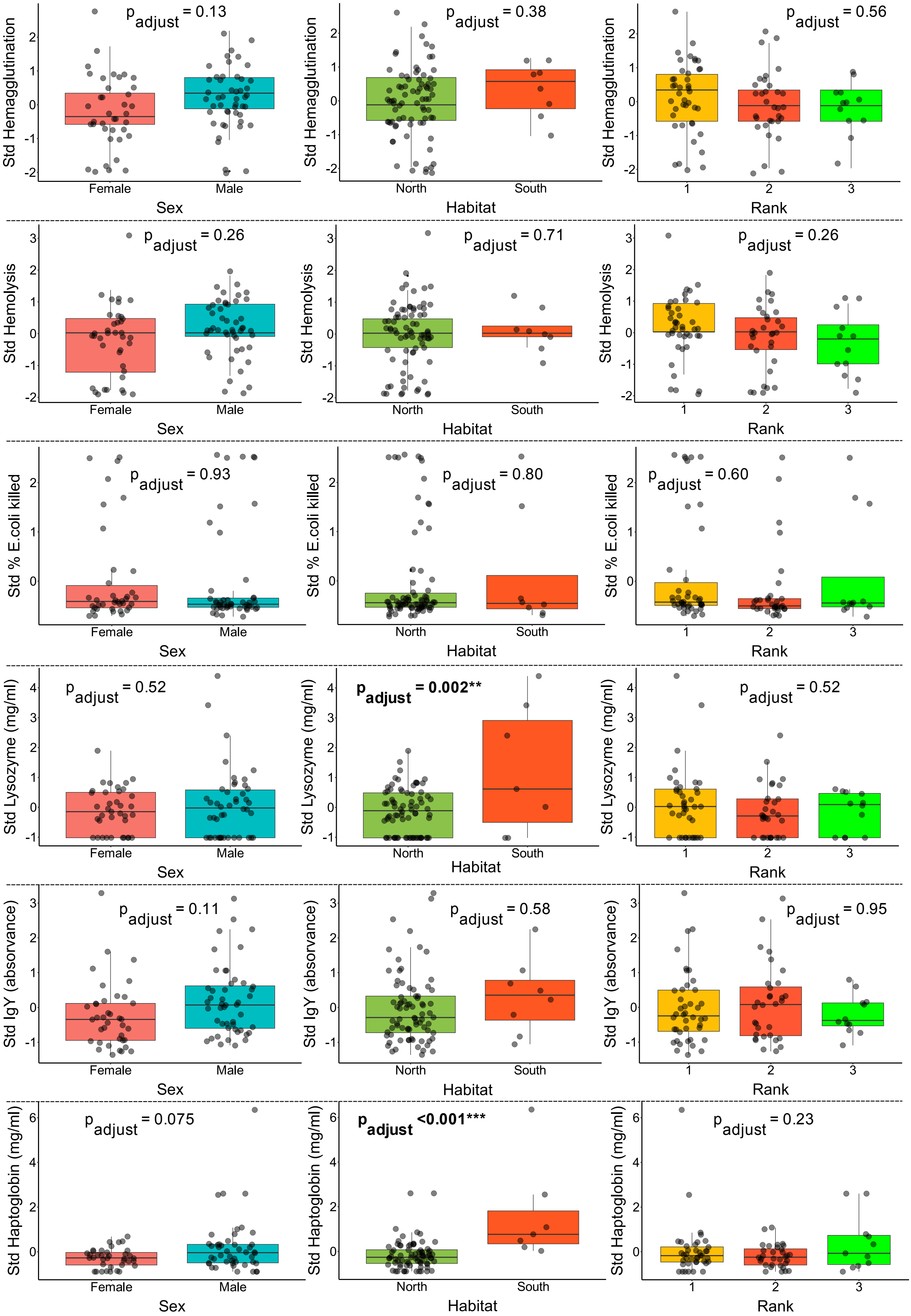

Std Hemagglutination
$p_{adjust}$ = 0.13
Female
Male
Sex
$p_{adjust}$ = 0.38
North
South
Habitat
$p_{adjust}$ = 0.56
1
2
3
Rank
Std Hemolysis
$p_{adjust}$ = 0.26
Female
Male
Sex
$p_{adjust}$ = 0.71
North
South
Habitat
$p_{adjust}$ = 0.26
1
2
3
Rank
Std % E.coli killed
$p_{adjust}$ = 0.93
Female
Male
Sex
$p_{adjust}$ = 0.80
North
South
Habitat
$p_{adjust}$ = 0.60
1
2
3
Rank
Std Lysozyme (mg/ml)
$p_{adjust}$ = 0.52
Female
Male
Sex
**$p_{adjust}$ = 0.002****
North
South
Habitat
$p_{adjust}$ = 0.52
1
2
3
Rank
Std IgY (absorvance)
$p_{adjust}$ = 0.11
Female
Male
Sex
$p_{adjust}$ = 0.58
North
South
Habitat
$p_{adjust}$ = 0.95
1
2
3
Rank
Std Haptoglobin (mg/ml)
$p_{adjust}$ = 0.075
Female
Male
Sex
**$p_{adjust}$ <0.001*****
North
South
Habitat
$p_{adjust}$ = 0.23
1
2
3
Rank
