## Supplementary material for "The gut microbiota-immune-brain axis in a wild vertebrate: dynamic interactions and health impacts": Bayesian-SEM-immunity

### Bayesian Structural Equation Modelling - constructing and modelling latent variable *"Immunity"*

#### Table of contents


**Bayesian Structural Equation Modelling - constructing and modelling latent variable *"Immunity"***

- Table of contents
- A) 16S rRNA (bacterial microbiota) SEM analysis
  - 1. Build a latent variable
  - 2. Define SEM for each diversity measurement
  - 3. Run brms
  - 4. Model Diagnostics Shannon
    - 4.1 Model Summary
    - 4.2 Model diagnostics
    - 4.3 Compare distribution of response variable to distributions of predicted response variable
    - 4.4 Plot model posterior and credible intervals
  - 5. Model diagnostics - Faith PD
    - 5.1 Model summary
    - 5.2 Model diagnostics
    - 5.3 Compare distribution of response variable to distributions of predicted response variable
    - 5.4 Plot model posterior and credible intervals
  - 6. Model diagnostics - N° of observed ASV's
    - 6.1 Model summary
    - 6.2 Model diagnostics
    - 6.3 Compare distribution of response variable to distributions of predicted response variable
    - 6.4 Plot model posterior and credible intervals
- B) 28S rRNA (eukaryotic microbiota) SEM analysis
  - 1. Build the latent variable
  - 2. Define SEM for each diversity measurement
  - 3. Run brms
  - 4. Model Diagnostics - Shannon
    - 4.1 Model Summary
    - 4.2 Model diagnostics
    - 4.3 Compare distribution of response variable to distributions of predicted response variable
    - 4.4 Plot model posterior and credible intervals
  - 5. Model diagnostics - Faith PD
    - 5.1 Model summary
    - 5.2 Model diagnostics
    - 5.3 Compare distribution of response variable to distributions of predicted response variable
    - 5.4 Plot model posterior and credible intervals
  - 6. Model diagnostics - N° of observed ASV's
    - 6.1 Model summary
    - 6.2 Model diagnostics
    - 6.3 Compare distribution of response variable to distributions of predicted response variable
    - 5.4 Plot model posterior and credible intervals

### A) 16S rRNA (bacterial microbiota) SEM analysis

#### 1. Build a latent variable

```
#Load Packages
library(brms)
library(rstan)
library(lavaan)
library(bayesplot)
library(bayestestR)
library(parallel)
library(svglite)
library(ggplot2)

#Load the data
metadata <- readRDS("16s_metadata_immune.rds")

#Scale immune assay scores
metadata$std_ha <- as.numeric (scale(metadata$ha))
metadata$std_hl <- as.numeric(scale(metadata$hl))
metadata$std_bka <- as.numeric(scale(metadata$bka))
metadata$std_lyso <-as.numeric(scale(metadata$lyso))
metadata$std_igy <- as.numeric(scale(metadata$igy))
metadata$std_hapto <- as.numeric(scale(metadata$hapto))

# Exploratory factor analysis - all imune assays included.

model_factor <- 'immunity =~  std_ha + std_hl + std_bka + std_lyso + std_igy + std_hapto'

fit_factor <-  efa(model_factor, data = metadata, cluster = c("ring_number"), missing = "fiml",
estimator = "MLR", std.lv=T)

#Model summary
summary(fit_factor, fit.measures = TRUE, standardized = TRUE, rsquare = TRUE)

lavaan 0.6.16 ended normally after 17 iterations

  Estimator                                         ML
  Optimization method                           NLMINB
  Number of model parameters                        18

  Number of observations                            86
  Number of clusters [ring_number]                  43
  Number of missing patterns                         4

Model Test User Model:
                                              Standard      Scaled
  Test Statistic                                42.002      56.933
  Degrees of freedom                                 9           9
  P-value (Chi-square)                           0.000       0.000
  Scaling correction factor                                  0.738
    Yuan-Bentler correction (Mplus variant)

Model Test Baseline Model:
```

```
  Test statistic                               199.481     251.107
  Degrees of freedom                                15          15
  P-value                                        0.000       0.000
  Scaling correction factor                                  0.794

User Model versus Baseline Model:

  Comparative Fit Index (CFI)                    0.821       0.797
  Tucker-Lewis Index (TLI)                       0.702       0.662

  Robust Comparative Fit Index (CFI)                         0.846
  Robust Tucker-Lewis Index (TLI)                            0.743

Loglikelihood and Information Criteria:

  Loglikelihood user model (H0)               -641.901    -641.901
  Scaling correction factor                                  1.674
      for the MLR correction
  Loglikelihood unrestricted model (H1)       -620.900    -620.900
  Scaling correction factor                                  1.362
      for the MLR correction

  Akaike (AIC)                                1319.802    1319.802
  Bayesian (BIC)                              1363.980    1363.980
  Sample-size adjusted Bayesian (SABIC)       1307.189    1307.189

Root Mean Square Error of Approximation:

  RMSEA                                          0.206       0.249
  90 Percent confidence interval - lower         0.146       0.180
  90 Percent confidence interval - upper         0.271       0.323
  P-value H_0: RMSEA <= 0.050                    0.000       0.000
  P-value H_0: RMSEA >= 0.080                    0.999       1.000

  Robust RMSEA                                               0.189
  90 Percent confidence interval - lower                     0.104
  90 Percent confidence interval - upper                     0.278
  P-value H_0: Robust RMSEA <= 0.050                         0.007
  P-value H_0: Robust RMSEA >= 0.080                         0.979

Standardized Root Mean Square Residual:

  SRMR                                           0.107       0.107

Parameter Estimates:

  Standard errors                         Robust.cluster
  Information                                   Observed
  Observed information based on                  Hessian

Latent Variables:
                   Estimate  Std.Err  z-value  P(>|z|)   Std.lv  Std.all
  immunity =~
    std_ha            0.929    0.091   10.196    0.000    0.929    0.934
    std_hl            0.905    0.098    9.196    0.000    0.905    0.910
    std_bka           0.309    0.112    2.768    0.006    0.309    0.311
    std_lyso          0.321    0.121    2.644    0.008    0.321    0.323
    std_igy           0.528    0.112    4.730    0.000    0.528    0.531
    std_hapto         0.256    0.146    1.759    0.079    0.256    0.257
```

```
Intercepts:
                   Estimate  Std.Err  z-value  P(>|z|)   Std.lv  Std.all
   .std_ha           -0.000    0.098   -0.000    1.000   -0.000   -0.000
   .std_hl           -0.000    0.111   -0.000    1.000   -0.000   -0.000
   .std_bka          -0.000    0.102   -0.000    1.000   -0.000   -0.000
   .std_lyso         -0.009    0.107   -0.080    0.936   -0.009   -0.009
   .std_igy          -0.000    0.114   -0.000    1.000   -0.000   -0.000
   .std_hapto        -0.010    0.111   -0.094    0.925   -0.010   -0.010
    immunity          0.000                               0.000    0.000

Variances:
                   Estimate  Std.Err  z-value  P(>|z|)   Std.lv  Std.all
   .std_ha            0.126    0.070    1.793    0.073    0.126    0.127
   .std_hl            0.170    0.074    2.311    0.021    0.170    0.172
   .std_bka           0.893    0.161    5.532    0.000    0.893    0.903
   .std_lyso          0.887    0.217    4.086    0.000    0.887    0.896
   .std_igy           0.710    0.116    6.108    0.000    0.710    0.718
   .std_hapto         0.924    0.440    2.100    0.036    0.924    0.934
    immunity          1.000                               1.000    1.000

R-Square:
                   Estimate
    std_ha            0.873
    std_hl            0.828
    std_bka           0.097
    std_lyso          0.104
    std_igy           0.282
    std_hapto         0.066

#Extract main model fit measures
fitMeasures(fit_factor, c("pvalue.scaled","cfi.robust","rmsea.robust","srmr"))

  pvalue.scaled    cfi.robust     rmsea.robust         srmr # not a good fit
      0.000           0.846           0.189            0.107



# Exploratory factor analysis excluding haptoglobin.

model_factor1 <- 'immunity =~  std_ha + std_hl + std_bka + std_lyso + std_igy'

fit_factor1 <-  efa(model_factor1, data = metadata, cluster = c("ring_number"), missing =
"fiml", estimator = "MLR", std.lv=T)

#Model summary
summary(fit_factor, fit.measures = TRUE, standardized = TRUE, rsquare = TRUE)

lavaan 0.6.16 ended normally after 17 iterations

  Estimator                                         ML
  Optimization method                           NLMINB
  Number of model parameters                        15

  Number of observations                            86
  Number of clusters [ring_number]                  43
  Number of missing patterns                         2

Model Test User Model:
                                              Standard      Scaled
  Test Statistic                                 8.638       7.338
```

```
  Degrees of freedom                                 5           5
  P-value (Chi-square)                           0.124       0.197
  Scaling correction factor                                  1.177
    Yuan-Bentler correction (Mplus variant)

Model Test Baseline Model:

  Test statistic                               161.049     151.364
  Degrees of freedom                                10          10
  P-value                                        0.000       0.000
  Scaling correction factor                                  1.064

User Model versus Baseline Model:

  Comparative Fit Index (CFI)                    0.976       0.983
  Tucker-Lewis Index (TLI)                       0.952       0.967

  Robust Comparative Fit Index (CFI)                         0.985
  Robust Tucker-Lewis Index (TLI)                            0.970

Loglikelihood and Information Criteria:

  Loglikelihood user model (H0)               -527.166    -527.166
  Scaling correction factor                                  1.201
      for the MLR correction
  Loglikelihood unrestricted model (H1)       -522.847    -522.847
  Scaling correction factor                                  1.195
      for the MLR correction

  Akaike (AIC)                                1084.332    1084.332
  Bayesian (BIC)                              1121.148    1121.148
  Sample-size adjusted Bayesian (SABIC)       1073.822    1073.822

Root Mean Square Error of Approximation:

  RMSEA                                          0.092       0.074
  90 Percent confidence interval - lower         0.000       0.000
  90 Percent confidence interval - upper         0.193       0.171
  P-value H_0: RMSEA <= 0.050                    0.212       0.300
  P-value H_0: RMSEA >= 0.080                    0.644       0.529

  Robust RMSEA                                               0.072
  90 Percent confidence interval - lower                     0.000
  90 Percent confidence interval - upper                     0.196
  P-value H_0: Robust RMSEA <= 0.050                         0.330
  P-value H_0: Robust RMSEA >= 0.080                         0.541

Standardized Root Mean Square Residual:

  SRMR                                           0.039       0.039

Parameter Estimates:

  Standard errors                           Robust.cluster
  Information                                     Observed
  Observed information based on                    Hessian

Latent Variables:
                   Estimate  Std.Err  z-value  P(>|z|)   Std.lv  Std.all
  immunity =~
```

```
    std_ha            0.936    0.094   10.007    0.000    0.936    0.942
    std_hl            0.898    0.103    8.758    0.000    0.898    0.904
    std_bka           0.305    0.118    2.576    0.010    0.305    0.307
    std_lyso          0.310    0.116    2.685    0.007    0.310    0.312
    std_igy           0.528    0.111    4.775    0.000    0.528    0.532

Intercepts:
                   Estimate  Std.Err  z-value  P(>|z|)   Std.lv  Std.all
   .std_ha           -0.000    0.098   -0.000    1.000   -0.000   -0.000
   .std_hl           -0.000    0.111   -0.000    1.000   -0.000   -0.000
   .std_bka           0.000    0.102    0.000    1.000    0.000    0.000
   .std_lyso         -0.008    0.107   -0.079    0.937   -0.008   -0.009
   .std_igy           0.000    0.114    0.000    1.000    0.000    0.000
    immunity          0.000                               0.000    0.000

Variances:
                   Estimate  Std.Err  z-value  P(>|z|)   Std.lv  Std.all
   .std_ha            0.112    0.089    1.261    0.207    0.112    0.113
   .std_hl            0.181    0.089    2.032    0.042    0.181    0.183
   .std_bka           0.895    0.164    5.456    0.000    0.895    0.906
   .std_lyso          0.894    0.223    4.010    0.000    0.894    0.903
   .std_igy           0.709    0.116    6.092    0.000    0.709    0.717
    immunity          1.000                               1.000    1.000

R-Square:
                   Estimate
    std_ha            0.887
    std_hl            0.817
    std_bka           0.094
    std_lyso          0.097
    std_igy           0.283


# Extract main model fit measures
fitMeasures(fit_factor1, c("pvalue.scaled","cfi.robust","rmsea.robust","srmr"))

pvalue.scaled      cfi.robust   rmsea.robust      srmr   # good fit
        0.197           0.985          0.072         0.039



# Predict and extract the values of the latent variable immunity.

metadata$pred_immunity <- lavPredict(fit_factor1, newdata = metadata, type = "lv", method =
"regression",
                                     transform = FALSE, se = "none", acov = "none",
                                     label = TRUE, fsm = FALSE,
                                     append.data = FALSE, assemble = FALSE,
                                     level = 1L, optim.method = "bfgs", ETA = NULL)

metadata$pred_immunity <- as.numeric(metadata$pred_immunity)
```

#### 2. Define SEM for each diversity measurement

```
#scale all predictors to range between 0-1 if they are not already naturally on that scale

#define scaling function:
```

```
range.use <- function(x,min.use,max.use){ (x - min(x,na.rm=T))/(max(x,na.rm=T)-min(x,na.rm=T))
* (max.use - min.use) + min.use }
scalecols<-c("bci_two", "shannon_entropy", "faith_pd", "observed_features", "cort",
"pred_immunity", "age_days")


for(i in 1:ncol(metadata[,which(colnames(metadata)%in%scalecols)])){
  metadata[,which(colnames(metadata)%in%scalecols)][,i]<-
range.use(metadata[,which(colnames(metadata)%in%scalecols)][,i],0,1)
}

# Define structural equation model paths

#Shannon
path1 <- bf(bci_two ~ shannon_entropy + cort + pred_immunity + age_days + (1|nest/ring_number))
path2 <- bf(pred_immunity ~  shannon_entropy + cort + age_days + (1|nest/ring_number))
path3 <- bf(shannon_entropy ~ cort + age_days + (1|nest/ring_number))
path4 <- bf(cort ~ age_days + (1|nest/ring_number)) + skew_normal()

sem_immunity_shannon <- path1 + path2 + path3 + path4

#Faith PD
path1 <- bf(bci_two ~ faith_pd + cort + pred_immunity + age_days + (1|nest/ring_number))
path2 <- bf(pred_immunity ~  faith_pd + cort + age_days + (1|nest/ring_number))
path3 <- bf(faith_pd ~ cort + age_days + (1|nest/ring_number))
path4 <- bf(cort ~ age_days + (1|nest/ring_number)) + skew_normal()

sem_immunity_faith <- path1 + path2 + path3 + path4

#N° of observed ASV's
path1 <- bf(bci_two ~ observed_features + cort + pred_immunity + age_days +
(1|nest/ring_number))
path2 <- bf(pred_immunity ~  observed_features + cort + age_days + (1|nest/ring_number))
path3 <- bf(observed_features ~ cort + age_days + (1|nest/ring_number))
path4 <- bf(cort ~ age_days + (1|nest/ring_number)) + skew_normal()

sem_immunity_asv <- path1 + path2 + path3 + path4
```

#### 3. Run brms

```
ncores = detectCores()
options(mc.cores = parallel::detectCores())

#Shannon
model_immunity_shannon <-brm(sem_immunity_shannon + set_rescor(FALSE),
                         data = metadata,
                         warmup = 50000, iter = 100000,
                         control = list(adapt_delta = 0.99, max_treedepth = 15),
                         cores=ncores, chains=4, init=1000)

#Faith PD
model_immunity_faith <-brm(sem_immunity_faith + set_rescor(FALSE),
                         data = metadata,
                         warmup = 50000, iter = 100000,
                         control = list(adapt_delta = 0.99, max_treedepth = 15),
                         cores=ncores, chains=4, init=1000)
```

```
#N° of observed ASV's
model_immunity_asv <-brm(sem_immunity_asv + set_rescor(FALSE),
                         data = metadata,
                         warmup = 50000, iter = 100000,
                         control = list(adapt_delta = 0.99, max_treedepth = 15),
                         cores=ncores, chains=4, init=1000)
```

### 4. Model Diagnostics Shannon

#### 4.1 Model Summary

```
#Model summary
summary_shannon<- summary(model_immunity_shannon)

#Bayes R2
R2m_shannon <- bayes_R2(model_immunity_shannon,re_formula=NA)
R2c_shannon <- bayes_R2(model_immunity_shannon)
```

#### 4.2 Model diagnostics

```
# Model diagnostics
diagnostic_shannon <- plot(model_immunity_shannon)

#Loop to save all diagnostic plots
diagnostic_plots <- list()
for (i in 1:length(diagnostic_shannon)) {
  diagnostic_plots[[i]] <- diagnostic_shannon[[i]]
  filename <- paste0("diagnostic", i, "_shannon")
  ggsave(filename = paste0(filename, ".png"), plot = diagnostic_plots[[i]], device = "png",
dpi=300)
}
```

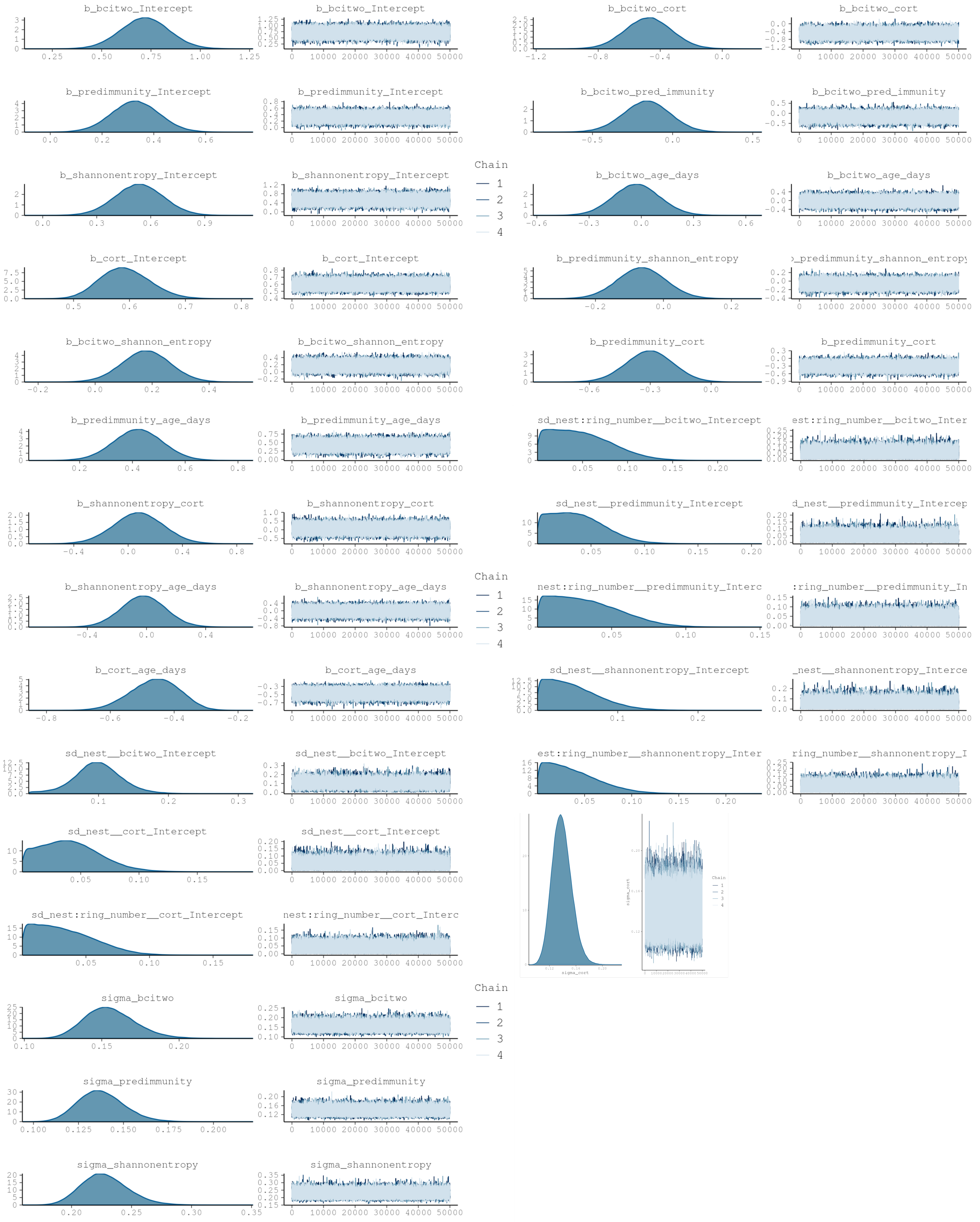


#### 4.3 Compare distribution of response variable to distributions of predicted response variable

```
#Posterior predictive checks (one by one)
distribution_shannon_path1 <- pp_check(model_immunity_shannon, resp="bcitwo", ndraws=200)
distribution_shannon_path2 <- pp_check(model_immunity_shannon, resp="predimmunity", ndraws=200)
distribution_shannon_path3 <- pp_check(model_immunity_shannon, resp="shannonentropy",
ndraws=200)
distribution_shannon_path4 <- pp_check(model_immunity_shannon, resp="cort", ndraws=200)
```

```
#Loop to save all distributions plot
responses <- c("bcitwo", "predimmunity", "shannonentropy", "cort")
response_names <- c("bci", "immune", "shannon", "cort")
for (i in seq_along(responses)) {
  pp_check_plot <- pp_check(model_immunity_shannon, resp = responses[i], ndraws = 200)
  filename <- paste0("distribution_shannon_", response_names[i])
  ggsave(filename = paste0(filename, ".svg"), plot = pp_check_plot, device = "svg", width = 8,
height = 10)
}
```

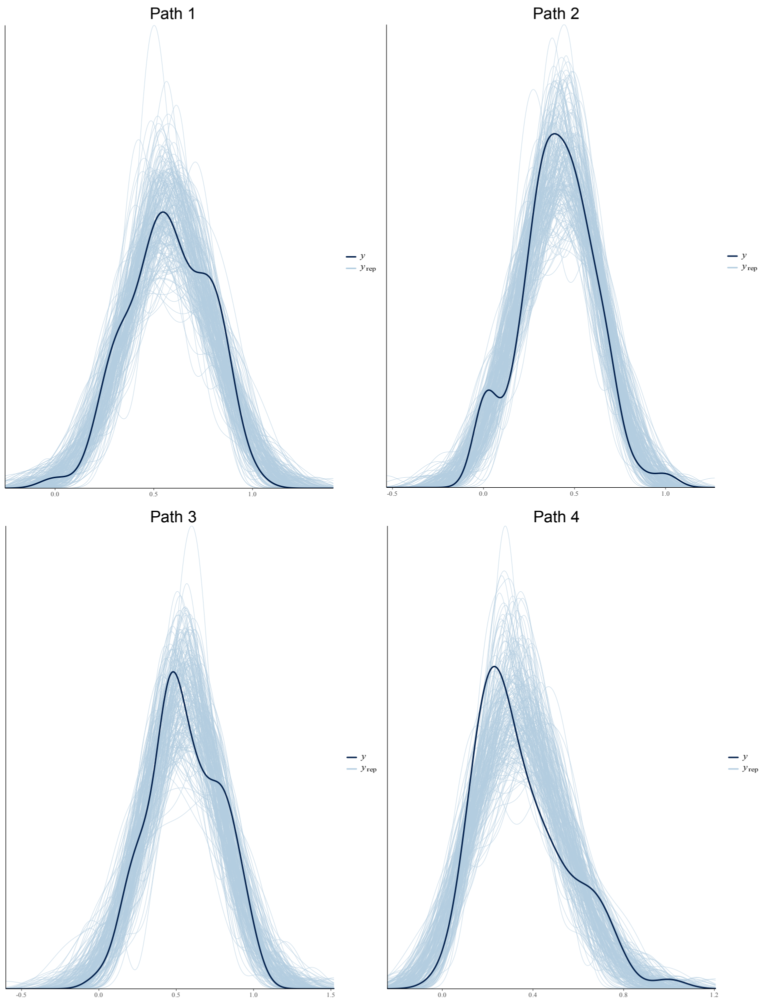


#### 4.4 Plot model posterior and credible intervals

```
#Plot Model effects
plot1 <-mcmc_plot(model_immunity_shannon, type = "intervals",prob_outer=0.95, prob=0.95,
                  variable =c("b_bcitwo_shannon_entropy", "b_bcitwo_cort",
"b_bcitwo_pred_immunity", "b_bcitwo_age_days",
                               "b_predimmunity_shannon_entropy", "b_predimmunity_cort",
"b_predimmunity_age_days",
                               "b_shannonentropy_cort", "b_shannonentropy_age_days",
                               "b_cort_age_days"))

plot1 <- plot1 + theme_classic() + geom_vline(xintercept = 0, linetype="dotted", color="blue")+
theme(axis.text.x = element_text(size = 16),  # Adjust the size as needed
axis.text.y = element_text(size = 16))+
theme(text = element_text(family = "Arial"))

ggsave(filename="16s_effect_sizes_shannon.svg", plot=plot1, device = "svg", width = 8, height =
10)
```

### 5. Model diagnostics - Faith PD

#### 5.1 Model summary

```
#Model summary
summary_faith<- summary(model_immunity_faith)

#Bayes R2
R2m_faith <- bayes_R2(model_immunity_faith,re_formula=NA)
R2c_faith <- bayes_R2(model_immunity_faith)
```

#### 5.2 Model diagnostics

```
# Model diagnostics
diagnostic_faith <- plot(model_immunity_faith)

#Loop to save all diagnostic plots
diagnostic_plots <- list()
for (i in 1:length(diagnostic_faith)) {
  diagnostic_plots[[i]] <- diagnostic_faith[[i]]
  filename <- paste0("diagnostic", i, "_faith")
  ggsave(filename = paste0(filename, ".png"), plot = diagnostic_plots[[i]], device = "png",
dpi=200)
}
```

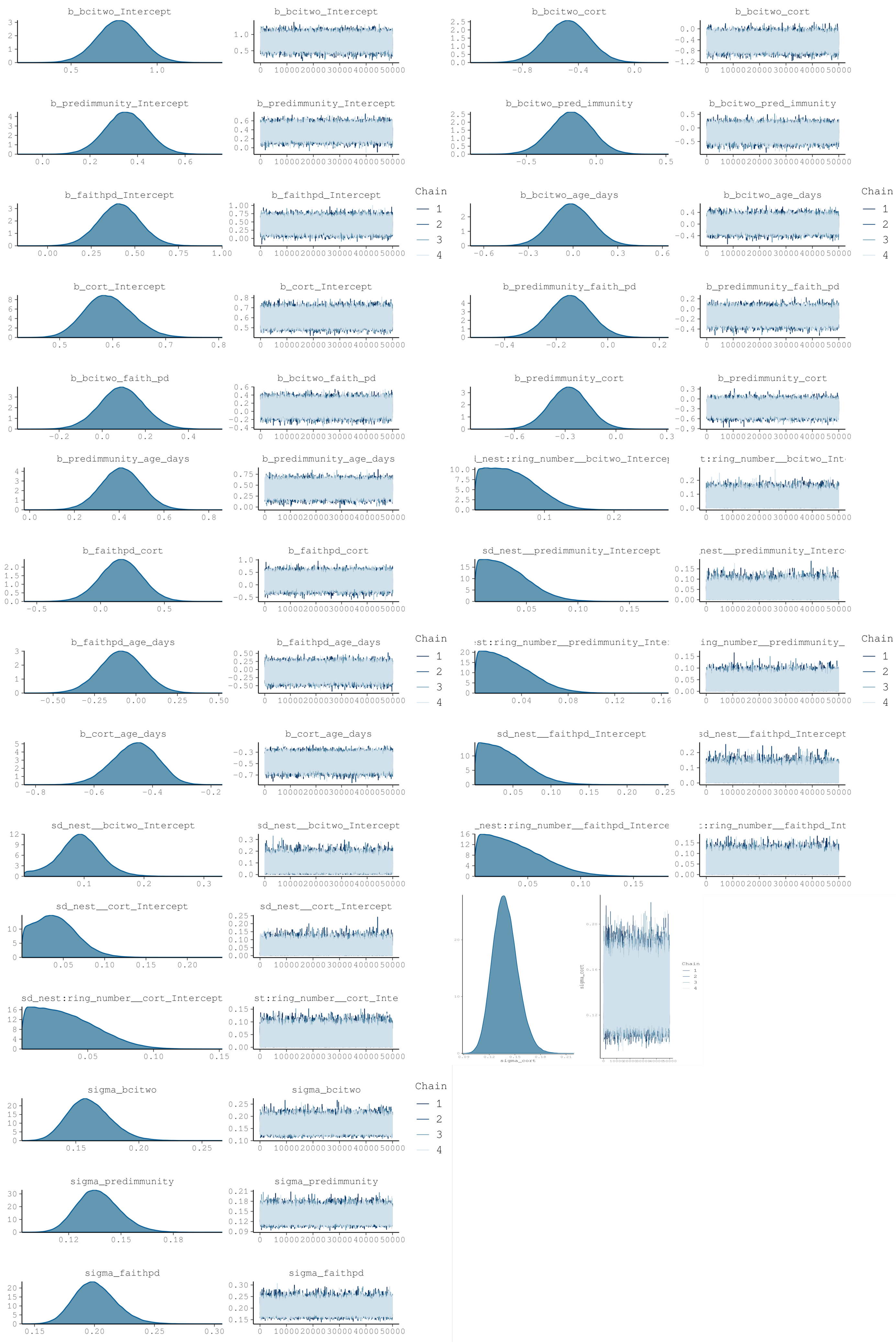
b_bcitwo_Intercept
b_bcitwo_cort
b_predimmunity_Intercept
b_bcitwo_pred_immunity
b_faithpd_Intercept
b_bcitwo_age_days
Chain
1
2
3
4
b_cort_Intercept
b_predimmunity_faith_pd
b_bcitwo_faith_pd
b_predimmunity_cort
b_predimmunity_age_days
b_faithpd_cort
sd_nest__predimmunity_Intercept
b_faithpd_age_days
b_cort_age_days
sd_nest__faithpd_Intercept
sd_nest__bcitwo_Intercept
sd_nest__cort_Intercept
sd_nest:ring_number__cort_Intercept
sigma_cort
sigma_bcitwo
sigma_predimmunity
sigma_faithpd

#### 5.3 Compare distribution of response variable to distributions of predicted response variable

```
#Posterior predictive checks
#Loop to save all distributions plot
responses <- c("bcitwo", "predimmunity", "faithpd", "cort")
response_names <- c("bci", "immune", "faith", "cort")
for (i in seq_along(responses)) {
  pp_check_plot <- pp_check(model_immunity_shannon, resp = responses[i], ndraws = 200)
  filename <- paste0("distribution_faith_", response_names[i])
  ggsave(filename = paste0(filename, ".svg"), plot = pp_check_plot, device = "svg", width = 8,
height = 10)
}
```

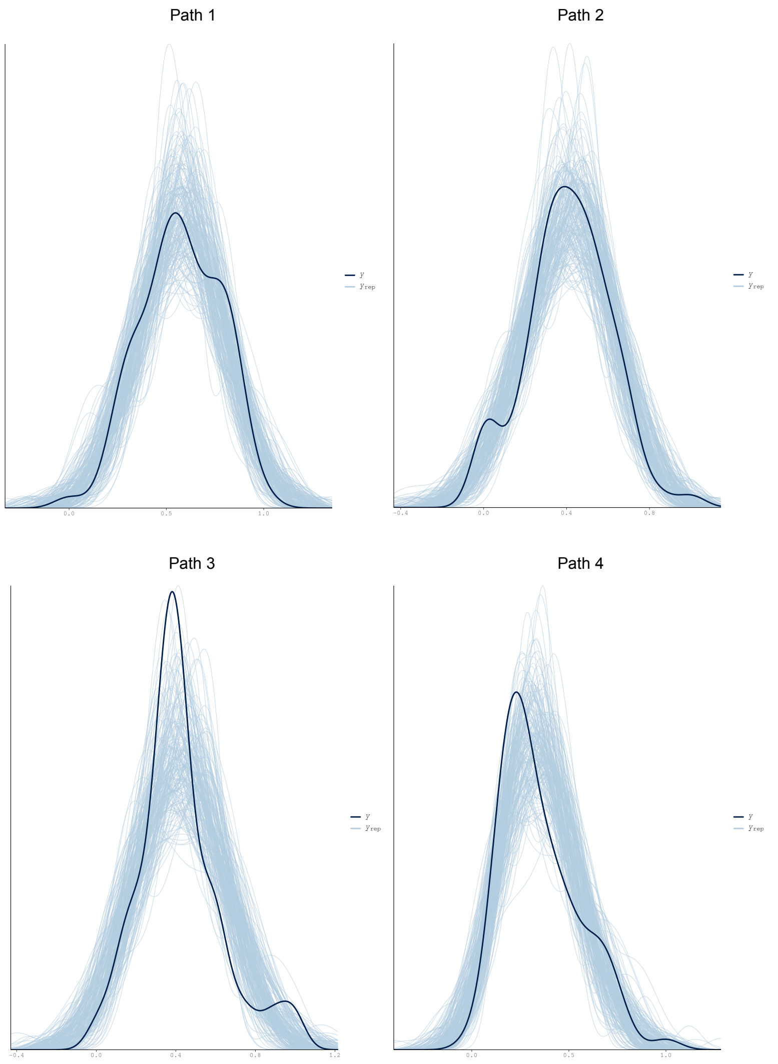


#### 5.4 Plot model posterior and credible intervals

```
#Plot Model effects
plot2 <-mcmc_plot(model_immunity_shannon, type = "intervals",prob_outer=0.95, prob=0.95,
                  variable =c("b_bcitwo_faith_pd", "b_bcitwo_cort", "b_bcitwo_pred_immunity",
"b_bcitwo_age_days",
                               "b_predimmunity_faith_pd", "b_predimmunity_cort",
"b_predimmunity_age_days",
                               "b_faithpd_cort", "b_faithpd_age_days",
                               "b_cort_age_days"))

plot2 <- plot2 + theme_classic() + geom_vline(xintercept = 0, linetype="dotted", color="blue")+
theme(axis.text.x = element_text(size = 16),  # Adjust the size as needed
axis.text.y = element_text(size = 16))+
theme(text = element_text(family = "Arial"))

ggsave(filename="16s_effect_sizes_faith.svg", plot=plot1, device = "svg", width = 8, height =
10)
```

### 6. Model diagnostics - N° of observed ASV's

#### 6.1 Model summary

```
#Model summary
summary_asv<- summary(model_immunity_asv)

#Bayes R2
R2m_asv <- bayes_R2(model_immunity_asv,re_formula=NA)
R2c_asv <- bayes_R2(model_immunity_asv)
```

#### 6.2 Model diagnostics

```
# Model diagnostics
diagnostic_asv <- plot(model_immunity_asv)

#Loop to save all diagnostic plots
diagnostic_plots <- list()
for (i in 1:length(diagnostic_asv)) {
  diagnostic_plots[[i]] <- diagnostic_asv[[i]]
  filename <- paste0("diagnostic", i, "_asv")
  ggsave(filename = paste0(filename, ".png"), plot = diagnostic_plots[[i]], device = "png",
dpi= 300)
}
```

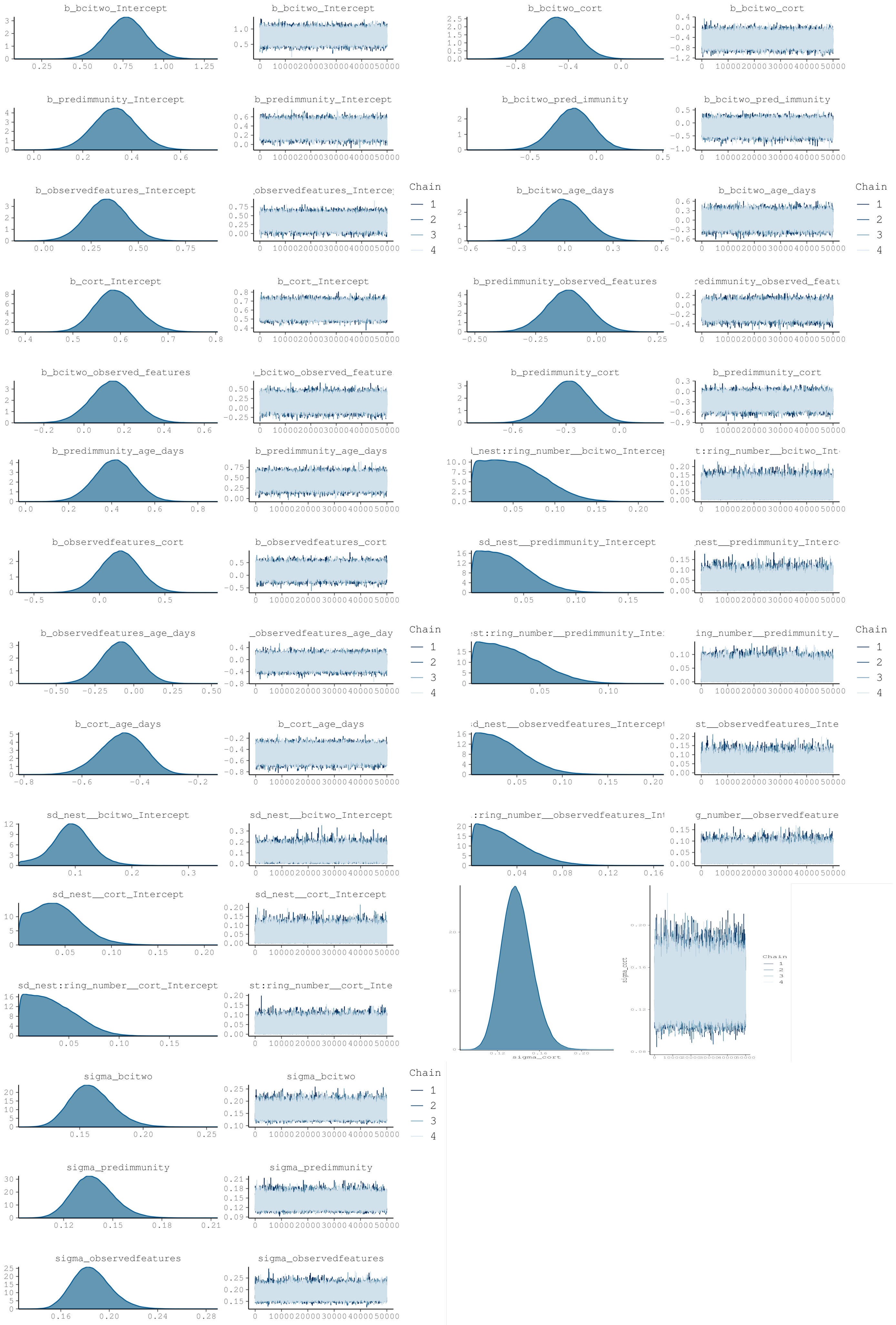

b_bcitwo_Intercept
b_bcitwo_cort
b_predimmunity_Intercept
b_bcitwo_pred_immunity
b_observedfeatures_Intercept
b_bcitwo_age_days
b_cort_Intercept
b_predimmunity_observed_features
b_bcitwo_observed_features
b_predimmunity_cort
b_predimmunity_age_days
b_observedfeatures_cort
sd_nest__predimmunity_Intercept
b_observedfeatures_age_days
b_cort_age_days
sd_nest__bcitwo_Intercept
sd_nest__cort_Intercept
sd_nest:ring_number__cort_Intercept
sigma_cort
sigma_bcitwo
sigma_predimmunity
sigma_observedfeatures
Chain
1
2
3
4

##### 6.3 Compare distribution of response variable to distributions of predicted response variable

```r
#Posterior predictive checks
#Loop to save all distributions plot
responses <- c("bcitwo", "predimmunity", "observedfeatures", "cort")
response_names <- c("bci", "immune", "asv", "cort")
for (i in seq_along(responses)) {
  pp_check_plot <- pp_check(model_immunity_asv, resp = responses[i], ndraws = 200)
  filename <- paste0("distribution_asv_", response_names[i])
  ggsave(filename = paste0(filename, ".svg"), plot = pp_check_plot, device = "svg", width = 8,
height = 10)
}
```

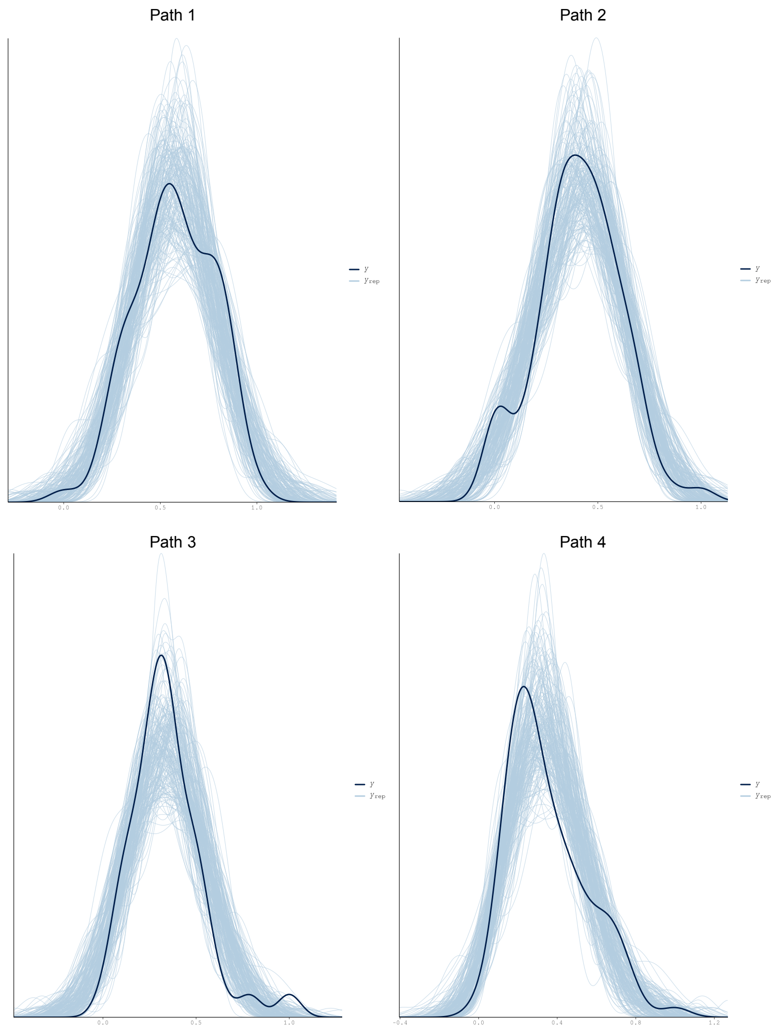


##### 6.4 Plot model posterior and credible intervals

```
#Plot Model effects
plot3 <-mcmc_plot(model_immunity_shannon, type = "intervals",prob_outer=0.95, prob=0.95,
                  variable =c("b_bcitwo_observed_features", "b_bcitwo_cort",
"b_bcitwo_pred_immunity", "b_bcitwo_age_days",
                               "b_predimmunity_observed_features", "b_predimmunity_cort",
"b_predimmunity_age_days",
                               "b_observedfeatures_cort", "b_observedfeatures_age_days",
                               "b_cort_age_days"))

plot3 <- plot3 + theme_classic() + geom_vline(xintercept = 0, linetype="dotted", color="blue")+
theme(axis.text.x = element_text(size = 16),  # Adjust the size as needed
axis.text.y = element_text(size = 16))+
theme(text = element_text(family = "Arial"))

ggsave(filename="16s_effect_sizes_asv.svg", plot=plot1, device = "svg", width = 8, height = 10)
```

#### B) 28S rRNA (eukaryotic microbiota) SEM analysis

##### 1. Build the latent variable

```
#Load the data
metadata <- readRDS("28s_metadata_immune.rds")

#Scale immune assay scores
metadata$std_ha <- as.numeric (scale(metadata$ha))
metadata$std_hl <- as.numeric(scale(metadata$hl))
metadata$std_bka <- as.numeric(scale(metadata$bka))
metadata$std_lyso <-as.numeric(scale(metadata$lyso))
metadata$std_igy <- as.numeric(scale(metadata$igy))
metadata$std_hapto <- as.numeric(scale(metadata$hapto))

# Exploratory factor analysis - all imune assays included.

model_factor <- 'immunity =~  std_ha + std_hl + std_bka + std_lyso + std_igy + std_hapto'
fit_factor <-  efa(model_factor, data = metadata, cluster = c("ring_number"), missing = "fiml",
estimator = "MLR", std.lv=T)

#Model summary
summary(fit_factor, fit.measures = TRUE, standardized = TRUE, rsquare = TRUE

#Extract main model fit measures
fitMeasures(fit_factor, c("pvalue.scaled","cfi.robust","rmsea.robust","srmr"))

# Exploratory factor analysis excluding haptoglobin.

model_factor1 <- 'immunity =~  std_ha + std_hl + std_bka + std_lyso + std_igy'

fit_factor1 <-  efa(model_factor1, data = metadata, cluster = c("ring_number"), missing =
"fiml", estimator = "MLR", std.lv=T)

#Model summary
summary(fit_factor1, fit.measures = TRUE, standardized = TRUE, rsquare = TRUE)
```

```
#Extract main model fit measures
fitMeasures(fit_factor1, c("pvalue.scaled","cfi.robust","rmsea.robust","srmr"))
```

#### 2. Define SEM for each diversity measurement

```
#scale all predictors to range between 0-1 if they are not already naturally on that scale

#define scaling function:
range.use <- function(x,min.use,max.use){ (x - min(x,na.rm=T))/(max(x,na.rm=T)-min(x,na.rm=T))
* (max.use - min.use) + min.use }
scalecols<-c("bci_two", "shannon_entropy", "faith_pd", "observed_features", "cort",
"pred_immunity", "age_days")


for(i in 1:ncol(metadata[,which(colnames(metadata)%in%scalecols)])){
  metadata[,which(colnames(metadata)%in%scalecols)][,i]<-
range.use(metadata[,which(colnames(metadata)%in%scalecols)][,i],0,1)
}

# Define structural equation model paths

#Shannon
path1 <- bf(bci_two ~ shannon_entropy + cort + pred_immunity + age_days + (1|nest/ring_number))
path2 <- bf(pred_immunity ~  shannon_entropy + cort + age_days + (1|nest/ring_number))
path3 <- bf(shannon_entropy ~ cort + age_days + (1|nest/ring_number))
path4 <- bf(cort ~ age_days + (1|nest/ring_number)) + skew_normal()

sem_immunity_shannon <- path1 + path2 + path3 + path4

#Faith PD
path1 <- bf(bci_two ~ faith_pd + cort + pred_immunity + age_days + (1|nest/ring_number))
path2 <- bf(pred_immunity ~  faith_pd + cort + age_days + (1|nest/ring_number))
path3 <- bf(faith_pd ~ cort + age_days + (1|nest/ring_number))
path4 <- bf(cort ~ age_days + (1|nest/ring_number)) + skew_normal()

sem_immunity_faith <- path1 + path2 + path3 + path4

#N° of observed ASV's
path1 <- bf(bci_two ~ observed_features + cort + pred_immunity + age_days +
(1|nest/ring_number))
path2 <- bf(pred_immunity ~  observed_features + cort + age_days + (1|nest/ring_number))
path3 <- bf(observed_features ~ cort + age_days + (1|nest/ring_number))
path4 <- bf(cort ~ age_days + (1|nest/ring_number)) + skew_normal()

sem_immunity_asv <- path1 + path2 + path3 + path4
```

#### 3. Run brms

```
ncores = detectCores()
options(mc.cores = parallel::detectCores())

#Shannon
model_immunity_shannon <-brm(sem_immunity_shannon + set_rescor(FALSE),
                         data = metadata,
```

```
                         warmup = 50000, iter = 100000,
                         control = list(adapt_delta = 0.99, max_treedepth = 15),
                         cores=ncores, chains=4, init=1000)

#Faith PD
model_immunity_faith <-brm(sem_immunity_faith + set_rescor(FALSE),
                         data = metadata,
                         warmup = 50000, iter = 100000,
                         control = list(adapt_delta = 0.99, max_treedepth = 15),
                         cores=ncores, chains=4, init=1000)

#N° of observed ASV's
model_immunity_asv <-brm(sem_immunity_asv + set_rescor(FALSE),
                         data = metadata,
                         warmup = 50000, iter = 100000,
                         control = list(adapt_delta = 0.99, max_treedepth = 15),
                         cores=ncores, chains=4, init=1000)
```

### 4. Model Diagnostics - Shannon

#### 4.1 Model Summary

```
#Model summary
summary_shannon<- summary(model_immunity_shannon)

#Bayes R2
R2m_shannon <- bayes_R2(model_immunity_shannon,re_formula=NA)
R2c_shannon <- bayes_R2(model_immunity_shannon)
```

#### 4.2 Model diagnostics

```
# Model diagnostics
diagnostic_shannon <- plot(model_immunity_shannon)

#Loop to save all diagnostic plots
diagnostic_plots <- list()
for (i in 1:length(diagnostic_shannon)) {
  diagnostic_plots[[i]] <- diagnostic_shannon[[i]]
  filename <- paste0("diagnostic", i, "_shannon")
  ggsave(filename = paste0(filename, ".png"), plot = diagnostic_plots[[i]], device = "png",
dpi=300)
}
```

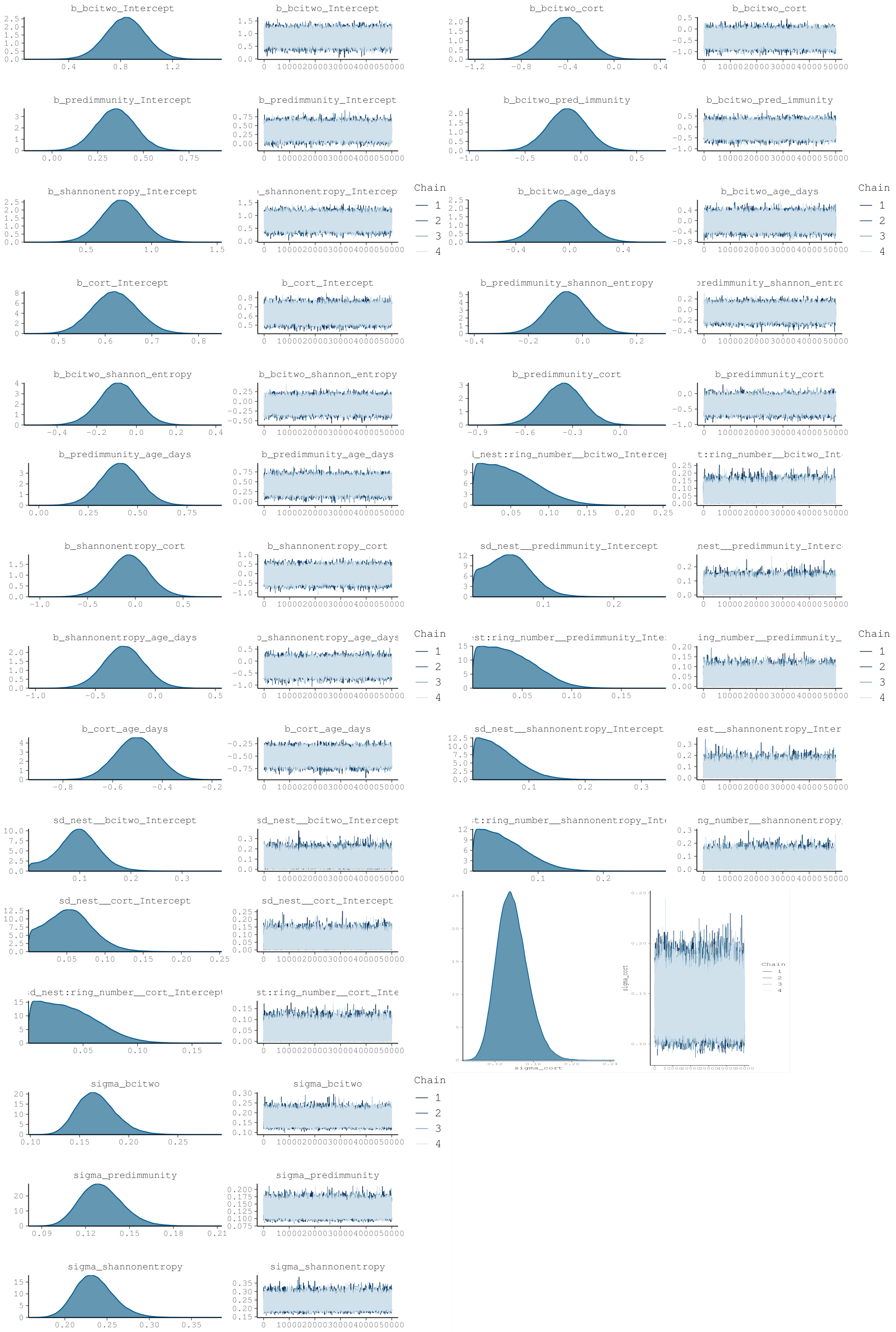
b_bcitwo_Intercept
b_bcitwo_cort
b_predimmunity_Intercept
b_bcitwo_pred_immunity
b_shannonentropy_Intercept
b_bcitwo_age_days
b_cort_Intercept
b_predimmunity_shannon_entropy
b_bcitwo_shannon_entropy
b_predimmunity_cort
b_predimmunity_age_days
b_shannonentropy_cort
sd_nest__predimmunity_Intercept
b_shannonentropy_age_days
b_cort_age_days
sd_nest__shannonentropy_Intercept
sd_nest__bcitwo_Intercept
sd_nest__cort_Intercept
sigma_cort
sigma_bcitwo
sigma_predimmunity
sigma_shannonentropy
Chain
1
2
3
4

#### 4.3 Compare distribution of response variable to distributions of predicted response variable

```
#Posterior predictive checks (one by one)
distribution_shannon_path1 <- pp_check(model_immunity_shannon, resp="bcitwo", ndraws=200)
distribution_shannon_path2 <- pp_check(model_immunity_shannon, resp="predimmunity", ndraws=200)
distribution_shannon_path3 <- pp_check(model_immunity_shannon, resp="shannonentropy",
ndraws=200)
distribution_shannon_path4 <- pp_check(model_immunity_shannon, resp="cort", ndraws=200)

#Loop to save all distributions plot
responses <- c("bcitwo", "predimmunity", "shannonentropy", "cort")
response_names <- c("bci", "immune", "shannon", "cort")
for (i in seq_along(responses)) {
  pp_check_plot <- pp_check(model_immunity_shannon, resp = responses[i], ndraws = 200)
  filename <- paste0("distribution_shannon_", response_names[i])
  ggsave(filename = paste0(filename, ".svg"), plot = pp_check_plot, device = "svg", width = 8,
height = 10)
}
```

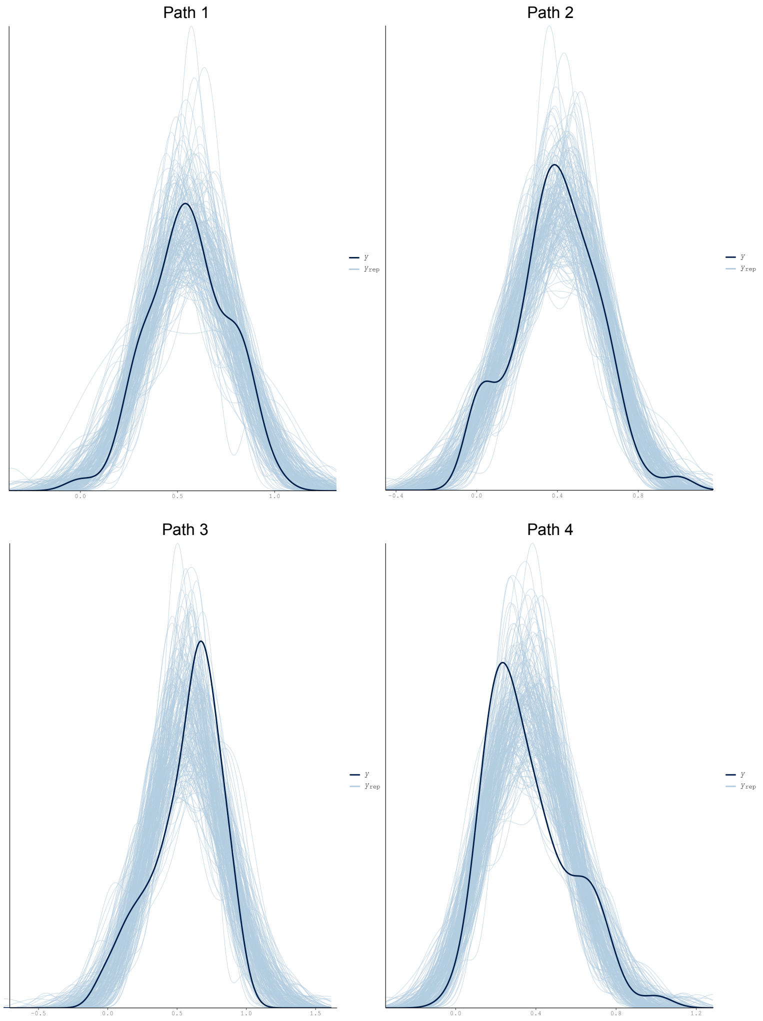

##### 4.4 Plot model posterior and credible intervals

```
#Plot Model effects
plot4 <-mcmc_plot(model_immunity_shannon, type = "intervals",prob_outer=0.95, prob=0.95,
                  variable =c("b_bcitwo_shannon_entropy", "b_bcitwo_cort",
"b_bcitwo_pred_immunity", "b_bcitwo_age_days",
                               "b_predimmunity_shannon_entropy", "b_predimmunity_cort",
"b_predimmunity_age_days",
                               "b_shannonentropy_cort", "b_shannonentropy_age_days",
                               "b_cort_age_days"))

plot4 <- plot4 + theme_classic() + geom_vline(xintercept = 0, linetype="dotted", color="blue")+
theme(axis.text.x = element_text(size = 16),  # Adjust the size as needed
axis.text.y = element_text(size = 16))+
theme(text = element_text(family = "Arial"))

ggsave(filename="16s_effect_sizes_shannon.svg", plot=plot4, device = "svg", width = 8, height =
10)
```

#### 5. Model diagnostics - Faith PD

##### 5.1 Model summary

```
#Model summary
summary_faith<- summary(model_immunity_faith)

#Bayes R2
R2m_faith <- bayes_R2(model_immunity_faith,re_formula=NA)
R2c_faith <- bayes_R2(model_immunity_faith)
```

##### 5.2 Model diagnostics

```
# Model diagnostics
diagnostic_faith <- plot(model_immunity_faith)

#Loop to save all diagnostic plots
diagnostic_plots <- list()
for (i in 1:length(diagnostic_faith)) {
  diagnostic_plots[[i]] <- diagnostic_faith[[i]]
  filename <- paste0("diagnostic", i, "_faith")
  ggsave(filename = paste0(filename, ".png"), plot = diagnostic_plots[[i]], device = "png",
dpi=200)
}
```

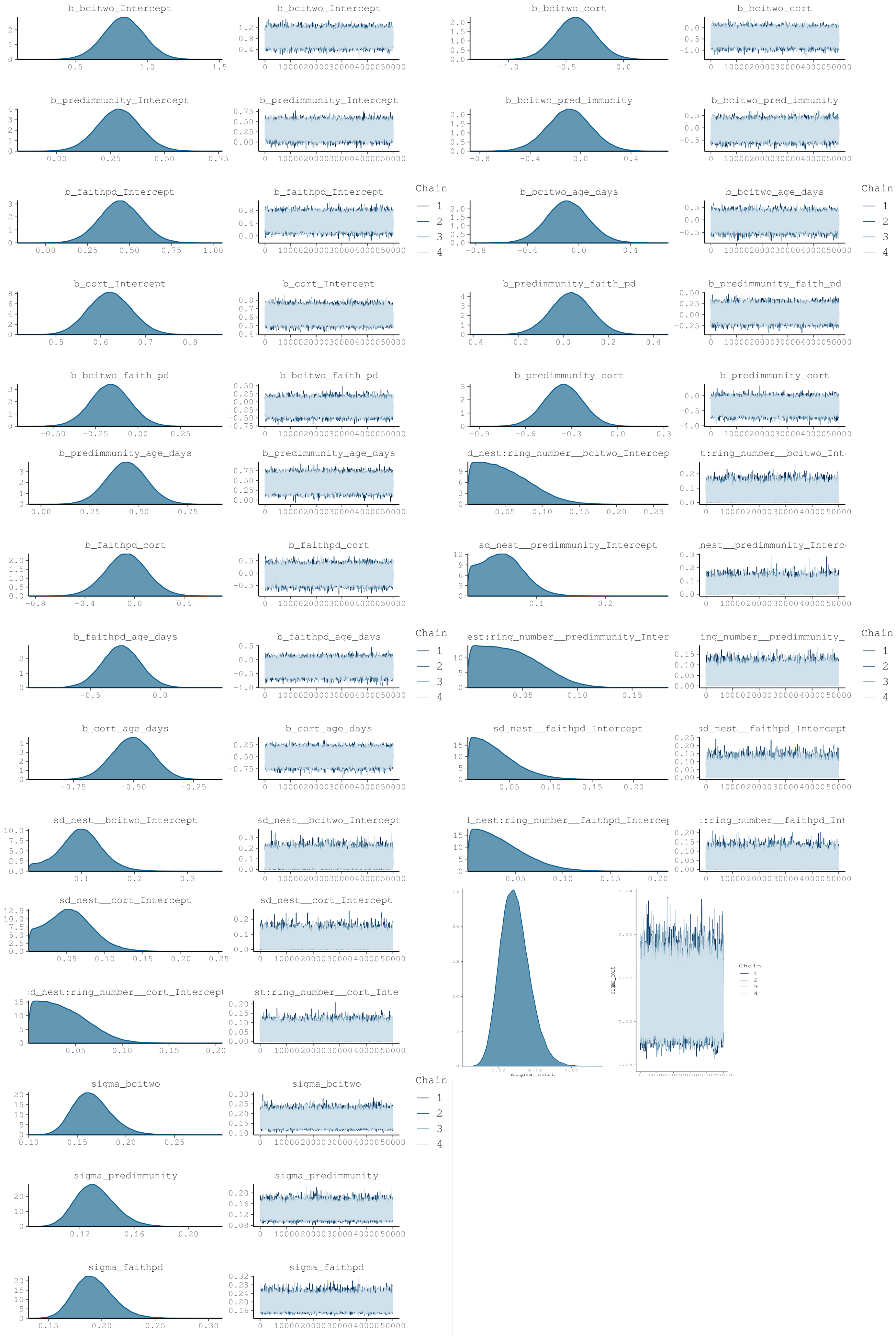
b_bciwo_Intercept
b_bcitwo_cort
b_predimmunity_Intercept
b_bcitwo_pred_immunity
b_faithpd_Intercept
b_bcitwo_age_days
Chain
1
2
3
4
b_cort_Intercept
b_predimmunity_faith_pd
b_bcitwo_faith_pd
b_predimmunity_cort
b_predimmunity_age_days
d_nest:ring_number__bcitwo_Intercep
t:ring_number__bcitwo_Int
b_faithpd_cort
sd_nest__predimmunity_Intercept
nest__predimmunity_Interc
b_faithpd_age_days
est:ring_number__predimmunity_Inter
ing_number__predimmunity_
b_cort_age_days
sd_nest__faithpd_Intercept
sd_nest__bcitwo_Intercept
l_nest:ring_number__faithpd_Intercep
:ring_number__faithpd_Int
sd_nest__cort_Intercept
sigma_cort
d_nest:ring_number__cort_Intercept
st:ring_number__cort_Inte
sigma_bcitwo
sigma_predimmunity
sigma_faithpd

##### 5.3 Compare distribution of response variable to distributions of predicted response variable

```r
#Posterior predictive checks
#Loop to save all distributions plot
responses <- c("bcitwo", "predimmunity", "faithpd", "cort")
response_names <- c("bci", "immune", "faith", "cort")
for (i in seq_along(responses)) {
  pp_check_plot <- pp_check(model_immunity_faith, resp = responses[i], ndraws = 200)
  filename <- paste0("distribution_faith_", response_names[i])
  ggsave(filename = paste0(filename, ".svg"), plot = pp_check_plot, device = "svg", width = 8,
height = 10)
}
```

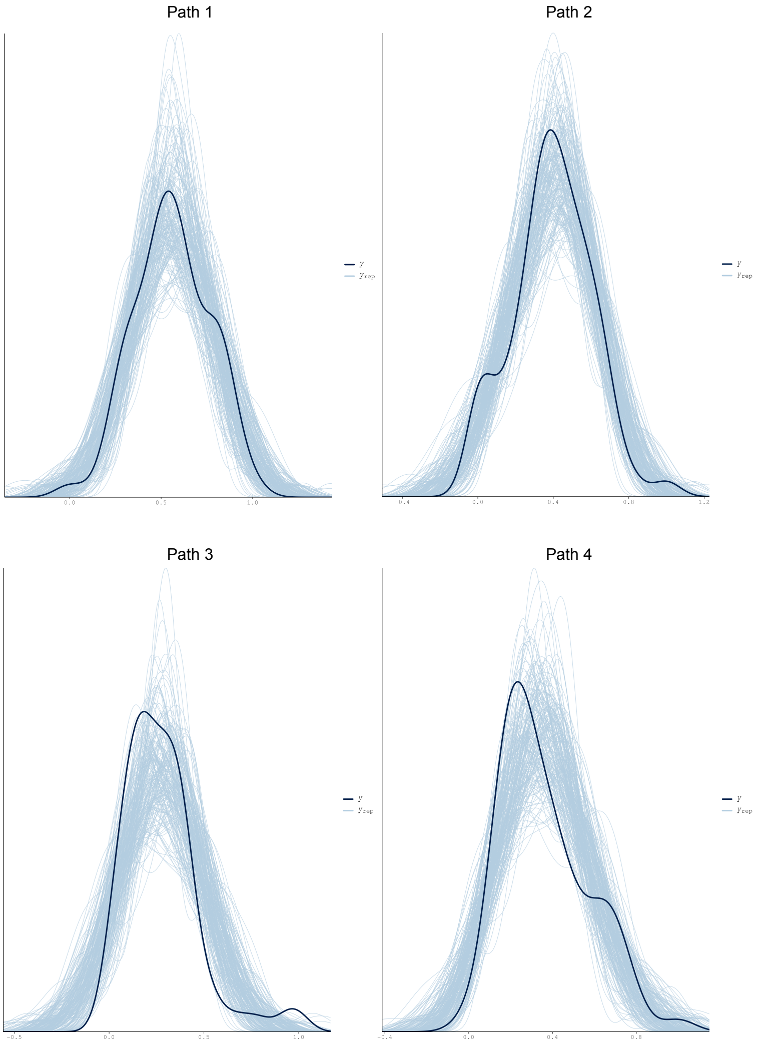


##### 5.4 Plot model posterior and credible intervals

```
#Plot Model effects
plot5 <-mcmc_plot(model_immunity_shannon, type = "intervals",prob_outer=0.95, prob=0.95,
                  variable =c("b_bcitwo_faith_pd", "b_bcitwo_cort", "b_bcitwo_pred_immunity",
"b_bcitwo_age_days",
                               "b_predimmunity_faith_pd", "b_predimmunity_cort",
"b_predimmunity_age_days",
                               "b_faithpd_cort", "b_faithpd_age_days",
                               "b_cort_age_days"))

plot5 <- plot5 + theme_classic() + geom_vline(xintercept = 0, linetype="dotted", color="blue")+
theme(axis.text.x = element_text(size = 16),  # Adjust the size as needed
axis.text.y = element_text(size = 16))+
theme(text = element_text(family = "Arial"))

ggsave(filename="16s_effect_sizes_faith.svg", plot=plot5, device = "svg", width = 8, height =
10)
```

### 6. Model diagnostics - N° of observed ASV's

#### 6.1 Model summary

```
#Model summary
summary_asv<- summary(model_immunity_asv)

#Bayes R2
R2m_asv <- bayes_R2(model_immunity_asv,re_formula=NA)
R2c_asv <- bayes_R2(model_immunity_asv)
```

#### 6.2 Model diagnostics

```
# Model diagnostics
diagnostic_asv <- plot(model_immunity_asv)

#Loop to save all diagnostic plots
diagnostic_plots <- list()
for (i in 1:length(diagnostic_asv)) {
  diagnostic_plots[[i]] <- diagnostic_asv[[i]]
  filename <- paste0("diagnostic", i, "_asv")
  ggsave(filename = paste0(filename, ".png"), plot = diagnostic_plots[[i]], device = "png",
dpi= 300)
}
```

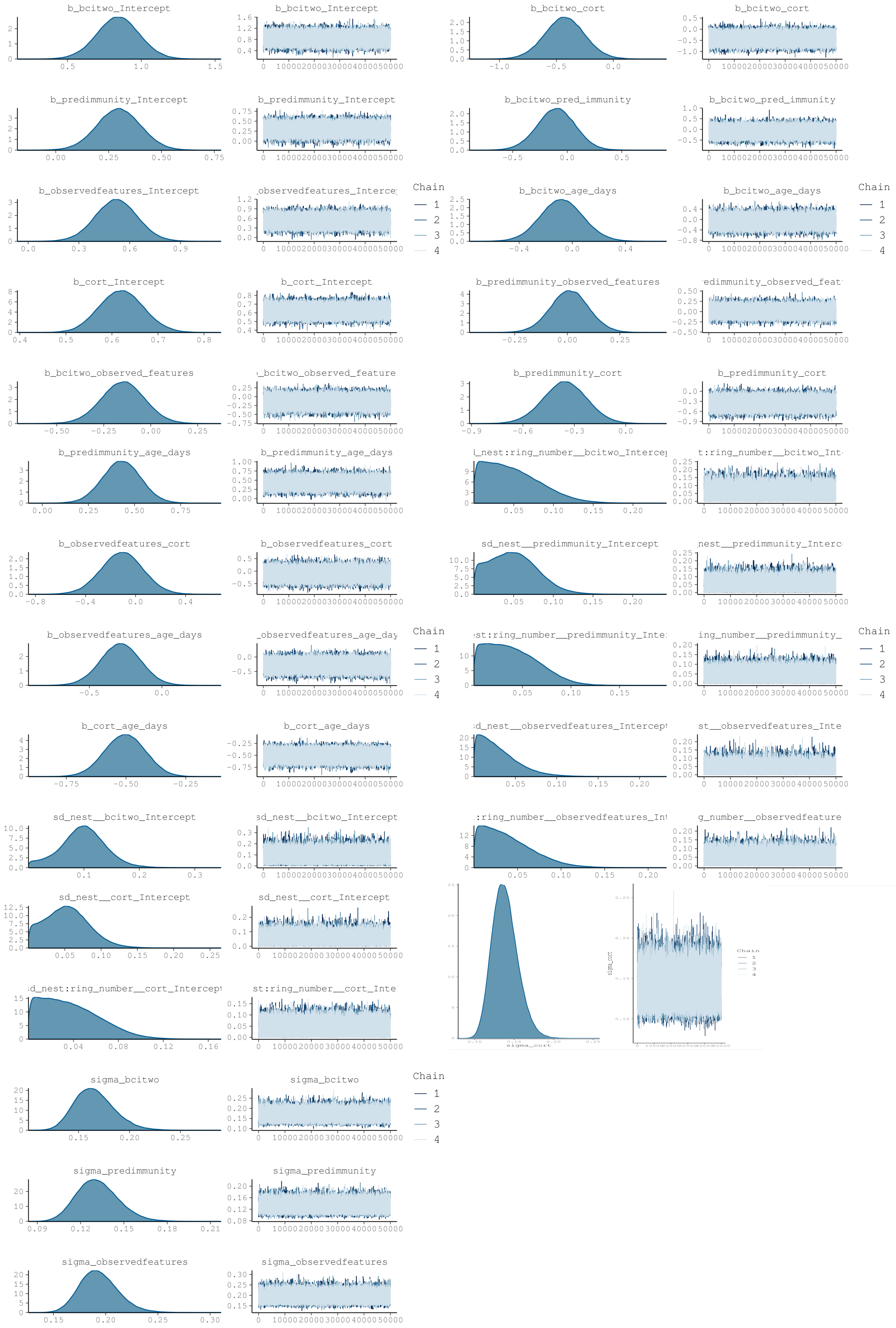
b_bcitwo_Intercept
b_bcitwo_cort
b_predimmunity_Intercept
b_bcitwo_pred_immunity
b_observedfeatures_Intercept
observedfeatures_Interce
Chain
1
2
3
4
b_bcitwo_age_days
b_cort_Intercept
b_predimmunity_observed_features
edimmunity_observed_feat
b_bcitwo_observed_features
bcitwo_observed_feature
b_predimmunity_cort
b_predimmunity_age_days
_nest:ring_number__bcitwo_Interce
t:ring_number__bcitwo_Int
b_observedfeatures_cort
sd_nest__predimmunity_Intercept
nest__predimmunity_Interc
b_observedfeatures_age_days
observedfeatures_age_day
st:ring_number__predimmunity_Inte
ing_number__predimmunity_
b_cort_age_days
d_nest__observedfeatures_Intercept
st__observedfeatures_Inte
sd_nest__bcitwo_Intercept
:ring_number__observedfeatures_Int
g_number__observedfeature
sd_nest__cort_Intercept
sigma_cort
d_nest:ring_number__cort_Intercept
st:ring_number__cort_Inte
sigma_bcitwo
sigma_predimmunity
sigma_observedfeatures

#### 6.3 Compare distribution of response variable to distributions of predicted response variable

```
#Posterior predictive checks
#Loop to save all distributions plot
responses <- c("bcitwo", "predimmunity", "observedfeatures", "cort")
response_names <- c("bci", "immune", "asv", "cort")
for (i in seq_along(responses)) {
  pp_check_plot <- pp_check(model_immunity_asv, resp = responses[i], ndraws = 200)
  filename <- paste0("distribution_asv_", response_names[i])
  ggsave(filename = paste0(filename, ".svg"), plot = pp_check_plot, device = "svg", width = 8,
height = 10)
}
```

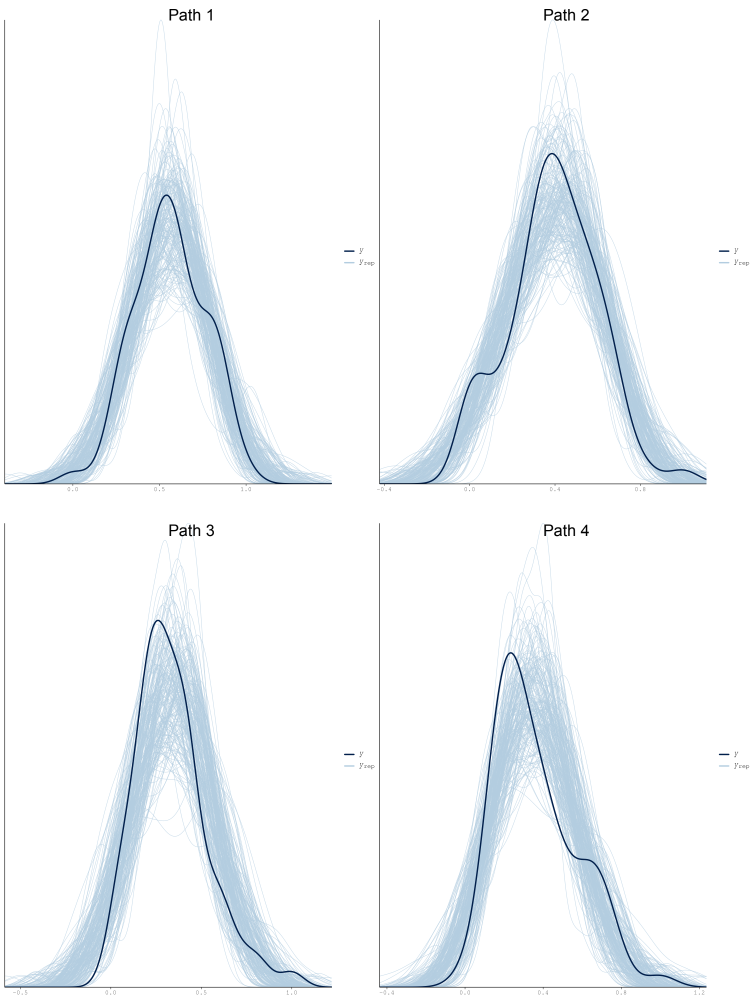


#### 5.4 Plot model posterior and credible intervals

```
#Plot Model effects
plot2 <-mcmc_plot(model_immunity_shannon, type = "intervals",prob_outer=0.95, prob=0.95,
                  variable =c("b_bcitwo_faith_pd", "b_bcitwo_cort", "b_bcitwo_pred_immunity",
"b_bcitwo_age_days",
                               "b_predimmunity_faith_pd", "b_predimmunity_cort",
"b_predimmunity_age_days",
                               "b_faithpd_cort", "b_faithpd_age_days",
                               "b_cort_age_days"))

plot2 <- plot2 + theme_classic() + geom_vline(xintercept = 0, linetype="dotted", color="blue")+
theme(axis.text.x = element_text(size = 16),  # Adjust the size as needed
axis.text.y = element_text(size = 16))+
theme(text = element_text(family = "Arial"))

ggsave(filename="16s_effect_sizes_faith.svg", plot=plot1, device = "svg", width = 8, height =
10)
```
