## Supplementary material for "The gut microbiota-immune-brain axis in a wild vertebrate: dynamic interactions and health impacts": Bayesian-SEM-Haptoglobin

### Bayesian Structural Equation Modelling - modelling Haptoglobin immune assay

#### Table of contents

**Bayesian Structural Equation Modelling - modelling Haptoglobin immune assay**
- Table of contents
- A) 16S rRNA (bacterial microbiota) SEM analysis
  - 1. Define SEM for each diversity measurement
  - 2. Run brms
  - 3. Model Diagnostics Shannon
    - 3.1 Model Summary
    - 3.2 Model diagnostics
    - 3.3 Compare distribution of response variable to distributions of predicted response variable
    - 3.4 Plot model posterior and credible intervals
  - 4. Model diagnostics - Faith PD
    - 4.1 Model summary
    - 4.2 Model diagnostics
    - 4.3 Compare distribution of response variable to distributions of predicted response variable
    - 4.4 Plot model posterior and credible intervals
  - 5. Model diagnostics - N° of observed ASV's
    - 5.1 Model summary
    - 5.2 Model diagnostics
    - 5.3 Compare distribution of response variable to distributions of predicted response variable
    - 5.4 Plot model posterior and credible intervals
- B) 28S rRNA (eukaryotic microbiota) SEM analysis
  - 1. Define SEM for each diversity measurement
  - 2. Run brms
  - 3. Model Diagnostics - Shannon
    - 3.1 Model Summary
    - 3.2 Model diagnostics
    - 3.3 Compare distribution of response variable to distributions of predicted response variable
    - 3.4 Plot model posterior and credible intervals
  - 4. Model diagnostics - Faith PD
    - 4.1 Model summary
    - 4.2 Model diagnostics
    - 4.3 Compare distribution of response variable to distributions of predicted response variable
    - 4.4 Plot model posterior and credible intervals
  - 5. Model diagnostics - N° of observed ASV's
    - 5.1 Model summary
    - 5.2 Model diagnostics
    - 5.3 Compare distribution of response variable to distributions of predicted response variable
    - 5.4 Plot model posterior and credible intervals

#### A) 16S rRNA (bacterial microbiota) SEM analysis

### 1. Define SEM for each diversity measurement

```
#Load Packages
library(brms)
library(rstan)
library(bayesplot)
library(bayestestR)
library(parallel)
library(svglite)
library(ggplot2)

#Load the data
metadata <- readRDS("16s_metadata_immune.rds")

#scale all predictors to range between 0-1 if they are not already naturally on that scale

#define scaling function:
range.use <- function(x,min.use,max.use){ (x - min(x,na.rm=T))/(max(x,na.rm=T)-min(x,na.rm=T))
* (max.use - min.use) + min.use }
scalecols<-c("bci_two", "shannon_entropy", "faith_pd", "observed_features", "cort", "hapto",
"age_days")


for(i in 1:ncol(metadata[,which(colnames(metadata)%in%scalecols)])){
  metadata[,which(colnames(metadata)%in%scalecols)][,i]<-
range.use(metadata[,which(colnames(metadata)%in%scalecols)][,i],0,1)
}

# Define structural equation model paths

#Shannon
path1 <- bf(bci_two ~ shannon_entropy + cort + hapto+ age_days + (1|nest/ring_number))
path2 <- bf(hapto~  shannon_entropy + cort + age_days + (1|nest/ring_number))
path3 <- bf(shannon_entropy ~ cort + age_days + (1|nest/ring_number))
path4 <- bf(cort ~ age_days + (1|nest/ring_number)) + skew_normal()

sem_hapto_shannon <- path1 + path2 + path3 + path4

#Faith PD
path1 <- bf(bci_two ~ faith_pd + cort + hapto+ age_days + (1|nest/ring_number))
path2 <- bf(hapto~  faith_pd + cort + age_days + (1|nest/ring_number))
path3 <- bf(faith_pd ~ cort + age_days + (1|nest/ring_number))
path4 <- bf(cort ~ age_days + (1|nest/ring_number)) + skew_normal()

sem_hapto_faith <- path1 + path2 + path3 + path4

#N° of observed ASV's
path1 <- bf(bci_two ~ observed_features + cort + hapto+ age_days + (1|nest/ring_number))
path2 <- bf(hapto~  observed_features + cort + age_days + (1|nest/ring_number))
path3 <- bf(observed_features ~ cort + age_days + (1|nest/ring_number))
path4 <- bf(cort ~ age_days + (1|nest/ring_number)) + skew_normal()

sem_hapto_asv <- path1 + path2 + path3 + path4
```

#### 2. Run brms

```
ncores = detectCores()
options(mc.cores = parallel::detectCores())

#Shannon
model_hapto_shannon <-brm(sem_hapto_shannon + set_rescor(FALSE),
                          data = metadata,
                          warmup = 50000, iter = 100000,
                          control = list(adapt_delta = 0.99, max_treedepth = 15),
                          cores=ncores, chains=4, init=1000)

#Faith PD
model_hapto_faith <-brm(sem_hapto_faith + set_rescor(FALSE),
                          data = metadata,
                          warmup = 50000, iter = 100000,
                          control = list(adapt_delta = 0.99, max_treedepth = 15),
                          cores=ncores, chains=4, init=1000)

#N° of observed ASV's
model_hapto_asv <-brm(sem_hapto_asv + set_rescor(FALSE),
                          data = metadata,
                          warmup = 50000, iter = 100000,
                          control = list(adapt_delta = 0.99, max_treedepth = 15),
                          cores=ncores, chains=4, init=1000)
```

#### 3. Model Diagnostics Shannon

##### 3.1 Model Summary

```
#Model summary
summary_shannon<- summary(model_hapto_shannon)

#Bayes R2
R2m_shannon <- bayes_R2(model_hapto_shannon,re_formula=NA)
R2c_shannon <- bayes_R2(model_hapto_shannon)
```

##### 3.2 Model diagnostics

```
# Model diagnostics
diagnostic_shannon <- plot(model_hapto_shannon)

#Loop to save all diagnostic plots
diagnostic_plots <- list()
for (i in 1:length(diagnostic_shannon)) {
  diagnostic_plots[[i]] <- diagnostic_shannon[[i]]
  filename <- paste0("diagnostic", i, "_shannon")
  ggsave(filename = paste0(filename, ".png"), plot = diagnostic_plots[[i]], device = "png",
dpi=300)
}
```

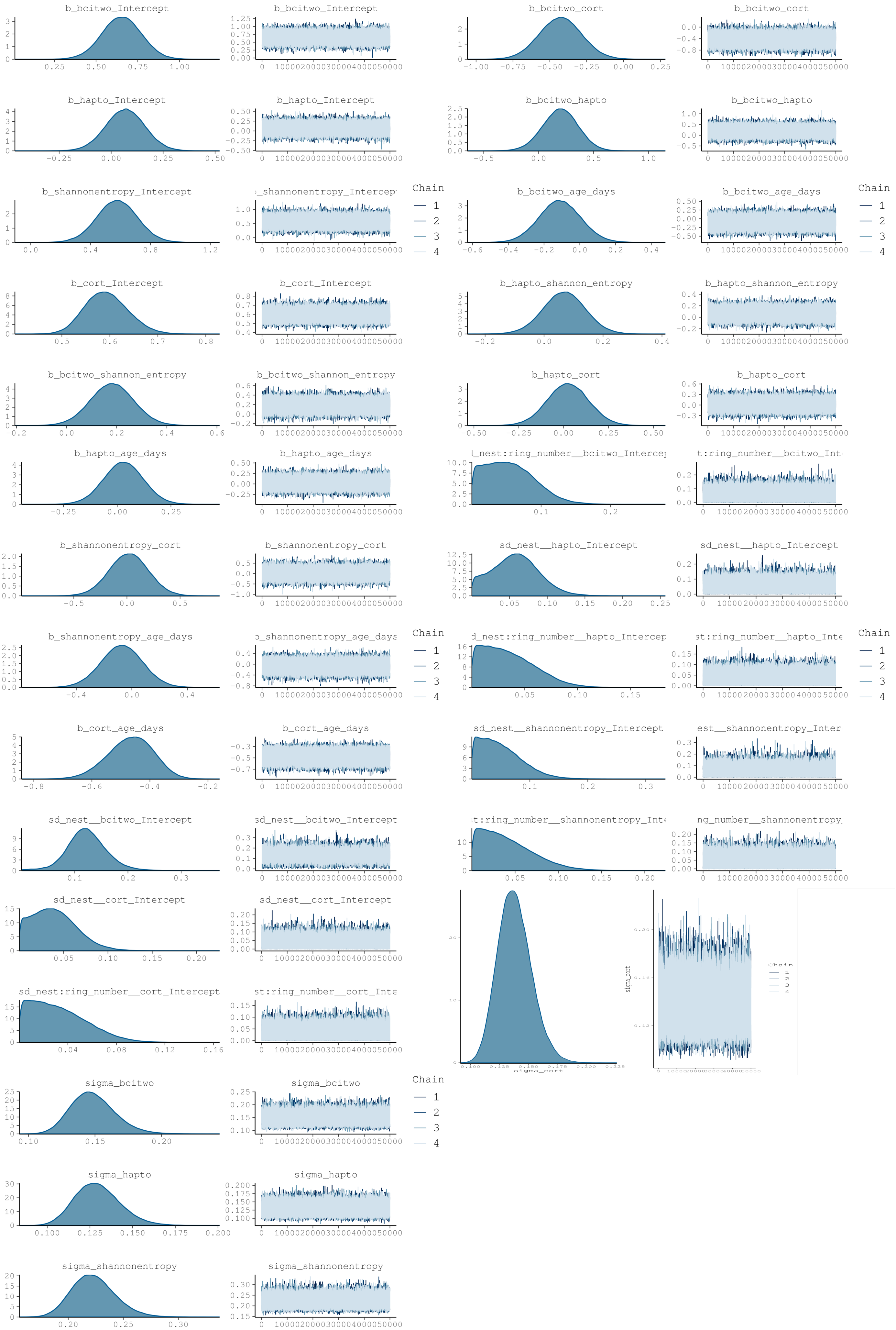
b_bcitwo_Intercept
b_bcitwo_cort
b_hapto_Intercept
b_bcitwo_hapto
b_shannonentropy_Intercept
b_bcitwo_age_days
Chain
1
2
3
4
b_cort_Intercept
b_hapto_shannon_entropy
b_bcitwo_shannon_entropy
b_hapto_cort
b_hapto_age_days
sd_nest__hapto_Intercept
b_shannonentropy_cort
b_shannonentropy_age_days
b_cort_age_days
sd_nest__shannonentropy_Intercept
sd_nest__bcitwo_Intercept
sd_nest__cort_Intercept
sd_nest:ring_number__cort_Intercept
sigma_cort
sigma_bcitwo
sigma_hapto
sigma_shannonentropy

##### 3.3 Compare distribution of response variable to distributions of predicted response variable

```
#Posterior predictive checks (one by one)
distribution_shannon_path1 <- pp_check(model_hapto_shannon, resp="bcitwo", ndraws=200)
distribution_shannon_path2 <- pp_check(model_hapto_shannon, resp="hapto", ndraws=200)
distribution_shannon_path3 <- pp_check(model_hapto_shannon, resp="shannonentropy", ndraws=200)
distribution_shannon_path4 <- pp_check(model_hapto_shannon, resp="cort", ndraws=200)

#Loop to save all distributions plot
responses <- c("bcitwo", "hapto", "shannonentropy", "cort")
response_names <- c("bci", "immune", "shannon", "cort")
for (i in seq_along(responses)) {
  pp_check_plot <- pp_check(model_hapto_shannon, resp = responses[i], ndraws = 200)
  filename <- paste0("distribution_shannon_", response_names[i])
  ggsave(filename = paste0(filename, ".svg"), plot = pp_check_plot, device = "svg", width = 8,
height = 10)
}
```

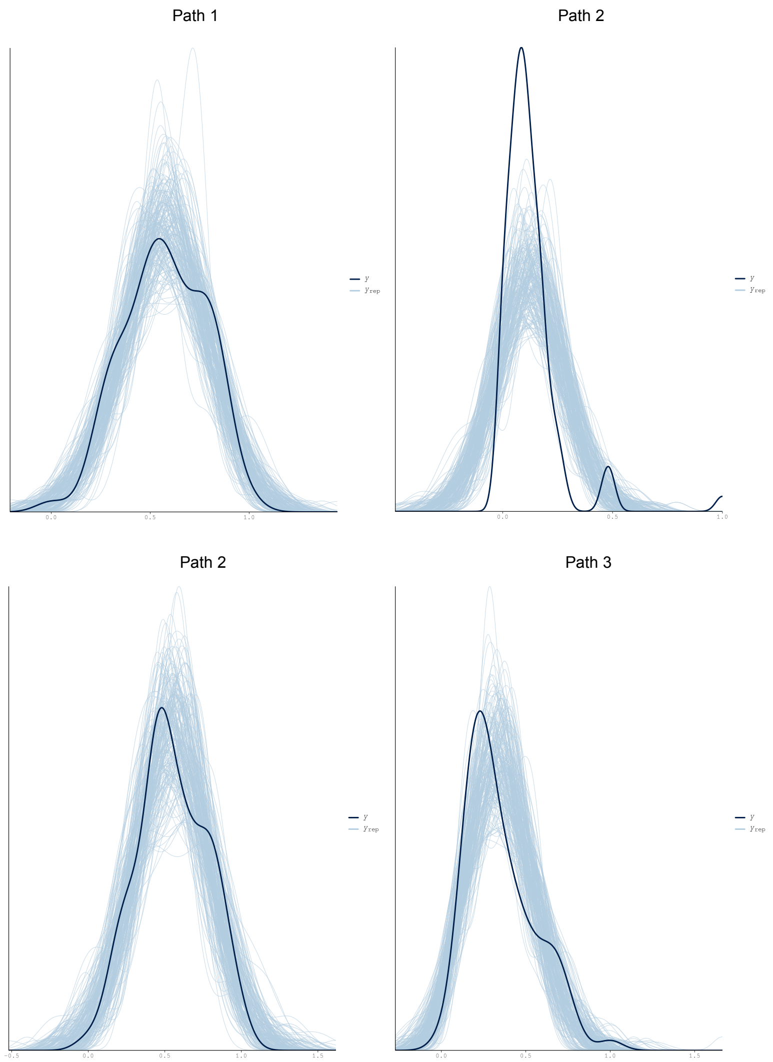

##### 3.4 Plot model posterior and credible intervals

```
#Plot Model effects
plot1 <-mcmc_plot(model_hapto_shannon, type = "intervals",prob_outer=0.95, prob=0.95,
                  variable =c("b_bcitwo_shannon_entropy", "b_bcitwo_cort", "b_bcitwo_hapto",
"b_bcitwo_age_days",
                              "b_hapto_shannon_entropy", "b_hapto_cort", "b_hapto_age_days",
                              "b_shannonentropy_cort", "b_shannonentropy_age_days",
                              "b_cort_age_days"))

plot1 <- plot1 + theme_classic() + geom_vline(xintercept = 0, linetype="dotted", color="blue")+
theme(axis.text.x = element_text(size = 16),  # Adjust the size as needed
axis.text.y = element_text(size = 16))+
theme(text = element_text(family = "Arial"))

ggsave(filename="16s_effect_sizes_shannon.svg", plot=plot1, device = "svg", width = 8, height =
10)
```

#### 4. Model diagnostics - Faith PD

##### 4.1 Model summary

```
#Model summary
summary_faith<- summary(model_hapto_faith)

#Bayes R2
R2m_faith <- bayes_R2(model_hapto_faith,re_formula=NA)
R2c_faith <- bayes_R2(model_hapto_faith)
```

##### 4.2 Model diagnostics

```
# Model diagnostics
diagnostic_faith <- plot(model_hapto_faith)

#Loop to save all diagnostic plots
diagnostic_plots <- list()
for (i in 1:length(diagnostic_faith)) {
  diagnostic_plots[[i]] <- diagnostic_faith[[i]]
  filename <- paste0("diagnostic", i, "_faith")
  ggsave(filename = paste0(filename, ".png"), plot = diagnostic_plots[[i]], device = "png",
dpi=200)
}
```

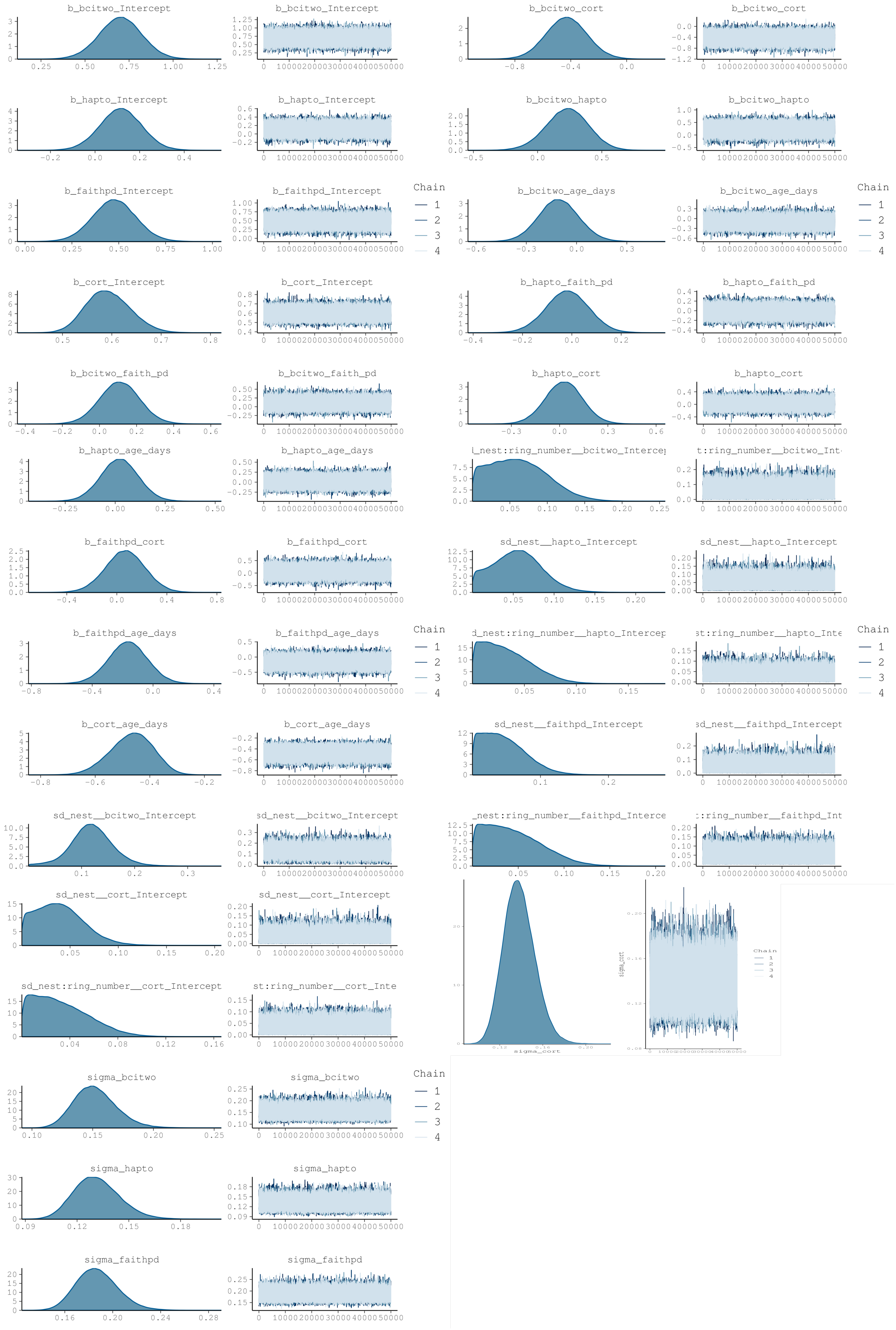
b_bcitwo_Intercept
b_bcitwo_cort
b_hapto_Intercept
b_bcitwo_hapto
b_faithpd_Intercept
b_bcitwo_age_days
b_cort_Intercept
b_hapto_faith_pd
b_bcitwo_faith_pd
b_hapto_cort
b_hapto_age_days
d_nest:ring_number__bcitwo_Intercep
t:ring_number__bcitwo_Int
b_faithpd_cort
sd_nest__hapto_Intercept
b_faithpd_age_days
d_nest:ring_number__hapto_Intercep
st:ring_number__hapto_Inte
b_cort_age_days
sd_nest__faithpd_Intercept
sd_nest__bcitwo_Intercept
_nest:ring_number__faithpd_Interce
:ring_number__faithpd_Int
sd_nest__cort_Intercept
sd_nest:ring_number__cort_Intercept
st:ring_number__cort_Inte
sigma_cort
sigma_bcitwo
sigma_hapto
sigma_faithpd
Chain
1
2
3
4

##### 4.3 Compare distribution of response variable to distributions of predicted response variable

```
#Posterior predictive checks
#Loop to save all distributions plot
responses <- c("bcitwo", "hapto", "faithpd", "cort")
response_names <- c("bci", "immune", "faith", "cort")
for (i in seq_along(responses)) {
  pp_check_plot <- pp_check(model_hapto_shannon, resp = responses[i], ndraws = 200)
  filename <- paste0("distribution_faith_", response_names[i])
  ggsave(filename = paste0(filename, ".svg"), plot = pp_check_plot, device = "svg", width = 8,
height = 10)
}
```

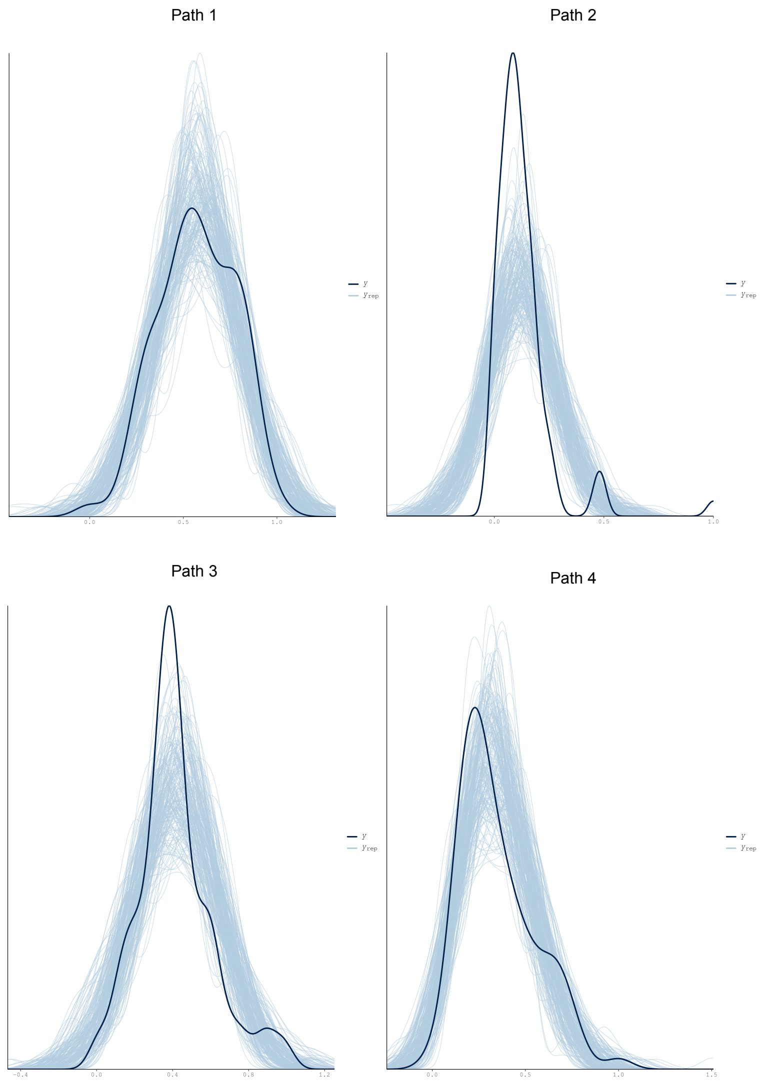


##### 4.4 Plot model posterior and credible intervals

```
#Plot Model effects
plot2 <-mcmc_plot(model_hapto_shannon, type = "intervals",prob_outer=0.95, prob=0.95,
                  variable =c("b_bcitwo_faith_pd", "b_bcitwo_cort", "b_bcitwo_hapto",
"b_bcitwo_age_days",
                               "b_hapto_faith_pd", "b_hapto_cort", "b_hapto_age_days",
                               "b_faithpd_cort", "b_faithpd_age_days",
                               "b_cort_age_days"))

plot2 <- plot2 + theme_classic() + geom_vline(xintercept = 0, linetype="dotted", color="blue")+
theme(axis.text.x = element_text(size = 16),  # Adjust the size as needed
axis.text.y = element_text(size = 16))+
theme(text = element_text(family = "Arial"))

ggsave(filename="16s_effect_sizes_faith.svg", plot=plot1, device = "svg", width = 8, height =
10)
```

### 5. Model diagnostics - N° of observed ASV's

#### 5.1 Model summary

```
#Model summary
summary_asv<- summary(model_hapto_asv)

#Bayes R2
R2m_asv <- bayes_R2(model_hapto_asv,re_formula=NA)
R2c_asv <- bayes_R2(model_hapto_asv)
```

#### 5.2 Model diagnostics

```
# Model diagnostics
diagnostic_asv <- plot(model_hapto_asv)

#Loop to save all diagnostic plots
diagnostic_plots <- list()
for (i in 1:length(diagnostic_asv)) {
  diagnostic_plots[[i]] <- diagnostic_asv[[i]]
  filename <- paste0("diagnostic", i, "_asv")
  ggsave(filename = paste0(filename, ".png"), plot = diagnostic_plots[[i]], device = "png",
dpi= 300)
}
```

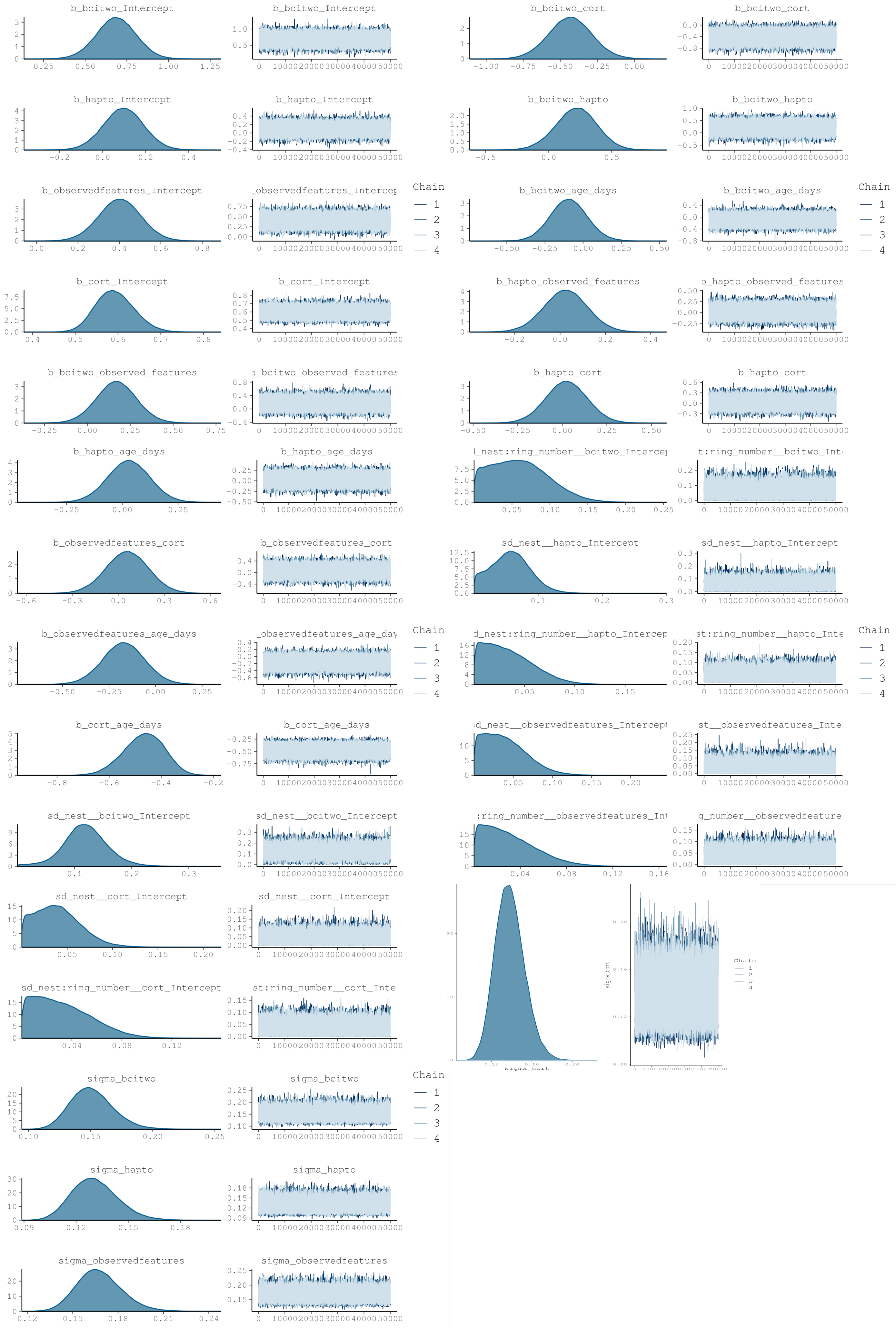
b_bcitwo_Intercept
b_bcitwo_Intercept
b_bcitwo_cort
b_bcitwo_cort
b_hapto_Intercept
b_hapto_Intercept
b_bcitwo_hapto
b_bcitwo_hapto
b_observedfeatures_Intercept
_observedfeatures_Intercep
Chain
1
2
3
4
b_bcitwo_age_days
b_bcitwo_age_days
Chain
1
2
3
4
b_cort_Intercept
b_cort_Intercept
b_hapto_observed_features
o_hapto_observed_features
b_bcitwo_observed_features
o_bcitwo_observed_features
b_hapto_cort
b_hapto_cort
b_hapto_age_days
b_hapto_age_days
l_nest:ring_number__bcitwo_Interce
t:ring_number__bcitwo_Int
b_observedfeatures_cort
b_observedfeatures_cort
sd_nest__hapto_Intercept
sd_nest__hapto_Intercept
b_observedfeatures_age_days
_observedfeatures_age_day
Chain
1
2
3
4
d_nest:ring_number__hapto_Intercep
st:ring_number__hapto_Inte
Chain
1
2
3
4
b_cort_age_days
b_cort_age_days
sd_nest__observedfeatures_Intercept
st__observedfeatures_Inte
sd_nest__bcitwo_Intercept
sd_nest__bcitwo_Intercept
:ring_number__observedfeatures_In
g_number__observedfeature
sd_nest__cort_Intercept
sd_nest__cort_Intercept
sigma_cort
sigma_cort
Chain
1
2
3
4
sd_nest:ring_number__cort_Intercept
st:ring_number__cort_Inte
sigma_bcitwo
sigma_bcitwo
Chain
1
2
3
4
sigma_hapto
sigma_hapto
sigma_observedfeatures
sigma_observedfeatures

##### 5.3 Compare distribution of response variable to distributions of predicted response variable

```
#Posterior predictive checks
#Loop to save all distributions plot
responses <- c("bcitwo", "hapto", "observedfeatures", "cort")
response_names <- c("bci", "immune", "asv", "cort")
for (i in seq_along(responses)) {
  pp_check_plot <- pp_check(model_hapto_asv, resp = responses[i], ndraws = 200)
  filename <- paste0("distribution_asv_", response_names[i])
  ggsave(filename = paste0(filename, ".svg"), plot = pp_check_plot, device = "svg", width = 8,
height = 10)
}
```

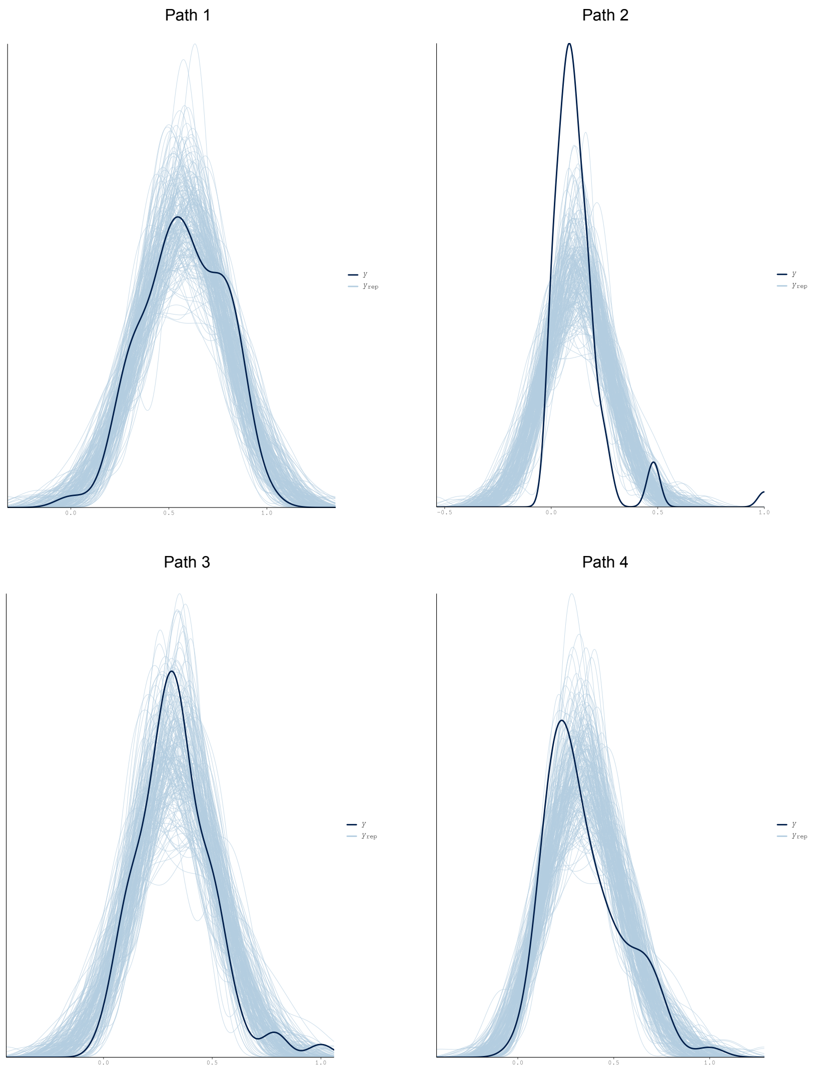


##### 5.4 Plot model posterior and credible intervals

```
#Plot Model effects
plot3 <-mcmc_plot(model_hapto_shannon, type = "intervals",prob_outer=0.95, prob=0.95,
                  variable =c("b_bcitwo_observed_features", "b_bcitwo_cort", "b_bcitwo_hapto",
"b_bcitwo_age_days",
                               "b_hapto_observed_features", "b_hapto_cort", "b_hapto_age_days",
                               "b_observedfeatures_cort", "b_observedfeatures_age_days",
                               "b_cort_age_days"))

plot3 <- plot3 + theme_classic() + geom_vline(xintercept = 0, linetype="dotted", color="blue")+
theme(axis.text.x = element_text(size = 16),  # Adjust the size as needed
axis.text.y = element_text(size = 16))+
theme(text = element_text(family = "Arial"))

ggsave(filename="16s_effect_sizes_asv.svg", plot=plot1, device = "svg", width = 8, height = 10)
```

#### B) 28S rRNA (eukaryotic microbiota) SEM analysis

##### 1. Define SEM for each diversity measurement

```
#Load the data
metadata <- readRDS("28s_metadata_immune.rds")

#scale all predictors to range between 0-1 if they are not already naturally on that scale

#define scaling function:
range.use <- function(x,min.use,max.use){ (x - min(x,na.rm=T))/(max(x,na.rm=T)-min(x,na.rm=T))
* (max.use - min.use) + min.use }
scalecols<-c("bci_two", "shannon_entropy", "faith_pd", "observed_features", "cort", "hapto",
"age_days")


for(i in 1:ncol(metadata[,which(colnames(metadata)%in%scalecols)])){
  metadata[,which(colnames(metadata)%in%scalecols)][,i]<-
range.use(metadata[,which(colnames(metadata)%in%scalecols)][,i],0,1)
}

# Define structural equation model paths

#Shannon
path1 <- bf(bci_two ~ shannon_entropy + cort + hapto+ age_days + (1|nest/ring_number))
path2 <- bf(hapto~  shannon_entropy + cort + age_days + (1|nest/ring_number))
path3 <- bf(shannon_entropy ~ cort + age_days + (1|nest/ring_number))
path4 <- bf(cort ~ age_days + (1|nest/ring_number)) + skew_normal()

sem_hapto_shannon <- path1 + path2 + path3 + path4

#Faith PD
path1 <- bf(bci_two ~ faith_pd + cort + hapto+ age_days + (1|nest/ring_number))
path2 <- bf(hapto~  faith_pd + cort + age_days + (1|nest/ring_number))
path3 <- bf(faith_pd ~ cort + age_days + (1|nest/ring_number))
path4 <- bf(cort ~ age_days + (1|nest/ring_number)) + skew_normal()

sem_hapto_faith <- path1 + path2 + path3 + path4
```

```
#N° of observed ASV's
path1 <- bf(bci_two ~ observed_features + cort + hapto+ age_days + (1|nest/ring_number))
path2 <- bf(hapto~  observed_features + cort + age_days + (1|nest/ring_number))
path3 <- bf(observed_features ~ cort + age_days + (1|nest/ring_number))
path4 <- bf(cort ~ age_days + (1|nest/ring_number)) + skew_normal()

sem_hapto_asv <- path1 + path2 + path3 + path4
```

#### 2. Run brms

```
ncores = detectCores()
options(mc.cores = parallel::detectCores())

#Shannon
model_hapto_shannon <-brm(sem_hapto_shannon + set_rescor(FALSE),
                         data = metadata,
                         warmup = 50000, iter = 100000,
                         control = list(adapt_delta = 0.99, max_treedepth = 15),
                         cores=ncores, chains=4, init=1000)

#Faith PD
model_hapto_faith <-brm(sem_hapto_faith + set_rescor(FALSE),
                         data = metadata,
                         warmup = 50000, iter = 100000,
                         control = list(adapt_delta = 0.99, max_treedepth = 15),
                         cores=ncores, chains=4, init=1000)

#N° of observed ASV's
model_hapto_asv <-brm(sem_hapto_asv + set_rescor(FALSE),
                         data = metadata,
                         warmup = 50000, iter = 100000,
                         control = list(adapt_delta = 0.99, max_treedepth = 15),
                         cores=ncores, chains=4, init=1000)
```

#### 3. Model Diagnostics - Shannon

##### 3.1 Model Summary

```
#Model summary
summary_shannon<- summary(model_hapto_shannon)

#Bayes R2
R2m_shannon <- bayes_R2(model_hapto_shannon,re_formula=NA)
R2c_shannon <- bayes_R2(model_hapto_shannon)
```

##### 3.2 Model diagnostics

```
# Model diagnostics
diagnostic_shannon <- plot(model_hapto_shannon)

#Loop to save all diagnostic plots
diagnostic_plots <- list()
for (i in 1:length(diagnostic_shannon)) {
  diagnostic_plots[[i]] <- diagnostic_shannon[[i]]
  filename <- paste0("diagnostic", i, "_shannon")
  ggsave(filename = paste0(filename, ".png"), plot = diagnostic_plots[[i]], device = "png",
dpi=300)
}
```

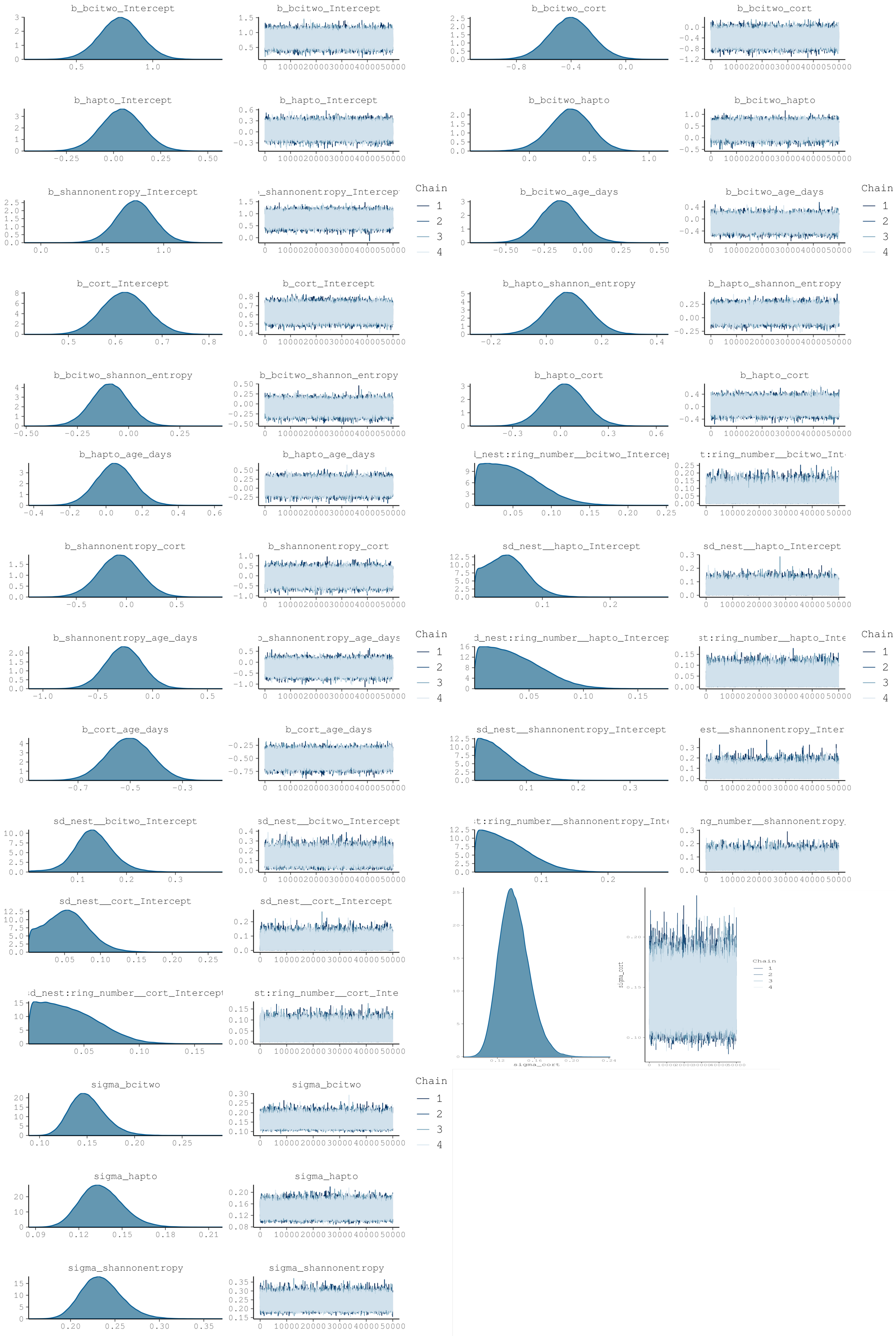
b_bcitwo_Intercept
b_bcitwo_cort
b_hapto_Intercept
b_bcitwo_hapto
b_shannonentropy_Intercept
Chain
1
2
3
4
b_bcitwo_age_days
b_cort_Intercept
b_hapto_shannon_entropy
b_bcitwo_shannon_entropy
b_hapto_cort
b_hapto_age_days
b_shannonentropy_cort
sd_nest__hapto_Intercept
b_shannonentropy_age_days
b_cort_age_days
sd_nest__shannonentropy_Intercept
sd_nest__bcitwo_Intercept
sd_nest__cort_Intercept
sigma_cort
sigma_bcitwo
sigma_hapto
sigma_shannonentropy

##### 3.3 Compare distribution of response variable to distributions of predicted response variable

```
#Posterior predictive checks (one by one)
distribution_shannon_path1 <- pp_check(model_hapto_shannon, resp="bcitwo", ndraws=200)
distribution_shannon_path2 <- pp_check(model_hapto_shannon, resp="hapto", ndraws=200)
distribution_shannon_path3 <- pp_check(model_hapto_shannon, resp="shannonentropy", ndraws=200)
distribution_shannon_path4 <- pp_check(model_hapto_shannon, resp="cort", ndraws=200)

#Loop to save all distributions plot
responses <- c("bcitwo", "hapto", "shannonentropy", "cort")
response_names <- c("bci", "immune", "shannon", "cort")
for (i in seq_along(responses)) {
  pp_check_plot <- pp_check(model_hapto_shannon, resp = responses[i], ndraws = 200)
  filename <- paste0("distribution_shannon_", response_names[i])
  ggsave(filename = paste0(filename, ".svg"), plot = pp_check_plot, device = "svg", width = 8,
height = 10)
}
```

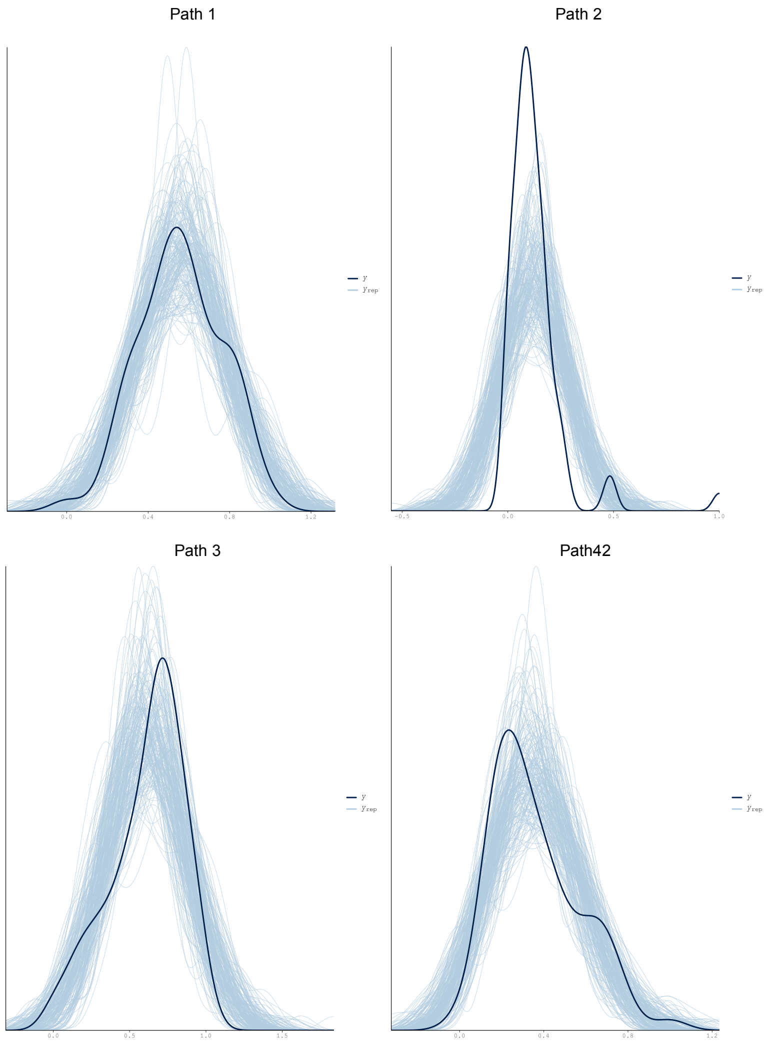

##### 3.4 Plot model posterior and credible intervals

```
#Plot Model effects
plot4 <-mcmc_plot(model_hapto_shannon, type = "intervals",prob_outer=0.95, prob=0.95,
                  variable =c("b_bcitwo_shannon_entropy", "b_bcitwo_cort", "b_bcitwo_hapto",
"b_bcitwo_age_days",
                               "b_hapto_shannon_entropy", "b_hapto_cort", "b_hapto_age_days",
                               "b_shannonentropy_cort", "b_shannonentropy_age_days",
                               "b_cort_age_days"))

plot4 <- plot4 + theme_classic() + geom_vline(xintercept = 0, linetype="dotted", color="blue")+
theme(axis.text.x = element_text(size = 16),  # Adjust the size as needed
axis.text.y = element_text(size = 16))+
theme(text = element_text(family = "Arial"))

ggsave(filename="16s_effect_sizes_shannon.svg", plot=plot4, device = "svg", width = 8, height =
10)
```

#### 4. Model diagnostics - Faith PD

##### 4.1 Model summary

```
#Model summary
summary_faith<- summary(model_hapto_faith)

#Bayes R2
R2m_faith <- bayes_R2(model_hapto_faith,re_formula=NA)
R2c_faith <- bayes_R2(model_hapto_faith)
```

##### 4.2 Model diagnostics

```
# Model diagnostics
diagnostic_faith <- plot(model_hapto_faith)

#Loop to save all diagnostic plots
diagnostic_plots <- list()
for (i in 1:length(diagnostic_faith)) {
  diagnostic_plots[[i]] <- diagnostic_faith[[i]]
  filename <- paste0("diagnostic", i, "_faith")
  ggsave(filename = paste0(filename, ".png"), plot = diagnostic_plots[[i]], device = "png",
dpi=200)
}
```

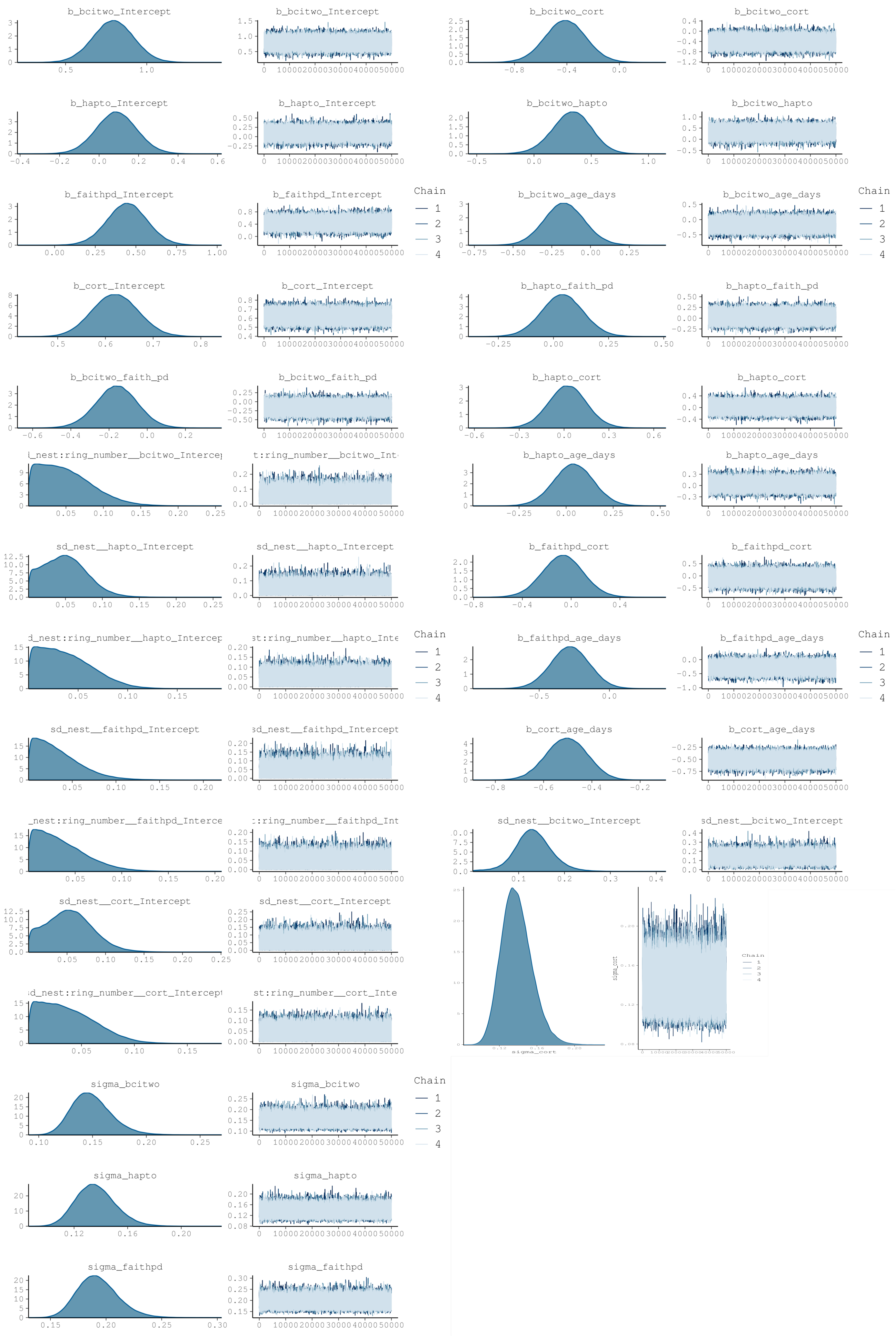

b_bcitwo_Intercept
b_bcitwo_cort
b_hapto_Intercept
b_bcitwo_hapto
b_faithpd_Intercept
b_bcitwo_age_days
Chain
1
2
3
4
b_cort_Intercept
b_hapto_faith_pd
b_bcitwo_faith_pd
b_hapto_cort
b_hapto_age_days
sd_nest__hapto_Intercept
b_faithpd_cort
b_faithpd_age_days
sd_nest__faithpd_Intercept
b_cort_age_days
sd_nest__bcitwo_Intercept
sd_nest__cort_Intercept
sigma_cort
sigma_bcitwo
sigma_hapto
sigma_faithpd

#### 4.3 Compare distribution of response variable to distributions of predicted response variable

```
#Posterior predictive checks
#Loop to save all distributions plot
responses <- c("bcitwo", "hapto", "faithpd", "cort")
response_names <- c("bci", "immune", "faith", "cort")
for (i in seq_along(responses)) {
  pp_check_plot <- pp_check(model_hapto_faith, resp = responses[i], ndraws = 200)
  filename <- paste0("distribution_faith_", response_names[i])
  ggsave(filename = paste0(filename, ".svg"), plot = pp_check_plot, device = "svg", width = 8,
height = 10)
}
```

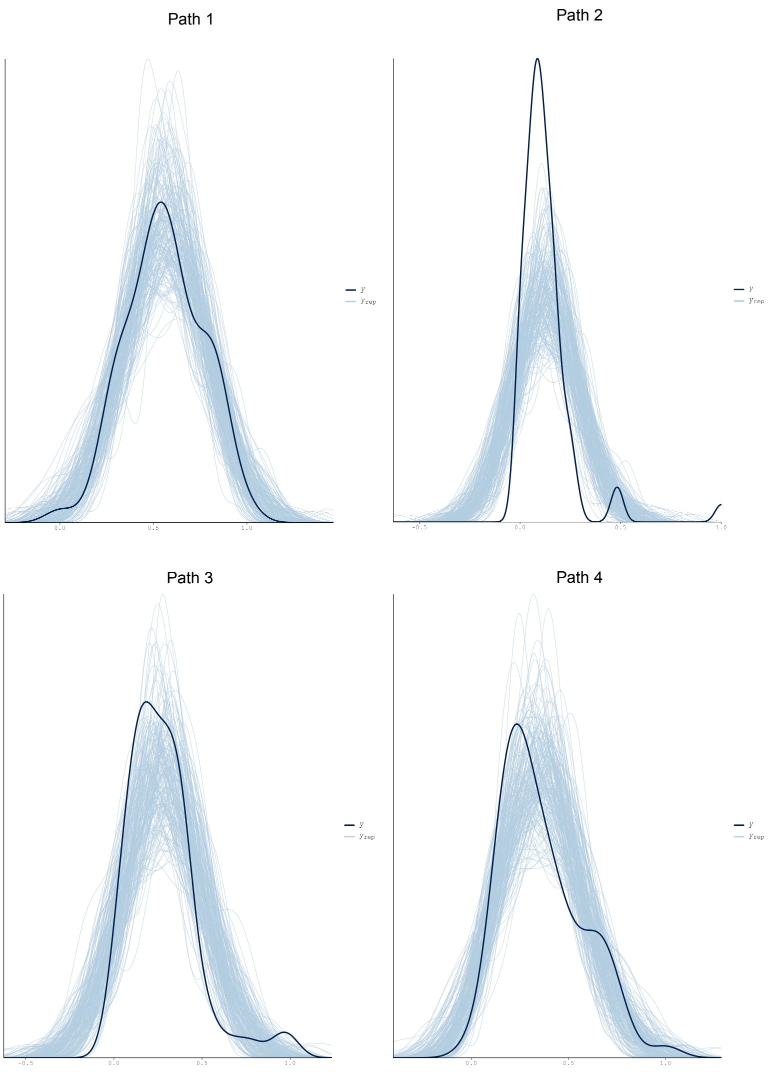


#### 4.4 Plot model posterior and credible intervals

```
#Plot Model effects
plot5 <-mcmc_plot(model_hapto_shannon, type = "intervals",prob_outer=0.95, prob=0.95,
                  variable =c("b_bcitwo_faith_pd", "b_bcitwo_cort", "b_bcitwo_hapto",
"b_bcitwo_age_days",
                               "b_hapto_faith_pd", "b_hapto_cort", "b_hapto_age_days",
                               "b_faithpd_cort", "b_faithpd_age_days",
                               "b_cort_age_days"))

plot5 <- plot5 + theme_classic() + geom_vline(xintercept = 0, linetype="dotted", color="blue")+
theme(axis.text.x = element_text(size = 16),  # Adjust the size as needed
axis.text.y = element_text(size = 16))+
theme(text = element_text(family = "Arial"))

ggsave(filename="16s_effect_sizes_faith.svg", plot=plot5, device = "svg", width = 8, height =
10)
```

### 5. Model diagnostics - N° of observed ASV's

#### 5.1 Model summary

```
#Model summary
summary_asv<- summary(model_hapto_asv)

#Bayes R2
R2m_asv <- bayes_R2(model_hapto_asv,re_formula=NA)
R2c_asv <- bayes_R2(model_hapto_asv)
```

#### 5.2 Model diagnostics

```
# Model diagnostics
diagnostic_asv <- plot(model_hapto_asv)

#Loop to save all diagnostic plots
diagnostic_plots <- list()
for (i in 1:length(diagnostic_asv)) {
  diagnostic_plots[[i]] <- diagnostic_asv[[i]]
  filename <- paste0("diagnostic", i, "_asv")
  ggsave(filename = paste0(filename, ".png"), plot = diagnostic_plots[[i]], device = "png",
dpi= 300)
}
```

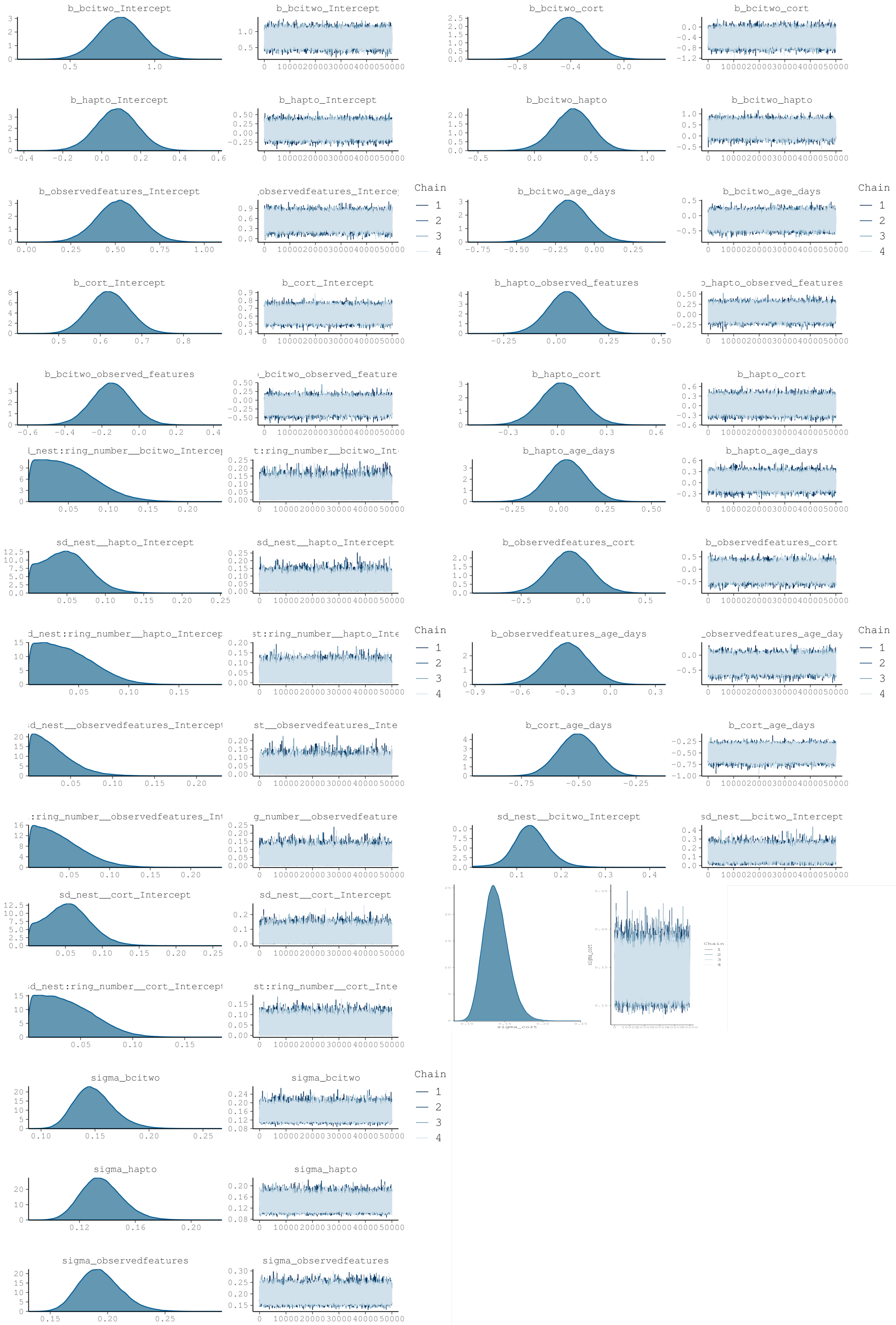
b_bcitwo_Intercept
b_bcitwo_Intercept
b_bcitwo_cort
b_bcitwo_cort
b_hapto_Intercept
b_hapto_Intercept
b_bcitwo_hapto
b_bcitwo_hapto
b_observedfeatures_Intercept
observedfeatures_Interce
Chain
1
2
3
4
b_bcitwo_age_days
b_bcitwo_age_days
Chain
1
2
3
4
b_cort_Intercept
b_cort_Intercept
b_hapto_observed_features
b_hapto_observed_features
b_bcitwo_observed_features
b_bcitwo_observed_feature
b_hapto_cort
b_hapto_cort
_nest:ring_number__bcitwo_Interce
t:ring_number__bcitwo_Int
b_hapto_age_days
b_hapto_age_days
sd_nest__hapto_Intercept
sd_nest__hapto_Intercept
b_observedfeatures_cort
b_observedfeatures_cort
d_nest:ring_number__hapto_Intercep
st:ring_number__hapto_Inte
Chain
1
2
3
4
b_observedfeatures_age_days
_observedfeatures_age_day
Chain
1
2
3
4
sd_nest__observedfeatures_Intercept
st__observedfeatures_Inte
b_cort_age_days
b_cort_age_days
:ring_number__observedfeatures_Int
g_number__observedfeature
sd_nest__bcitwo_Intercept
sd_nest__bcitwo_Intercept
sd_nest__cort_Intercept
sd_nest__cort_Intercept
d_nest:ring_number__cort_Intercept
st:ring_number__cort_Inte
Chain
1
2
3
4
sigma_bcitwo
sigma_bcitwo
sigma_hapto
sigma_hapto
sigma_observedfeatures
sigma_observedfeatures

##### 5.3 Compare distribution of response variable to distributions of predicted response variable

```
#Posterior predictive checks
#Loop to save all distributions plot
responses <- c("bcitwo", "hapto", "observedfeatures", "cort")
response_names <- c("bci", "immune", "asv", "cort")
for (i in seq_along(responses)) {
  pp_check_plot <- pp_check(model_hapto_asv, resp = responses[i], ndraws = 200)
  filename <- paste0("distribution_asv_", response_names[i])
  ggsave(filename = paste0(filename, ".svg"), plot = pp_check_plot, device = "svg", width = 8,
height = 10)
}
```

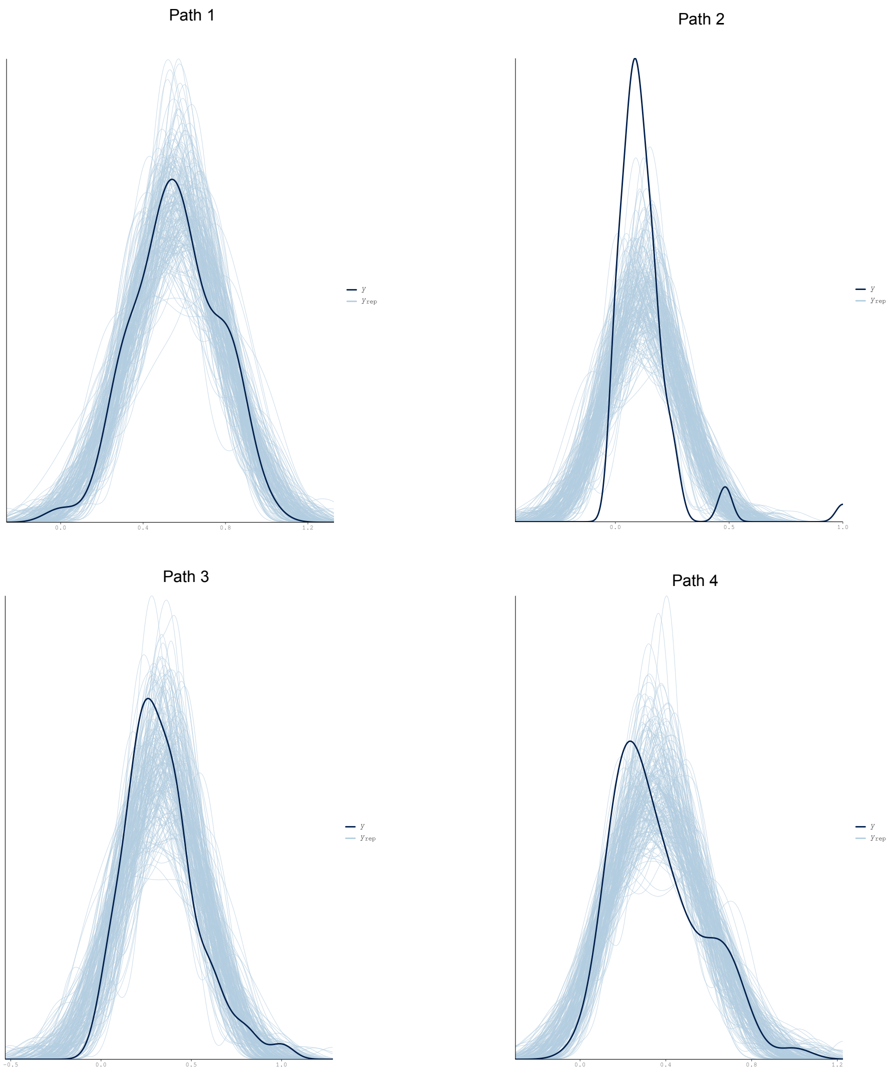


##### 5.4 Plot model posterior and credible intervals

```
#Plot Model effects
plot2 <-mcmc_plot(model_hapto_shannon, type = "intervals",prob_outer=0.95, prob=0.95,
                  variable =c("b_bcitwo_faith_pd", "b_bcitwo_cort", "b_bcitwo_hapto",
"b_bcitwo_age_days",
                               "b_hapto_faith_pd", "b_hapto_cort", "b_hapto_age_days",
                               "b_faithpd_cort", "b_faithpd_age_days",
                               "b_cort_age_days"))

plot2 <- plot2 + theme_classic() + geom_vline(xintercept = 0, linetype="dotted", color="blue")+
theme(axis.text.x = element_text(size = 16),  # Adjust the size as needed
axis.text.y = element_text(size = 16))+
theme(text = element_text(family = "Arial"))

ggsave(filename="16s_effect_sizes_faith.svg", plot=plot1, device = "svg", width = 8, height =
10)
```
