## Supplementary material for "The gut microbiota-immune-brain axis in a wild vertebrate: dynamic interactions and health impacts": DAA

### Differential Abundant Analysis with ANCOM-BC2

Pipeline adapted from https://bioconductor.org/packages/release/bioc/vignettes/ANCOMBC/inst/doc/ANCOMBC2.html

#### Table of Contents


**Differential Abundant Analysis with ANCOM-BC2**
- Table of Contents

**A) DAA 16s rRNA**
- 1. Read in the data
- 2. Run ANCOM-BC2
  - 2.1 ANCOM-BC2 at ASV level
    - 2.1.1 ANCOM-BC2 primary analysis
    - 2.1.2 Sensitivity scores
    - 2.2.3 Plot ANCOM-BC2 results

**B) DAA 28s rRNA**
- 1. Read in the data
- 2. Run ANCOM-BC2
  - 2.1 ANCOM-BC2 at ASV level
    - 2.1.1 ANCOM-BC2 primary analysis
    - 2.1.2 Sensitivity scores
    - 2.2.3 Plot ANCOM-BC2 results


### A) DAA 16s rRNA

#### 1. Read in the data

```
# Load libraries
library(phyloseq)
library(tidyverse)
library(janitor)
library(microbiome)
library(ANCOMBC)
library(ggrepel)

# read in the phyloseq object

ps_immune <- readRDS("ps_immune.rds")
```

#### 2. Run ANCOM-BC2

##### 2.1 ANCOM-BC2 at ASV level

```
# Diferential abundace analysis on final model.

set.seed(123)
output_immunity = ancombc2(data = ps_immune, assay_name = "counts", tax_level = NULL,
                        fix_formula = "std_bci_two + std_cort + pred_immunity + std_age",
rand_formula = "(1|nest/ring_number)",
                        p_adj_method = "holm", pseudo = 0, pseudo_sens = TRUE,
                        prv_cut = 0.10, lib_cut = 0, s0_perc = 0.05,
                        #group = NULL, struc_zero = TRUE, neg_lb = TRUE,
                        alpha = 0.05, n_cl = 2, verbose = TRUE,
                        #global = TRUE, pairwise = TRUE, dunnet = TRUE, trend = TRUE,
                        iter_control = list(tol = 1e-2, max_iter = 50, verbose = TRUE),
                        em_control = list(tol = 1e-5, max_iter = 100),
                        lme_control = lme4::lmerControl(optimizer ="Nelder_Mead"),
                        mdfdr_control = list(fwer_ctrl_method = "holm", B = 100),
                        trend_control = list(contrast = list(matrix(c(1, 0, -1, 1),
                        nrow = 2, byrow = TRUE), matrix(c(-1, 0, 1, -1), nrow = 2,
                        byrow = TRUE)), node = list(2, 2), solver = "ECOS", B = 1000))

#run the same model for haptoglobin
output_hapto = ancombc2(data = ps_immune, assay_name = "counts", tax_level = NULL,
                        fix_formula = "std_bci_two + std_cort + std_hapto + std_age",
rand_formula = "(1|nest/ring_number)",
                        p_adj_method = "holm", pseudo = 0, pseudo_sens = TRUE,
                        prv_cut = 0.10, lib_cut = 0, s0_perc = 0.05,
                        #group = NULL, struc_zero = TRUE, neg_lb = TRUE,
                        alpha = 0.05, n_cl = 2, verbose = TRUE,
                        #global = TRUE, pairwise = TRUE, dunnet = TRUE, trend = TRUE,
                        iter_control = list(tol = 1e-2, max_iter = 50, verbose = TRUE),
                        em_control = list(tol = 1e-5, max_iter = 100),
                        lme_control = lme4::lmerControl(optimizer ="Nelder_Mead"),
                        mdfdr_control = list(fwer_ctrl_method = "holm", B = 100),
                        trend_control = list(contrast = list(matrix(c(1, 0, -1, 1),
                        nrow = 2, byrow = TRUE), matrix(c(-1, 0, 1, -1), nrow = 2,
                        byrow = TRUE)), node = list(2, 2), solver = "ECOS", B = 1000))
```

###### 2.1.1 ANCOM-BC2 primary analysis

```
res_prim_immunity = output_immunity$res

res_prim_hapto = output_hapto$res

 # Check taxonomy of DAA taxa
 DAA_hapto <- c("86f88dec7777f0258548b6bdb3ea53c9")
 DAA_hapto_ps <- prune_taxa(taxa_names(ps_immune) %in% DAA_hapto, ps_immune)
 DAA_hapto_ps@tax_table
                                   #Kingdom      Phylum       Class         Order
Family           Genus
 #86f88dec7777f0258548b6bdb3ea53c9 "Bacteria" "Bacteroidota" "Bacteroidia" "Bacteroidales"
"Prevotellaceae" "Alloprevotella"
#Species
#"Alloprevotella_rava"
```

**No deferentially abundant ASVs found for models with latente variable Immunity**

**One deferentially abundant ASVs (with age) found for models with haptoglobin did not pass the sensitivity analysis**

##### 2.1.2 Sensitivity scores

ANCOM-BC2 uses a sensitivity analysis to assess the impact of different pseudo-counts on zero counts for each taxon. The sensitivity score is determined by performing linear regression models on the bias-corrected log abundance table using various pseudo-counts and calculating the proportion of times the p-value exceeds the significance level (alpha). This helps identify taxa that are not sensitive to the pseudo-count addition, ensuring robustness in the analysis.

```
tab_sens_immunity = output_immunity$ss_tab

tab_sens_hapto = output_hapto$ss_tab
```

##### 2.2.3 Plot ANCOM-BC2 results

```
# Volcano plots - immunity
volc_immunity_bci <- ggplot(data=res_prim_immunity, aes(x=lfc_std_bci_two, y=-
log10(p_std_bci_two), col=diff_std_bci_two)) + geom_point(size=3) +
   geom_text_repel(aes(label = ifelse(diff_std_bci_two, taxon, ""))) +
   theme_classic()+
  labs(x = "Lfc std. BCI", y = "-Log10 (p std. BCI)") +
  theme(axis.text.x = element_text(size = 16),  # Adjust the size as needed
        axis.text.y = element_text(size = 16))+
  theme(axis.title.x = element_text(size = 16),  # Adjust the size as needed
        axis.title.y = element_text(size = 16))+
  theme(axis.title.x = element_text(margin = margin(t = 13)))+
  theme(axis.title.y = element_text(margin = margin(r = 14)))+
  theme(legend.position = "none")+
  theme(text = element_text(family = "Arial"))
 volc_immunity_bci

 volc_immunity_immunity <- ggplot(data=res_prim_immunity, aes(x=lfc_pred_immunity, y=-
log10(p_pred_immunity), col=diff_pred_immunity)) + geom_point(size=3) +
   geom_text_repel(aes(label = ifelse(diff_pred_immunity, taxon, ""))) +
   theme_classic()+
   labs(x = "Lfc Immunity", y = "-Log10 (p Immunity)") +
   theme(axis.text.x = element_text(size = 16),  # Adjust the size as needed
         axis.text.y = element_text(size = 16))+
   theme(axis.title.x = element_text(size = 16),  # Adjust the size as needed
         axis.title.y = element_text(size = 16))+
   theme(axis.title.x = element_text(margin = margin(t = 13)))+
   theme(axis.title.y = element_text(margin = margin(r = 14)))+
   theme(legend.position = "none")+
   theme(text = element_text(family = "Arial"))
 volc_immunity_immunity

 volc_immunity_cort <- ggplot(data=res_prim_immunity, aes(x=lfc_std_cort, y=-log10(p_std_cort),
col=diff_std_cort)) + geom_point(size=3) +
   geom_text_repel(aes(label = ifelse(diff_std_cort, taxon, ""))) +
   theme_classic()+
   labs(x = "Lfc std. CORT", y = "-Log10 (p std. CORT)") +
   theme(axis.text.x = element_text(size = 16),  # Adjust the size as needed
         axis.text.y = element_text(size = 16))+
   theme(axis.title.x = element_text(size = 16),  # Adjust the size as needed
```

```
        axis.title.y = element_text(size = 16))+
  theme(axis.title.x = element_text(margin = margin(t = 13)))+
  theme(axis.title.y = element_text(margin = margin(r = 14)))+
  theme(legend.position = "none")+
  theme(text = element_text(family = "Arial"))
 volc_immunity_cort

 volc_immunity_age <- ggplot(data=res_prim_immunity, aes(x=lfc_std_age, y=-log10(p_std_age),
col=diff_std_age)) + geom_point(size=3) +
  geom_text_repel(aes(label = ifelse(diff_std_age, taxon, ""))) +
  theme_classic()+
  labs(x = "Lfc std. Age", y = "-Log10 (p std. Age)") +
  theme(axis.text.x = element_text(size = 16),  # Adjust the size as needed
        axis.text.y = element_text(size = 16))+
  theme(axis.title.x = element_text(size = 16),  # Adjust the size as needed
        axis.title.y = element_text(size = 16))+
  theme(axis.title.x = element_text(margin = margin(t = 13)))+
  theme(axis.title.y = element_text(margin = margin(r = 14)))+
  theme(text = element_text(family = "Arial"))
 volc_immunity_age


# Volcano plots - haptoglobin

volc_hapto_bci <- ggplot(data=res_prim_hapto, aes(x=lfc_std_bci_two, y=-log10(p_std_bci_two),
col=diff_std_bci_two)) + geom_point(size=3) +
  geom_text_repel(aes(label = ifelse(diff_std_bci_two, taxon, ""))) +
  theme_classic()+
 labs(x = "Lfc std. BCI", y = "-Log10 (p std. BCI)") +
 theme(axis.text.x = element_text(size = 16),  # Adjust the size as needed
       axis.text.y = element_text(size = 16))+
 theme(axis.title.x = element_text(size = 16),  # Adjust the size as needed
       axis.title.y = element_text(size = 16))+
 theme(axis.title.x = element_text(margin = margin(t = 13)))+
 theme(axis.title.y = element_text(margin = margin(r = 14)))+
 theme(legend.position = "none")+
 theme(text = element_text(family = "Arial"))
 volc_hapto_bci

 volc_hapto_hapto <- ggplot(data=res_prim_hapto, aes(x=lfc_std_hapto, y=-log10(p_std_hapto),
col=diff_std_hapto)) + geom_point(size=3) +
  geom_text_repel(aes(label = ifelse(diff_std_hapto, taxon, ""))) +
  theme_classic()+
  labs(x = "Lfc Haptoglobin", y = "-Log10 (p Haptoglobin)") +
  theme(axis.text.x = element_text(size = 16),  # Adjust the size as needed
        axis.text.y = element_text(size = 16))+
  theme(axis.title.x = element_text(size = 16),  # Adjust the size as needed
        axis.title.y = element_text(size = 16))+
  theme(axis.title.x = element_text(margin = margin(t = 13)))+
  theme(axis.title.y = element_text(margin = margin(r = 14)))+
  theme(legend.position = "none")+
  theme(text = element_text(family = "Arial"))
 volc_hapto_hapto

 volc_hapto_cort <- ggplot(data=res_prim_hapto, aes(x=lfc_std_cort, y=-log10(p_std_cort),
col=diff_std_cort)) + geom_point(size=3) +
  geom_text_repel(aes(label = ifelse(diff_std_cort, taxon, ""))) +
  theme_classic()+
  labs(x = "Lfc std. CORT", y = "-Log10 (p std. CORT)") +
  theme(axis.text.x = element_text(size = 16),  # Adjust the size as needed
```

```
        axis.text.y = element_text(size = 16))+
  theme(axis.title.x = element_text(size = 16),  # Adjust the size as needed
        axis.title.y = element_text(size = 16))+
  theme(axis.title.x = element_text(margin = margin(t = 13)))+
  theme(axis.title.y = element_text(margin = margin(r = 14)))+
  theme(legend.position = "none")+
  theme(text = element_text(family = "Arial"))
 volc_hapto_cort

 volc_hapto_age <- ggplot(data=res_prim_hapto, aes(x=lfc_std_age, y=-log10(p_std_age),
col=diff_std_age)) + geom_point(size=3) +
  geom_text_repel(aes(label = ifelse(diff_std_age, taxon, ""))) +
  theme_classic()+
  labs(x = "Lfc std. Age", y = "-Log10 (p std. Age)") +
  theme(axis.text.x = element_text(size = 16),  # Adjust the size as needed
        axis.text.y = element_text(size = 16))+
  theme(axis.title.x = element_text(size = 16),  # Adjust the size as needed
        axis.title.y = element_text(size = 16))+
  theme(axis.title.x = element_text(margin = margin(t = 13)))+
  theme(axis.title.y = element_text(margin = margin(r = 14)))+
  theme(text = element_text(family = "Arial"))
 volc_hapto_age
```

```
# Pseudo-count sensitivity analysis for age

df_age = res_prim_hapto %>% dplyr::select(taxon, ends_with("age")) # create a dataframe with
values only for age

# Sensitivity scores
tab_sens_age <- output_hapto$ss_tab

## Pseudo count sensitivity analysis for age
sens_age <- tab_sens_age %>%
  transmute(taxon, sens_age = std_age) %>%
  left_join(df_age, by = "taxon")
sens_age$diff_std_age = recode(sens_age$diff_std_age * 1,
                               `1` = "Significant",
                               `0` = "Nonsignificant")

fig_sens_age = sens_age %>%
  ggplot(aes(x = taxon, y = sens_age, color = diff_std_age)) +
  geom_point() +
  scale_color_brewer(palette = "Dark2", name = NULL) +
  labs(x = NULL, y = "Sensitivity Score") +
  theme_classic() +
  theme(axis.text.x = element_text(angle = 60, vjust = 0.5))
fig_sens_age
```

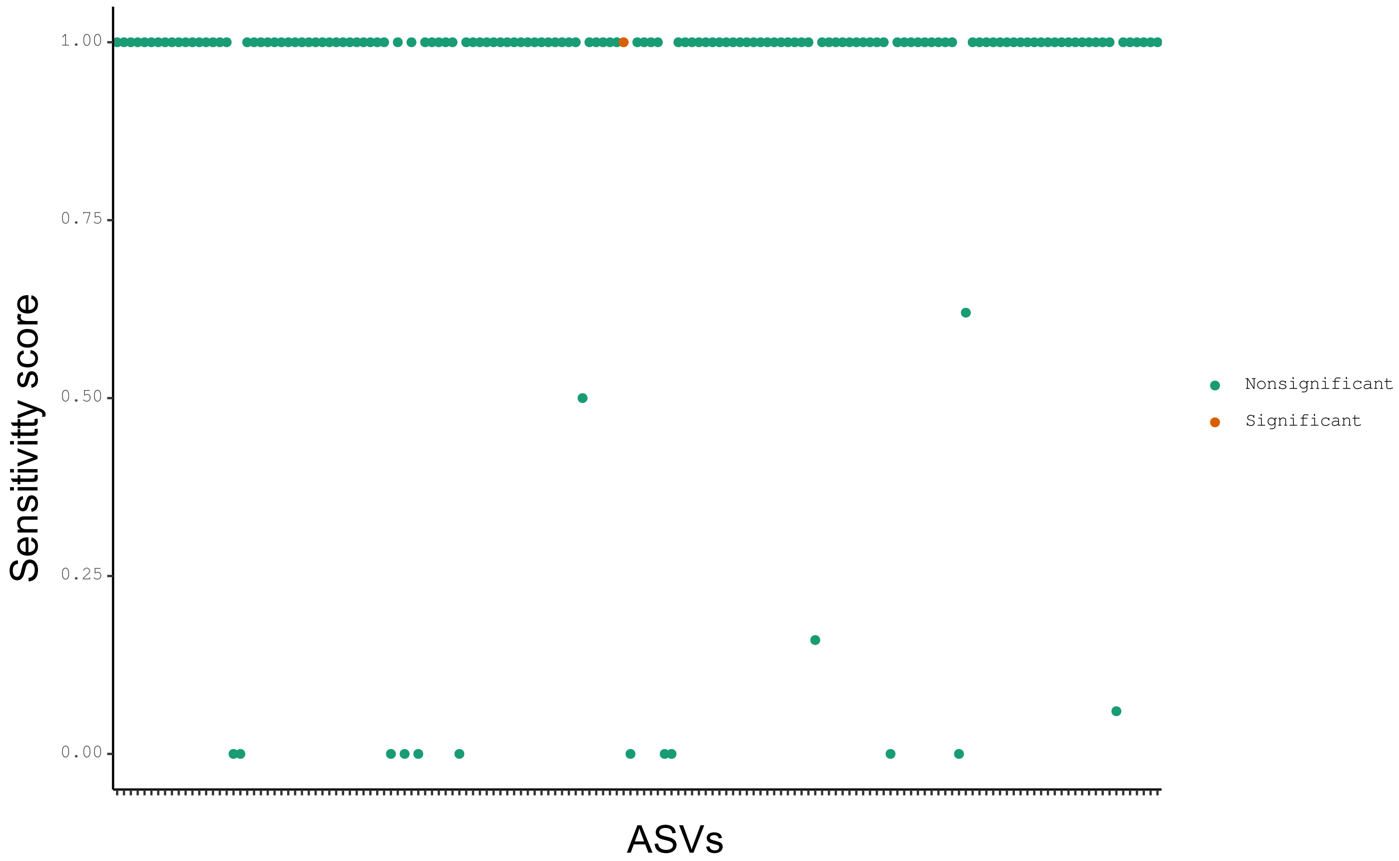


**For the co-variate of age, the significant taxa have high sensitivity scores.**

### B) DAA 28s rRNA

#### 1. Read in the data

```
# read in the phyloseq object
ps_immune <- readRDS("ps_immune.rds")
```

#### 2. Run ANCOM-BC2

##### 2.1 ANCOM-BC2 at ASV level

```
# Diferential abundace analysis on final model.

set.seed(123)
output_immunity = ancombc2(data = ps_immune, assay_name = "counts", tax_level = NULL,
                        fix_formula = "std_bci_two + std_cort + pred_immunity + std_age",
rand_formula = "(1|nest/ring_number)",
                        p_adj_method = "holm", pseudo = 0, pseudo_sens = TRUE,
                        prv_cut = 0.10, lib_cut = 0, s0_perc = 0.05,
                        #group = NULL, struc_zero = TRUE, neg_lb = TRUE,
                        alpha = 0.05, n_cl = 2, verbose = TRUE,
                        #global = TRUE, pairwise = TRUE, dunnet = TRUE, trend = TRUE,
```

```
                         iter_control = list(tol = 1e-2, max_iter = 50, verbose = TRUE),
                         em_control = list(tol = 1e-5, max_iter = 100),
                         lme_control = lme4::lmerControl(optimizer ="Nelder_Mead"),
                         mdfdr_control = list(fwer_ctrl_method = "holm", B = 100),
                         trend_control = list(contrast = list(matrix(c(1, 0, -1, 1),
                         nrow = 2, byrow = TRUE), matrix(c(-1, 0, 1, -1), nrow = 2,
                         byrow = TRUE)), node = list(2, 2), solver = "ECOS", B = 1000))

#run the same model for haptoglobin
output_hapto = ancombc2(data = ps_immune, assay_name = "counts", tax_level = NULL,
                         fix_formula = "std_bci_two + std_cort + std_hapto + std_age",
rand_formula = "(1|nest/ring_number)",
                         p_adj_method = "holm", pseudo = 0, pseudo_sens = TRUE,
                         prv_cut = 0.10, lib_cut = 0, s0_perc = 0.05,
                         #group = NULL, struc_zero = TRUE, neg_lb = TRUE,
                         alpha = 0.05, n_cl = 2, verbose = TRUE,
                         #global = TRUE, pairwise = TRUE, dunnet = TRUE, trend = TRUE,
                         iter_control = list(tol = 1e-2, max_iter = 50, verbose = TRUE),
                         em_control = list(tol = 1e-5, max_iter = 100),
                         lme_control = lme4::lmerControl(optimizer ="Nelder_Mead"),
                         mdfdr_control = list(fwer_ctrl_method = "holm", B = 100),
                         trend_control = list(contrast = list(matrix(c(1, 0, -1, 1),
                         nrow = 2, byrow = TRUE), matrix(c(-1, 0, 1, -1), nrow = 2,
                         byrow = TRUE)), node = list(2, 2), solver = "ECOS", B = 1000))
```

###### 2.1.1 ANCOM-BC2 primary analysis

```
res_prim_immunity = output_immunity$res

res_prim_hapto = output_hapto$res


# Check taxonomy of DAA taxa

DAA_bci <- c("ASV_201")
DAA_bci_ps <- prune_taxa(taxa_names(ps_immune) %in% DAA_bci, ps_immune)
DAA_bci_ps@tax_table

            #Kingdom        Phylum       Class              Order          Family         Genus
      Species
#ASV_201 "d__Eukaryota" "Ascomycota" "Dothideomycetes" "Dothideales" "Dothideales"
"Dothideales" "Hormonema_carpetanum"

DAA_age <- c("ASV_381")
DAA_age_ps <- prune_taxa(taxa_names(ps_immune) %in% DAA_age, ps_immune)
DAA_age_ps@tax_table

             #Kingdom        Phylum               Class              Order
Family               Genus            Species
#ASV_381"d__Eukaryota;p_Phragmoplastophyta;c_Phragmoplastophyta;o_Phragmoplastophyta;f_Phragmop
lastophyta;g_Phragmoplastophyta;s_Pinus_taeda"
```

**Two deferentially abundant ASVs co-vary with Age and BCI when the model incorporates the variable immunity.**
**ASV that co-varies with age did not pass sensitivity analysis**
**No differential abundant taxa found when modelling haptoglobin**

##### 2.1.2 Sensitivity scores

```
tab_sens_immunity = output_immunity$ss_tab

tab_sens_hapto = output_hapto$ss_tab
```

##### 2.2.3 Plot ANCOM-BC2 results

```
# Volcano plots - immunity
volc_immunity_bci <- ggplot(data=res_prim_immunity, aes(x=lfc_std_bci_two, y=-
log10(p_std_bci_two), col=diff_std_bci_two)) + geom_point(size=3) +
   geom_text_repel(aes(label = ifelse(diff_std_bci_two, taxon, ""))) +
   theme_classic()+
  labs(x = "Lfc std. BCI", y = "-Log10 (p std. BCI)") +
  theme(axis.text.x = element_text(size = 16),  # Adjust the size as needed
        axis.text.y = element_text(size = 16))+
  theme(axis.title.x = element_text(size = 16),  # Adjust the size as needed
        axis.title.y = element_text(size = 16))+
  theme(axis.title.x = element_text(margin = margin(t = 13)))+
  theme(axis.title.y = element_text(margin = margin(r = 14)))+
  theme(legend.position = "none")+
  theme(text = element_text(family = "Arial"))
 volc_immunity_bci

 volc_immunity_immunity <- ggplot(data=res_prim_immunity, aes(x=lfc_pred_immunity, y=-
log10(p_pred_immunity), col=diff_pred_immunity)) + geom_point(size=3) +
   geom_text_repel(aes(label = ifelse(diff_pred_immunity, taxon, ""))) +
   theme_classic()+
   labs(x = "Lfc Immunity", y = "-Log10 (p Immunity)") +
   theme(axis.text.x = element_text(size = 16),  # Adjust the size as needed
         axis.text.y = element_text(size = 16))+
   theme(axis.title.x = element_text(size = 16),  # Adjust the size as needed
         axis.title.y = element_text(size = 16))+
   theme(axis.title.x = element_text(margin = margin(t = 13)))+
   theme(axis.title.y = element_text(margin = margin(r = 14)))+
   theme(legend.position = "none")+
   theme(text = element_text(family = "Arial"))
 volc_immunity_immunity

 volc_immunity_cort <- ggplot(data=res_prim_immunity, aes(x=lfc_std_cort, y=-log10(p_std_cort),
col=diff_std_cort)) + geom_point(size=3) +
   geom_text_repel(aes(label = ifelse(diff_std_cort, taxon, ""))) +
   theme_classic()+
   labs(x = "Lfc std. CORT", y = "-Log10 (p std. CORT)") +
   theme(axis.text.x = element_text(size = 16),  # Adjust the size as needed
         axis.text.y = element_text(size = 16))+
   theme(axis.title.x = element_text(size = 16),  # Adjust the size as needed
         axis.title.y = element_text(size = 16))+
   theme(axis.title.x = element_text(margin = margin(t = 13)))+
   theme(axis.title.y = element_text(margin = margin(r = 14)))+
   theme(legend.position = "none")+
   theme(text = element_text(family = "Arial"))
 volc_immunity_cort

 volc_immunity_age <- ggplot(data=res_prim_immunity, aes(x=lfc_std_age, y=-log10(p_std_age),
col=diff_std_age)) + geom_point(size=3) +
   geom_text_repel(aes(label = ifelse(diff_std_age, taxon, ""))) +
```

```
   theme_classic()+
   labs(x = "Lfc std. Age", y = "-Log10 (p std. Age)") +
   theme(axis.text.x = element_text(size = 16),  # Adjust the size as needed
         axis.text.y = element_text(size = 16))+
   theme(axis.title.x = element_text(size = 16),  # Adjust the size as needed
         axis.title.y = element_text(size = 16))+
   theme(axis.title.x = element_text(margin = margin(t = 13)))+
   theme(axis.title.y = element_text(margin = margin(r = 14)))+
   theme(text = element_text(family = "Arial"))
 volc_immunity_age


# Volcano plots - haptoglobin

volc_hapto_bci <- ggplot(data=res_prim_hapto, aes(x=lfc_std_bci_two, y=-log10(p_std_bci_two),
col=diff_std_bci_two)) + geom_point(size=3) +
   geom_text_repel(aes(label = ifelse(diff_std_bci_two, taxon, ""))) +
   theme_classic()+
  labs(x = "Lfc std. BCI", y = "-Log10 (p std. BCI)") +
  theme(axis.text.x = element_text(size = 16),  # Adjust the size as needed
        axis.text.y = element_text(size = 16))+
  theme(axis.title.x = element_text(size = 16),  # Adjust the size as needed
        axis.title.y = element_text(size = 16))+
  theme(axis.title.x = element_text(margin = margin(t = 13)))+
  theme(axis.title.y = element_text(margin = margin(r = 14)))+
  theme(legend.position = "none")+
  theme(text = element_text(family = "Arial"))
 volc_hapto_bci

 volc_hapto_hapto <- ggplot(data=res_prim_hapto, aes(x=lfc_std_hapto, y=-log10(p_std_hapto),
col=diff_std_hapto)) + geom_point(size=3) +
   geom_text_repel(aes(label = ifelse(diff_std_hapto, taxon, ""))) +
   theme_classic()+
   labs(x = "Lfc Haptoglobin", y = "-Log10 (p Haptoglobin)") +
   theme(axis.text.x = element_text(size = 16),  # Adjust the size as needed
         axis.text.y = element_text(size = 16))+
   theme(axis.title.x = element_text(size = 16),  # Adjust the size as needed
         axis.title.y = element_text(size = 16))+
   theme(axis.title.x = element_text(margin = margin(t = 13)))+
   theme(axis.title.y = element_text(margin = margin(r = 14)))+
   theme(legend.position = "none")+
   theme(text = element_text(family = "Arial"))
 volc_hapto_hapto

 volc_hapto_cort <- ggplot(data=res_prim_hapto, aes(x=lfc_std_cort, y=-log10(p_std_cort),
col=diff_std_cort)) + geom_point(size=3) +
   geom_text_repel(aes(label = ifelse(diff_std_cort, taxon, ""))) +
   theme_classic()+
   labs(x = "Lfc std. CORT", y = "-Log10 (p std. CORT)") +
   theme(axis.text.x = element_text(size = 16),  # Adjust the size as needed
         axis.text.y = element_text(size = 16))+
   theme(axis.title.x = element_text(size = 16),  # Adjust the size as needed
         axis.title.y = element_text(size = 16))+
   theme(axis.title.x = element_text(margin = margin(t = 13)))+
   theme(axis.title.y = element_text(margin = margin(r = 14)))+
   theme(legend.position = "none")+
   theme(text = element_text(family = "Arial"))
 volc_hapto_cort
```

```
 volc_hapto_age <- ggplot(data=res_prim_hapto, aes(x=lfc_std_age, y=-log10(p_std_age),
col=diff_std_age)) + geom_point(size=3) +
   geom_text_repel(aes(label = ifelse(diff_std_age, taxon, ""))) +
   theme_classic()+
   labs(x = "Lfc std. Age", y = "-Log10 (p std. Age)") +
   theme(axis.text.x = element_text(size = 16),  # Adjust the size as needed
         axis.text.y = element_text(size = 16))+
   theme(axis.title.x = element_text(size = 16),  # Adjust the size as needed
         axis.title.y = element_text(size = 16))+
   theme(axis.title.x = element_text(margin = margin(t = 13)))+
   theme(axis.title.y = element_text(margin = margin(r = 14)))+
   theme(text = element_text(family = "Arial"))
 volc_hapto_age
```

```
# Pseudo count sensitivity analysis

df_bci_asv = res_prim_immunity %>% dplyr::select(taxon, ends_with("two")) # create a dataframe
with values only for bci

df_age_asv = res_prim_immunity %>% dplyr::select(taxon, ends_with("age")) # create a dataframe
with values only for age

# Pseudo count sensitivity analysis for BCI
sens_asv_bci = tab_sens_immunity %>%
  transmute(taxon, sens_asv_bci = std_bci_two) %>%
  left_join(df_bci_asv, by = "taxon")

sens_asv_bci$diff_std_bci_asv = recode(sens_asv_bci$diff_std_bci_two * 1,
                                       `1` = "Significant",
                                       `0` = "Non-significant")

fig_sens_asv_bci = sens_asv_bci %>%
  ggplot(aes(x = taxon, y = sens_asv_bci, color = diff_std_bci_two)) +
  geom_point() +
  scale_color_brewer(palette = "Dark2", name = NULL) +
  labs(x = "ASVs", y = "Sensitivity Score") +
  theme_bw()

fig_sens_asv_bci

# Pseudo count sensitivity analysis for Age
sens_asv_age = tab_sens_immunity %>%
  transmute(taxon, sens_asv_age = std_age) %>%
  left_join(df_age_asv, by = "taxon")

sens_asv_bci$diff_std_age_asv = recode(sens_asv_age$diff_std_age * 1,
                                       `1` = "Significant",
                                       `0` = "Non-significant")

fig_sens_asv_age = sens_asv_age %>%
  ggplot(aes(x = taxon, y = sens_asv_age, color = diff_std_age)) +
  geom_point() +
  scale_color_brewer(palette = "Dark2", name = NULL) +
  labs(x = "ASVs", y = "Sensitivity Score") +
  theme_bw()
```

```
fig_sens_asv_age
```

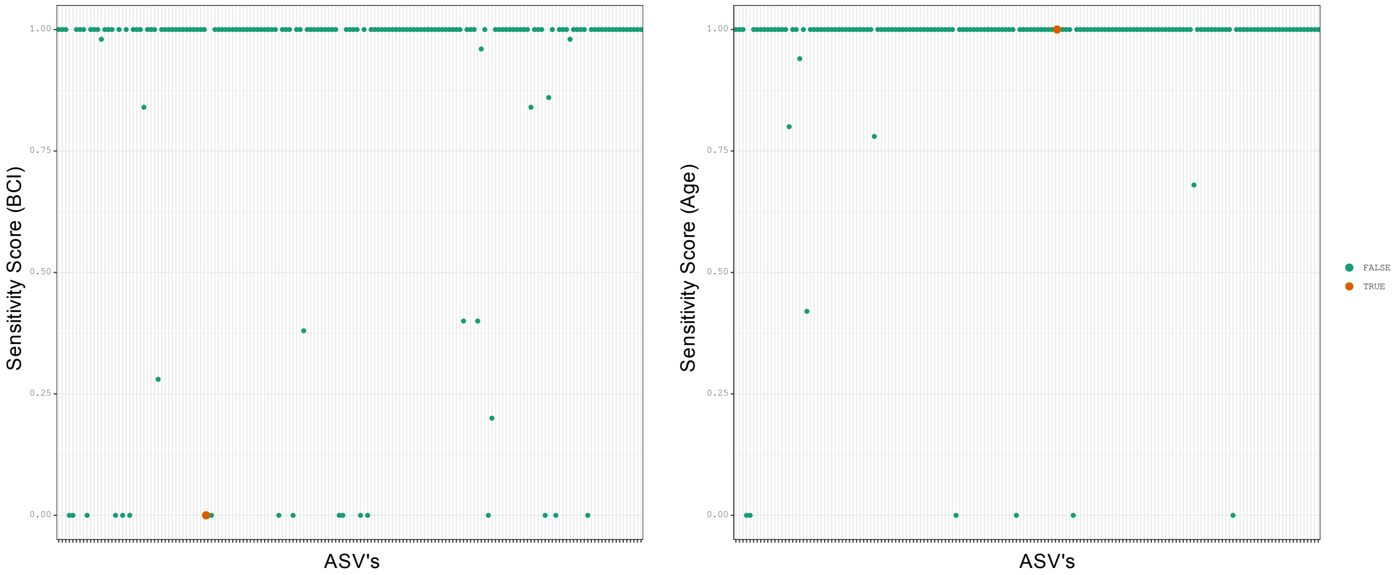


**For the co-variate of Age the deferentially abundant ASV has high sensitivity scores**

```
# Plot log fold changes with unit of BCI
df_fig_bci_asv = df_bci_asv %>%
  filter(diff_std_bci_two == TRUE) %>%
  arrange(desc(lfc_std_bci_two)) %>%
  mutate(direct = ifelse(lfc_std_bci_two > 0, "Positive LFC", "Negative LFC")) # prepare the
dataframe for plotting

df_fig_bci_asv$taxon = factor(df_fig_bci_asv$taxon, levels = df_fig_bci_asv$taxon)
df_fig_bci_asv$direct = factor(df_fig_bci_asv$direct,
                               levels = c("Positive LFC", "Negative LFC"))

fig_bci_asv = df_fig_bci_asv %>%
  ggplot(aes(x = taxon, y = lfc_std_bci_two, fill = direct)) +
  geom_bar(stat = "identity", width = 0.7, color = "black",
           position = position_dodge(width = 0.4)) +
  geom_errorbar(aes(ymin = lfc_std_bci_two - se_std_bci_two, ymax = lfc_std_bci_two +
se_std_bci_two),
                width = 0.2, position = position_dodge(0.05), color = "black") +
  labs(x = NULL, y = "Log fold change", title = "Log fold changes as one unit increase of BCI")
+
  scale_fill_discrete(name = NULL) +
  scale_color_discrete(name = NULL) +
  theme_bw() +
  theme(plot.title = element_text(hjust = 0.5),
        panel.grid.minor.y = element_blank(),
        axis.text.x = element_text(angle = 60, hjust = 1))
fig_bci_asv
```

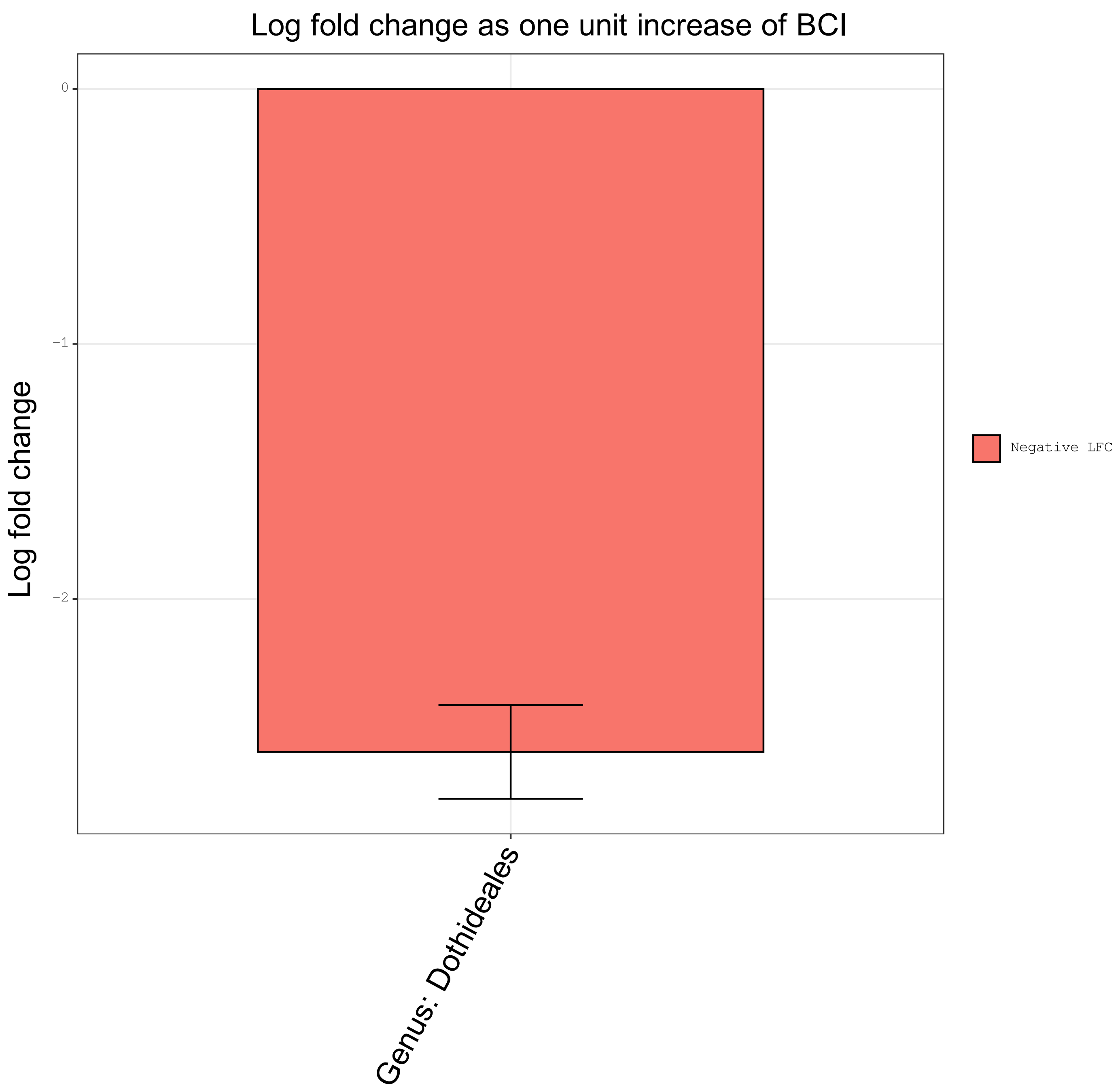

Log fold change as one unit increase of BCI
0
−1
−2
Log fold change
Genus: Dothideales
Negative LFC
