## Supplementary material for "The gut microbiota-immune-brain axis in a wild vertebrate: dynamic interactions and health impacts": f-CORT-assay-validation

### Cross reactions and inter and intra-assay variation

Three samples of different concentrations were run in duplicate on a total of 7 plates to assess inter-assay variation, which was on average 9.1 %. Furthermore, three pools of different hormone concentrations were used to determine intra-assay variance: corticosterone concentrations of one pool was were determined 16 times, the second pool for 14 times, the third pool for 10 times for one plate, resulting in an average intra-assay coefficient of 6.2 %. Finally, all samples were determined in triplicate, and determination of the sample was repeated if CV was larger than 10%.

The antibody showed the following cross-reactivities: corticosterone 100%, 11-deoxycorticosterone 15.8%, prednisolone 3.4%, 11 dehydrocorticosterone 2.9%, cortisol 2.5%, progesterone 1.4%, aldosterone 0.47%, 17α-hydroxyprogesterone 0.21%, 11-deoxycortisol 0.14%, androstendione 0.11% and all other tested steroids < 0.1%.

#### Table 1: Linearity

To assess linearity, we ran a serial dilution of three pooled samples in duplicate.

| | expected (pg/mg) | measured (pg/mg) | linearity (%) |
|---|---|---|---|
| **sample 1** | | | |
| neat | | 1066 | |
| 1:2 | 533 | 453.3 | 85.0 % |
| 1:4 | 266.6 | 245.7 | 92.2 % |
| 1:8 | 133.3 | 125.4 | 94.1 % |
| 1:16 | 66.7 | 62.8 | 94.2 % |
| 1:32 | 33.3 | 28.5 | 85.2 % |
| **sample 2** | | | |
| neat | | 716 | |
| 1:2 | 358 | 308.7 | 86.2 |
| 1:4 | 179 | 162.9 | 91.0 |
| 1:8 | 89.5 | 76.5 | 85.5 |
| 1:16 | 44.75 | 42.5 | 95.0 |
| **sample 3** | | | |
| neat | | 429.4 | |
| 1:2 | 214.7 | 220.9 | 102.9 |
| 1:4 | 107.4 | 116.7 | 108.7 |
| 1:8 | 53.7 | 59.8 | 111.4 |

| | expected (pg/mg) | measured (pg/mg) | linearity (%) |
|---|---|---|---|
| 1:16 | 26.8 | 26.3 | 98.0 |

### Table 2: Recovery rate

To assess recovery rate, three pooled samples of different concentrations were spiked with different amounts of corticosterone and measured in duplicate.

| | measured pg/ml | expected pg/ml | Recovery rate % |
|---|---|---|---|
| **sample 1** | | | |
| initial value | 1061 | | |
| + 64 pg/ml | 1074 | 1125 | 95.4 |
| + 160 pg/ml | 1120 | 1221 | 91.7 |
| + 400 pg/ml | 1433 | 1461 | 98.0 |
| **sample 2** | | | |
| initial value | 648 | | |
| + 64 pg/ml | 698 | 712 | 98.0 |
| + 160 pg/ml | 707 | 808 | 87.5 |
| + 400 pg/ml | 1130 | 1048 | 107.8 |
| **sample 3** | | | |
| initial value | 145 | | |
| + 64 pg/ml | 172 | 209 | 82.3 |
| + 160 pg/ml | 277 | 305 | 90.8 |
| + 400 pg/ml | 564 | 545 | 103.5 |
